# Fusogens for Axon Repair in Spinal Cord and Peripheral Nerve Injuries – Studies, Methods, and Mechanisms (systematic review with meta-analysis)

**DOI:** 10.64898/2026.03.20.712959

**Authors:** Michael Lebenstein-Gumovski, Yuriy Romanenko, Dmitry Kovalev, Tanzila Rasueva, Sergio Canavero, Andrey Zhirov, Alexander Talypov, Andrey Grin

**Affiliations:** Neurosurgery Department, Sklifosovsky Research and Clinical Institute for Emergency Medicine, Moscow, Russia; Laboratory of Biochemistry, Stavropol Scientific Research Anti-Plague Institute, Stavropol, Russia; GEMINI/HEAVEN International Collaborative Group, Turin, Italy; Pirogov Russian National Research Medical University, Moscow, Russia

**Keywords:** PEG, fusogen, axonal fusion, spinal cord injury, rapid nerve repair, spinal cord regeneration, neuroprotection, polyethylene glycol

## Abstract

**Introduction:** The exploration of alternative strategies for neural tissue regeneration and repair is giving rise to a novel paradigm in neurosurgery: fusogenic therapy. This approach promises rapid restoration of peripheral nerve and spinal cord function by circumventing Wallerian degeneration and eliminating the delay associated with axonal regrowth. Its potential stems from the capacity of fusogens to induce axonal fusion and achieve immediate membrane sealing, complemented by their pronounced neuroprotective properties. However, experimental data on fusogens and their effects are inconsistent, often contentious, and derived using heterogeneous methodologies.

**Methods:** We present the first comprehensive systematic review covering nearly four decades of research on fusogens for axonal membrane repair and 26 years of their experimental and clinical application in mammalian and human models for peripheral and central nervous system restoration. The review includes a meta-analysis of fusogen efficacy following traumatic spinal cord and peripheral nerve injuries.

**Results:** Conducted in accordance with the PRISMA 2020 flow protocol and PICO criteria, our analysis incorporates 86 sources, 20 of which were included in the meta-analysis.

**Discussion:** In summary, we have systematized the prevailing approaches and methods for fusogen application, delineated key contentious issues, and identified promising directions for the development of axonal fusion technology.

## 1 Introduction

Spinal cord injury (SCI) and peripheral nerve injury (PNI) frequently result in the severe and debilitating loss of motor and sensory function in affected patients, thereby constituting major global healthcare challenges. However, the efficacy of existing treatments remains limited, and there is a critical need for novel therapeutic strategies. While primary avenues for achieving functional recovery involve promoting axonal regeneration and the remyelination of damaged nerve fibers, strategies aimed at directly stimulating the fusion of cellular membranes present a particularly promising route for repairing severed axons. In this context, fusogens—substances that mediate membrane fusion and are well characterized in various fields of cell biology—are increasingly recognized for their notable capacity for activating regenerative processes in both the central and peripheral nervous systems (Neumann et al., 2019).

This review provides the first detailed synthesis of fusogen applications for the repair of spinal cord and peripheral nerve injuries spanning the past four decades. Beyond a systematic analysis of the literature, the review incorporates a meta-analysis of selected qualifying studies. Furthermore, we present a comprehensive critical discussion of the fusogenic techniques used for spinal cord repair, informed by both the analyzed data and our extensive experience in this field.

## 2 The Phenomenon of Cell Membrane Fusion

Membrane fusion is a fundamental cellular mechanism indispensable for a variety of physiological processes, including intracellular vesicular transport, intercellular communication, and tissue formation during embryonic development. Fusion events occur between gametes during fertilization, myoblasts during muscle formation, macrophages, and trophoblasts, and are also required for viral entry into cells (Chen E. H. et al., 2007; Oren-Suissa and Podbilewicz, 2007; Huppertz and Borges, 2008; Brodbeck and Anderson, 2009; Helming and Gordon, 2009; Pavlath, 2010).

Evolutionarily, membrane fusion is a tightly regulated process mediated by specialized protein machinery. Irrespective of the system’s complexity, fusion is primarily governed by molecular interactions at the membrane interface, driven by a combination of repulsive forces, hydration, hydrophobic attraction, and van der Waals forces (Yeagle, 1993).

A hypothetical capacity for spontaneous membrane fusion would lead to a catastrophic destabilization of cellular organization. The arbitrary fusion of cells and the indiscriminate merging of trillions of intracellular vesicles and organelles would disrupt essential compartmentalization and tissue integrity. However, even during prolonged and close contact, biological membranes do not spontaneously fuse. This stability is actively maintained by dense protein packing at membrane contact sites and by the presence of significant energy barriers that oppose membrane deformation, lipid mixing, and fusion pore expansion (Blobel et al., 1990). For fusion to become energetically favorable, membranes must overcome the repulsive forces generated by charged, hydrated phospholipid head groups while facilitating lipid mixing with minimal perturbation of their hydrophobic cores (Chen E. H. et al., 2007; Kozlov et al., 2010).

Conceptually, the fusion process can be divided into four principal stages, each constituting a complex sequence of molecular events (Zimmerberg and Chernomordik, 1999). The initial step involves membrane aggregation, where the participating membranes are brought within a few nanometers of each other. Subsequently, the two bilayers establish close contact (within a few Ångströms), a process that requires the partial dehydration of the membrane surfaces to overcome the strong repulsive forces generated by bound interfacial water at this range. Third, localized destabilization occurs at a single point between the two bilayers, inducing a highly restricted restructuring. Finally, as this point defect expands, the lipid components of the two bilayers intermix and diffuse away from the contact site. The internal aqueous contents of the compartments may also mix at this stage, depending on whether the outcome is hemifusion or complete fusion (Helm et al., 1992).

Certain exogenous substances can induce fusion by modulating the physicochemical and rheological properties of membranes. These agents, recently termed fusogens or fusogen-sealants, can target different sites within the membrane. For instance, following activation, most protein fusogens assemble into trimers anchored via transmembrane domains at one end, while exposing hydrophobic fusion peptides or amphipathic loops for insertion into the target membrane. At this stage, the interacting domains are located in separate membranes. Subsequent refolding of the fusogenic complex into a hairpin structure draws the membranes to within approximately 1 nm of each other. The insertion of hydrophobic peptides generates local membrane tension, thereby promoting deformation and lipid mixing. As the bilayers converge, the stored energy drives the formation of a hemifusion diaphragm, wherein only the proximal membrane leaflets merge while the distal leaflets remain intact. The expansion of this diaphragm and subsequent fusion of the distal leaflets completes the process, resulting in the opening of a fusion pore that allows the contents of the two compartments to mix. The expansion of this fusion pore represents the final energetic barrier that must be overcome to achieve irreversible membrane fusion (Chen E. H. et al., 2007; Kozlov et al., 2010).

The degeneration of the distal segment of an axon after injury has been a recognized phenomenon since Waller’s foundational work in 1850. In mammals, the balance at the lesion site favors degeneration over regeneration (Waller, 1850; Neumann et al., 2019). Interestingly, mammalian central nervous system neurons exhibit a capacity for axonal regeneration *in vitro* comparable to that of peripheral nerves (Kerschensteiner et al., 2005). However, *in vivo*, robust axonal regeneration in vertebrates is largely confined to amphibians and fish, and to a lesser extent, birds, owing to inhibitory cues present in the cellular microenvironment. While mammals retain a latent potential for spinal cord regeneration, it is not effectively executed. Instead, functional recovery is often achieved through mechanisms that are more energetically favorable than full axonal regeneration (Kerschensteiner et al., 2005; Neumann et al., 2019).

Endogenous fusogen molecules are known to facilitate the fusion of severed axonal membranes in lower vertebrates (Neumann et al., 2019). In invertebrates, this spontaneous axonal fusion enables rapid repair without the need for protracted axonal regrowth. For instance, in the nematode *Caenorhabditis elegans*, axon fragments can reconnect via axonal fusion, whereby the proximal and distal stumps merge minutes to hours after injury, re-establishing axonal continuity. However, this spontaneous repair mechanism does not occur in mammals (Neumann et al., 2019).

Endogenous molecules that mediate fusion are known as fusogens. An example of such a molecule is epithelial fusion factor-1 (EFF-1), whose activity in invertebrates is regulated by the GTPase RAB-5 (Villarroel-Campos, et al., 2016). The impairment of RAB-5 secretion leads to the accumulation of EFF-1. Provided that the fragments remain in close apposition after transection, the accumulated EFF-1 molecules then draw together the ruptured axonal membranes (Kerschensteiner et al., 2005). In mammals, annexin A1/A2, which participates in membrane repair, is thought to serve as a functional analog of EFF-1 (Neumann et al., 2019). The identification of additional molecules involved in axonal fusion has revealed intriguing parallels between this process and axonal apoptosis, with these ostensibly opposing mechanisms sharing common physiological pathways. During non-injury-induced apoptosis, the phospholipid phosphatidylserine is externalized on the axonal plasmalemma, marking the damaged membrane for recognition by phagocytes. The asymmetric localization of phosphatidylserine generates a high anionic charge, which induces membrane curvature and leads to the recruitment of lipid-binding proteins. Identical processes are observed after axonal injury: phosphatidylserine molecules on the distal axon segment act as markers for the proximal segment’s plasmalemma, inducing membrane protrusion and the formation of a “bridge”. Upon contact, membrane fluidity increases at the bridge site, culminating in plasmalemma fusion and the restoration of axonal integrity (Neumann et al., 2015).

The phenomenon of cell fusion in the nervous system is also observed during aging, during viral infections, and following the transplantation of stem or progenitor cells (Giordano-Santini et al., 2016). Two endogenous fusogens—syncytin-1 and syncytin-2—have been identified in humans, though their expression is normally restricted to the placenta (Blaise et al., 2003). Notably, syncytin-1 is expressed in the brain of patients with multiple sclerosis (Mameli et al., 2015). Confirming the presence of such fusogens in humans could provide key insights into the potential for axonal fusion and repair mechanisms in mammals.

The effect of proximal axon degeneration, the expulsion of Schwann cell bands, and the formation of a growth cone on the distal axon—all of this formed the basis of the classical neurorrhaphy method. Nerve suturing creates conditions for axon degeneration and growth (Figure 1). However, the time for axon growth is limited by a rate of 1-4 mm per day, the onset of muscle atrophy, and connective tissue degeneration of the nerve during growth, which can last for months. The principles of neurorrhaphy are completely inapplicable to SCI, due to the different microenvironment, cellular responses, and immune cascades. The search for alternative methods of nerve tissue regeneration leads to both cell technologies and the use of biosynthetic substances as a fusogens.

**Figure 1.**
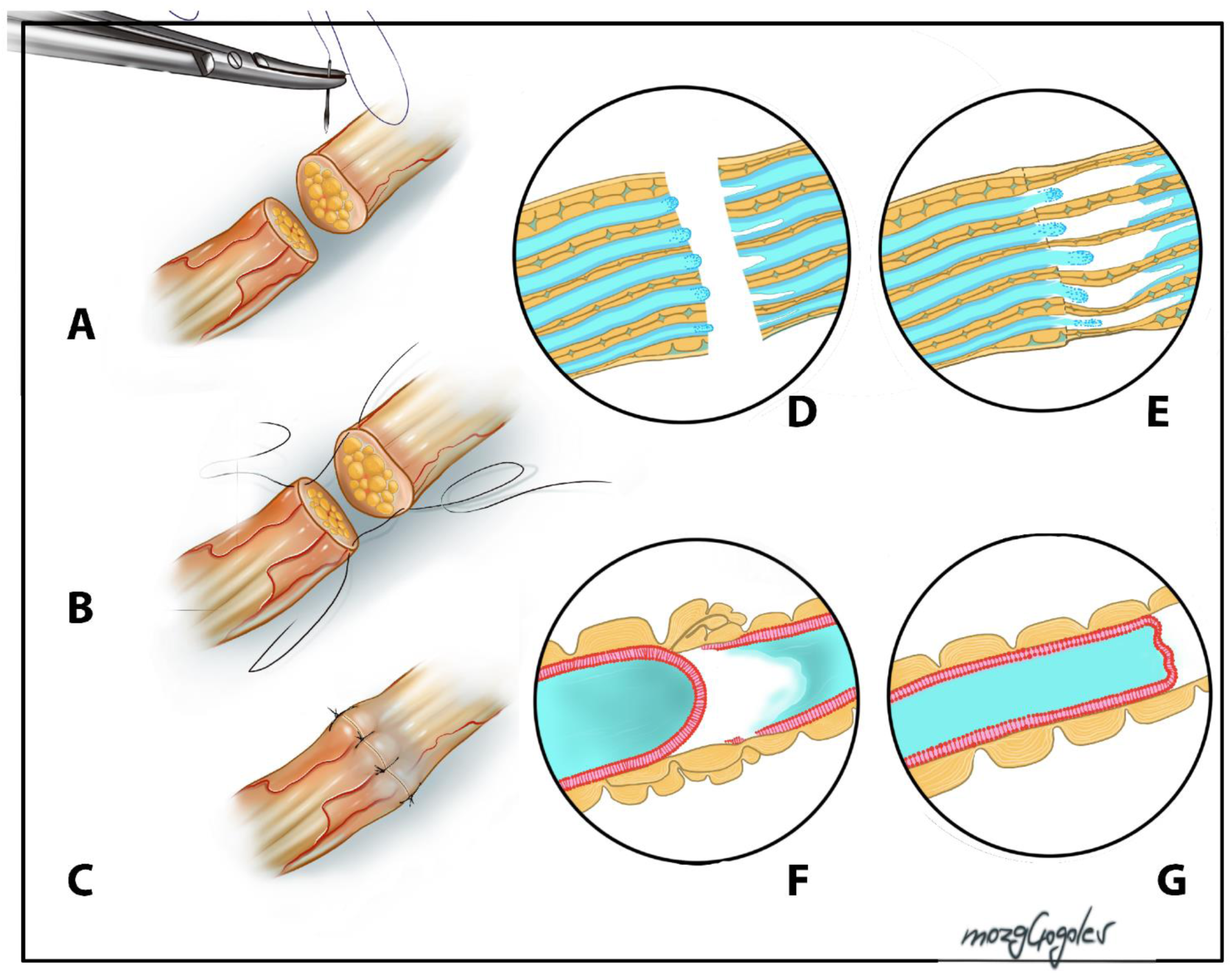
Classic neurorrhaphy technique. **A** - preparation of the nerve ends; **B** - suturing and alignment of the nerve ends; **C** - final appearance of the sutured nerve; **D** - schematic representation of axons at the proximal (left) and distal (right) ends of the nerve. Growth bulbs are forming at the proximal end, while axonal degeneration begins at the distal end, preserving Schwann cells; **E** - schematic representation of axons in the sutured nerve. Continued degeneration of the distal portions of axons on the right and gradual growth of proximal axons through the anastomosis site; **F** and **G** - schematic representation of an individual axon at the site of injury. Growth bulb formation, degeneration of the distal axon, and growth of the proximal portion.

## 3 Exogenous Fusogens

It has been reported that exogenous fusogens, such as polyethylene glycol (PEG), chitosan, and poloxamer-188, among others, can also initiate a reparative process, one that avoids the substantial energetic costs of apoptosis and axonal regrowth associated with classical regeneration (Cho and Borgens, 2012). PEG is a linear polymer known for its high stability, biocompatibility, and water solubility. It is widely employed in clinical and pharmaceutical applications, ranging from an oral laxative to a key component in PEGylated therapeutics. PEG has received FDA approval as an additive for organ preservation prior to transplantation, where it functions to mitigate cold ischemia and reperfusion injury (Pasut et al., 2016). It does not bioaccumulate and can traverse the blood-brain and blood-spinal cord barriers. Furthermore, investigations into the long-term systemic (intravenous) administration of high-dose, low-molecular-weight PEG in dogs have uncovered no evidence of toxicity (Li et al., 2011).

PEG has been employed as a cell-fusion agent for over 50 years, enabling the creation of giant cells from smaller counterparts and thereby facilitating electrophysiological studies and membrane manipulations (Kao and Michayluk, 1974; Pontecorvo, 1975; Greenfield, 2018; Chang et al., 2023). Furthermore, PEG-induced cell fusion also promotes genetic material exchange and is used for hybridoma formation in monoclonal antibody production. It also serves as a model for studying vesicular fusion, a cornerstone of cellular biology (Lentz, 1994; Lee J. and Lentz, 1997). Nonetheless, the precise molecular mechanisms underpinning PEG-mediated membrane fusion are incompletely understood and remain an area of active investigation.

PEG-mediated cellular fusion is hypothesized to occur in stages, commencing with contact between adjacent membranes in the presence of PEG, a prerequisite for full cellular fusion and cytoplasmic mixing (Ahkong et al., 1987). At the membrane level, PEG exerts an acute dehydrating effect on the apposed plasmalemmae, which promotes the intermixing of glycolipid and protein components. These components dissolve into one another first in the outer membrane leaflet, and then in the inner leaflet (Lee J. and Lentz, 1997). Subsequent rehydration appears to spontaneously induce structural self-assembly within the aqueous membrane environment (Shi and Borgens, 1999). PEG reduces the surface potential of phospholipids in monolayers and binds to the cell membrane surface, interacting with lipid head groups. This interaction compromises membrane stability and promotes bilayer disassembly. Thus, PEG functions not only as a membrane modifier but also as an effective destabilizing agent, complementing its aggregative properties (Abdou and Henderson, 2019). It has been proposed that when applied to injured nerve cells, PEG creates a linkage between damaged membranes and their underlying axoplasm via interaction with the Epsin N-terminal homology domains of membrane proteins, thereby preserving the integrity of the ruptured membrane. Among PEG polymers, PEG-600, PEG-400, and PEG-200 exhibit the strongest interaction with lipid-binding domains (Rad et al., 2015).

The use of PEG for repairing severed axons was first described in 1986 (Bittner et al., 1986). Studies using animal models demonstrated that when PEG is applied early following surgical intervention, it effectively fuses transected peripheral nerves, thereby improving functional outcomes. Further research has demonstrated that PEG has considerable potential for use in diverse SCI repair methodologies (Lebenstein-Gumovski et al., 2023b; Shen T. et al., 2024; Zhang et al., 2024a).

PEG possesses marked neuroprotective properties, which are attributable to its capacity for membrane sealing and fusion, and can also stimulate neoangiogenesis (Robinson and Madison, 2016; Kong et al., 2017; You et al., 2023). This pro-angiogenic characteristic is critically important in the treatment of nervous system injuries and revascularizing affected areas. PEG has also been reported to exert angioprotective effects in spinal cord trauma (Kong et al., 2017; Lee J. et al., 2023).

PEG can reduce neuronal membrane tension and enhance membrane fluidity, thus enabling effective membrane sealing even at low temperatures (Nehrt et al., 2010; Shi, 2013). High-molecular-weight PEG is used in organ preservation for its protective and restorative effects, including the prevention of apoptosis (Goddard et al., 1999). For instance, PEG-3500 at a dose of 10 mg/kg has been shown to reduce the levels of biomarkers of liver injury, attenuate mitochondrial depolarization, decrease caspase-3 and caspase-9 activity, and prevent the decline in the F-actin/G-actin ratio (Chiang et al., 2009). Furthermore, a 5-minute topical application of PEG at the injury site was reported to be sufficient to seal neuronal membranes and reduce biomarkers of oxidative stress (decreased lipid peroxidation and protein carbonyl content, alongside increased glutathione levels) in and around the lesion site following acute SCI in guinea pigs.

PEG exerts its neuroprotective effects through several mechanisms. Its primary mechanism of action involves the facilitation of the restoration of damaged membrane integrity, thereby mitigating primary injury. Concurrently, PEG reduces the severity of secondary injury, mainly by diminishing the production of reactive oxygen species (ROS). *In vitro* studies have demonstrated that PEG significantly suppresses superoxide formation within the first hour post-injury (Shi, 2013). Moreover, PEG application markedly reduces the levels of lipid peroxidation, a key component of the secondary injury cascade in spinal cord trauma pathogenesis. Nevertheless, PEG does not exhibit direct antioxidant activity; for instance, it neither neutralizes free radicals nor inhibits xanthine oxidase, a major enzymatic source of superoxide. Instead, the mechanism underlying its anti-oxidative stress activity likely involves the restoration of membrane integrity and the lowering of cellular calcium overload, which interrupts the ensuing cycle of excitotoxicity (Luo et al., 2002, 2004; Luo and Shi, 2004).

PEG also attenuates Ca²⁺-induced functional impairments, prevents cellular swelling, and inhibits glutathione oxidation (Luo et al., 2004; Chen H. et al., 2009). By sealing membrane breaches that would otherwise lead to the release of glutamate, aspartate, and intracellular calcium—triggering excitotoxicity—PEG helps halt the cascade of toxic secondary injury events.

*In vitro* models have established that the observed reduction in endogenous toxicity occurs only when PEG penetrates the cytosol of cells whose membranes are only partially destroyed, highlighting the importance of the fusogen’s interaction with cellular structures (Liu-Snyder et al., 2007). Traumatic SCI results in neuronal membrane rupture and the disruption of ionic balance, leading to rapid organelle damage and cell death. While the primary action of PEG involves the immediate sealing of membrane ruptures, a fraction of PEG molecules also enters the cell through these membrane discontinuities and interacts with mitochondria, preventing the formation of mitochondrial permeability transition pores. This, in turn, suppresses or reduces mitochondrial swelling, helps maintain mitochondrial membrane potential, and inhibits cytochrome c release and the subsequent depletion of antioxidant systems. PEG was also observed to slow the depletion of intracellular glutathione, a key mitochondrial antioxidant, likely due to its stabilizing effect on the mitochondrial membrane. These effects not only reduce oxidative stress but also contribute to improved mitochondrial function (Luo and Shi, 2007). These observations imply that PEG restores mechanically damaged cells through at least two mechanisms: reestablishing damaged plasma membrane integrity and directly protecting mitochondria (Shi, 2013). The *in vivo* administration of PEG has also been demonstrated to suppress apoptosis. Its local application immediately after injury results in a reduced number of apoptotic cells and decreased caspase-3 activity (Luo and Shi, 2007).

The neuroprotective properties of PEG are attributable to its ability to seal membranes, its influence on intracellular organelles, particularly mitochondria, preventing apoptosis, and preserving cytoskeletal elements (Liu-Snyder et al., 2007; Baptiste et al., 2009; Chen H. et al., 2009).

Injectable formulations of PEG have also been explored. For example, PEG combined with magnesium chloride was shown to improve ladder test performance in rats with cervical spinal cord hemiconcussion within just 2 hours of administration (Lee J.H.T et al., 2010). Cho and Borgens (2012) tested a modified PEG—monomethoxy-polyethylene glycol-poly-D,L-lactic acid (mPEG(2000)-PDLLA)—featuring a micellar structure with a hydrophobic core and a mean diameter of 60 nm. They noted that such micelles persist in the bloodstream for a much longer duration, readily cross the blood-brain barrier, and demonstrate efficacy at low doses comparable to that of pure PEG at higher, subtoxic concentrations. Moreover, the micelles exerted a reparative effect on myelinated spinal cord axons (Gao et al., 2014). In the spinal trauma model employed by Cho and Borgens (2012), this formulation achieved significant recovery of spinal cord function and rapid regression of hemiplegia symptoms (within 2–3 weeks).

Chitosan is another promising substance exhibiting neuroprotective, fusogenic, and scaffolding properties. It is widely recognized for its biocompatible (Kim I. Y. et al., 2008) and capacity for degradation into non-toxic and non-immunogenic byproducts (Muzzarelli, 1993). Chitosan has been extensively investigated for numerous biomedical applications, including tissue culture scaffolds (Kim I. Y. et al., 2008), drug and gene delivery vectors (Bowman and Leong, 2006), and wound healing (Kozen et al., 2008). Its adhesive properties are likely due to strong intermolecular interactions, such as hydrogen bonding and ionic attraction between the polycationic chitosan and anionic moieties of collagen, which is ubiquitous in mammalian tissues (Taravel and Domard, 1993).

For instance, the administration of chitosan combined with the neurotrophic factor NT3 into a 1-cm hemisection and resection site in the thoracic spinal cord of adult rhesus macaques resulted in axonal regeneration within the injury area. Corticospinal tract axons successfully grew through the 1-cm gap and reached the distal spinal cord, accompanied by the recovery of both motor and sensory functions (Rao J. S. et al., 2018). Comparable outcomes were obtained in rodent models (Zhao C. et al., 2022). Like PEG, chitosan is a fusogen that functions by promoting cell aggregation and inducing phospholipid fusion through electrostatic interactions between its cationic amino group and the anionic phosphate groups of cell membranes (Abdou and Henderson, 2019). Chitosan can modulate membrane permeability and disrupt negativelycharged membranes (Mertins and Dimova, 2013). Additionally, it can form large phospholipid aggregates, inducing the fusion of small dipalmitoylphosphatidylcholine bilayers, a major component of the plasma membrane (Zuo et al., 2006; Pavinatto et al., 2007).

Chitosan and its derivatives (chitooligosaccharides, COS) also demonstrate neuroprotective effects mediated through the stabilization of damaged membranes, thereby inhibiting ROS production and lipid peroxidation (Cho et al., 2010). However, unlike PEG, chitosan also exhibits anti-inflammatory properties. Chitosan was reported to suppress the production of tumor necrosis factor-alpha (TNF-α) and interleukin-6 (IL-6), inhibit NF-κB activation, and block apoptosis both via the modulation of p53 and by protecting against excitotoxicity (Pangestuti and Kim S. K., 2010).

The combined application of chitosan and PEG in the treatment of nervous tissue injuries has demonstrated superior efficacy. In a complete transection model of the spinal cord in pigs, this combination elicited significant recovery of motor and sensory functions between 9 and 60 days of treatment (Lebenstein-Gumovski et al., 2023a; Lebenstein-Gumovski et al., 2023b; Lebenstein-Gumovski et al., 2023c). Mixtures of photocrosslinkable chitosan modified with 4-azidobenzoic acid (Az-C) and PEG form a semi-interpenetrating network (semi-IPN), where PEG permeates the Az-C structure, enhancing its mechanical properties. Nerves anastomosed with Az-C/PEG gel demonstrate greater strength than those connected with fibrin glue, display good compatibility with nerve tissues and cells, and provide sustained PEG release, ensuring continuous delivery of the polymer to the injury site (Amoozgar et al., 2012).

## 4 Hemostatic Properties of Fusogens

It is postulated that PEG can arrest minor capillary bleeding in both bone and soft tissues. For example, capillary bleeding was halted within 2 minutes of PEG application to the spinal cord (J. P. Toombs and R. B. Borgens, unpublished observations). PEG likely seals endothelial injuries, which may also contribute to mitigating progressive spinal stroke associated with severe compression (Borgens and Shi, 2000; Borgens and Bohnert, 2001). Furthermore, PEG is known to form covalent cross-links with tissue proteins, thereby creating physical barriers that prevent bleeding. PEG-based hydrogels possess good hydrophilic properties and can concentrate platelets and clotting factors at the injury site (Le Huec et al., 2022; Guo et al., 2023).

Various hemostatic materials have been developed based on chitosan. The mechanisms underlying chitosan-mediated hemostasis likely involve its ability to increase Ca^2+^ ion concentration in platelets, thus promoting their aggregation and mutual adhesion. Platelet activation exposes numerous negatively charged phosphatidylserine units on the platelet surface, which electrostatically interact with the positively charged chitosan, resulting in increased platelet viscosity. Furthermore, through its positive surface charge, chitosan interacts with receptors containing sialic acid residues (negative charge) on the surface of erythrocytes, promoting their adhesion. Concurrently, the contact activation system is triggered through interaction with blood clotting coagulation factors, which amplifies the coagulation cascade (Rao S. B. and Sharma, 1997; Hu et al., 2018; Guo et al., 2023).

These hemostatic properties of fusogens are of particular importance when they are applied locally to the spinal cord, where they act to prevent hemorrhages, hematoma formation, and hemorrhagic infiltration of nervous tissue.

The properties of fusogens described above have been repeatedly confirmed in numerous studies. However, considerable methodological variability, alongside the diverse range of fusogenic agents employed for the treatment of peripheral nerve and spinal cord injuries, has generated several inconsistencies and unresolved questions. To address this, we subsequently provide a comprehensive analysis of available studies and discuss key considerations regarding the action of fusogens and their potential for clinical application.

## 5 Methods

A systematic review of both preclinical and clinical studies investigating fusogen-based treatments for spinal cord or peripheral nerve injuries was conducted. Data were extracted from articles meeting predefined inclusion criteria. The extracted data underwent both qualitative synthesis and quantitative meta-analysis.

### 5.1 Search Strategy

This systematic review adhered to Preferred Reporting Items for Systematic Reviews and Meta-Analyses (PRISMA) 2020 guidelines (Figure 2). A comprehensive search of the MEDLINE/PubMed and ScienceDirect databases was performed in June 2025, encompassing all available records from database inception. The following search keywords were applied: (“Spinal cord” OR “Peripheral nerve”) AND (“Polyethylene glycol” OR “PEG” OR “Fusogen” OR “Chitosan” OR “Axonal Fusion” OR “PEG-fusion”).

**Figure 2.**
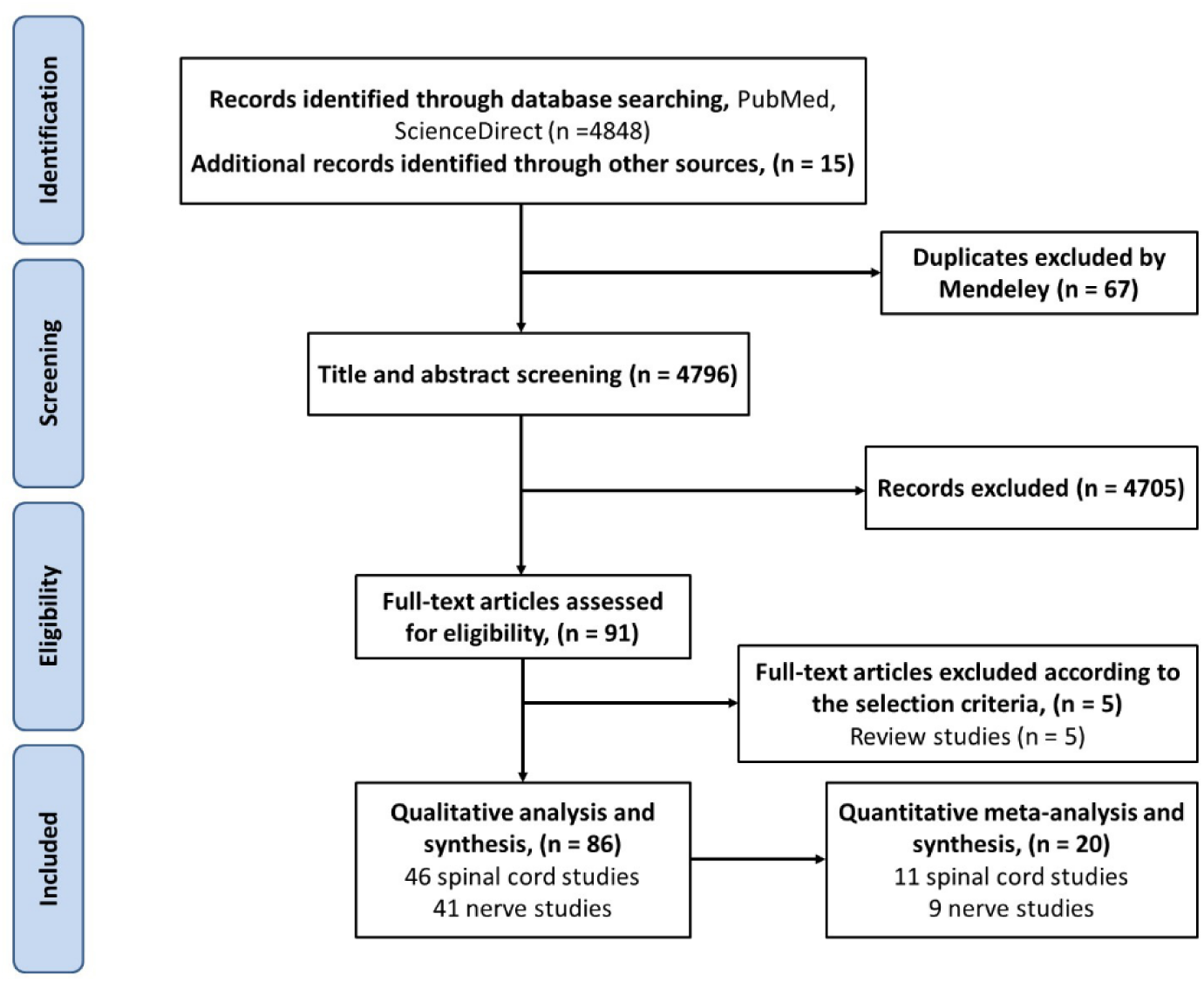
PRISMA flow diagram of the literature search for this systematic review. To ensure a complete dataset, the primary search was supplemented by a targeted author search focusing on key contributors to the field (R.B. Borgens and G.D. Bittner). Furthermore, the reference lists of all included articles were screened to identify any additional relevant publications. This multi-faceted approach mitigated the risk of omitting studies due to indexing constraints or terminological variance.

### 5.2 Selection Criteria and PICO Definition

Studies were deemed eligible if they reported on the use of fusogens (e.g., PEG, chitosan) to treat spinal cord or peripheral nerve injuries in mammalian models or clinical studies. The review adhered to the PICO framework:

- **P** (Problem): Mammals or humans with spinal cord or peripheral nerve injuries.
- **I** (Intervention): Application of fusogenic agents.
- **C** (Comparison): Treatment groups compared to negative controls receiving standard care, placebo, or no intervention.
- **O** (Outcome): Primary outcomes included improved motor function and the restoration of neural signal conduction and structural integrity.

### 5.3 Study Selection Process

Following the compilation of an initial article library, duplicates were removed using Mendeley. Three reviewers independently screened the remaining titles and abstracts. Included studies featured experimental groups treated with fusogens post-injury. Non-English publications were excluded unless a complete English translation was available.

Conference abstracts, videos, inaccessible articles, and previously published narrative or systematic reviews were excluded. The relevance of each study was first assessed based on its title and abstract; if inconclusive, the full text was evaluated. Disagreements regarding eligibility were resolved through consensus among the screening authors, with arbitration undertaken by a senior reviewer when necessary.

### 5.4 Participants

Eligible participants included mammalian subjects of any species, age, sex, or strain that sustained a SCI or PNI via any experimental model. Given the scarcity of human trials, clinical studies were also included.

### 5.5 Interventions

Included interventions comprised the use of any fusogen in liquid form, regardless of molecular weight (MW), compared against a placebo or standard care control group. Modes of administration included topical application at the injury site; intravenous, intraperitoneal, subcutaneous, or subdural injection; and combination therapies. Studies using fusogens in conjunction with other treatments (e.g., PEG with magnesium salts) were included, provided they met all other criteria.

Studies were excluded if they used fusogens in solid formulations (e.g., scaffolds, conduits, nanoparticles, microspheres, films, matrices, polyplexes) or as drug delivery vehicles. Research focusing on injuries with large segmental defects repaired with artificial grafts, rather than auto- or allografts, was also excluded, as this model precludes the axonal fusion mechanism targeted in this review.

### 5.6 Data Extraction and Qualitative Synthesis

The following data were extracted from eligible studies: lead authors and publication year; fusogen type (including MW, concentration, and administration route); model system (species, strain); injury paradigm; maximum follow-up duration; functional outcomes (assessed via validated neurological scales or electrophysiological measures); and histological results. The therapeutic benefits of various fusogen treatments were qualitatively assessed by comparing outcomes with control groups and other experimental cohorts.

### 5.7 Quantitative Data Extraction and Statistical Analysis

Descriptive charts summarizing preclinical data (e.g., studies by species, injury model, lesion location, and PEG type) were generated in Microsoft Excel.

For meta-analysis, the following data were extracted from preclinical studies: first author and year; animal model; sample sizes for experimental and control groups; injury details; fusogen type; and functional scores (e.g., Basso-Beattie-Bresnahan (BBB), Sciatic Functional Index (SFI)) from weeks 1 to 6. Only data from fusogen-only groups and negative controls were included for studies with multiple arms. Numerical data not available in the text were extracted from figures using GetData Graph Digitizer 2.26 and linearly interpolated in Microsoft Excel. Raw data were obtained from the author (Lebenstein-Gumovski et al., 2023a; Lebenstein-Gumovski et al., 2023b) for precise mean ± standard deviation (SD) calculation.

Data are presented as mean ± SD. When only the standard error of the mean was reported, SD was calculated as standard error of the mean (SEM) × √n. To account for the use of different functional assessment scales across species (e.g., modified Basso, Beattie, Bresnahan scale (mBBB) for dogs), treatment efficacy was quantified using the standardized mean difference (SMD, Hedges’ g).

Pairwise meta-analysis and subgroup analyses were conducted in Review Manager v.5.4.1 (Cochrane Collaboration). A random-effects model was employed to estimate the pooled effect size, presented as SMD with 95% confidence intervals (CIs). Heterogeneity was assessed using *χ*² and *I*² statistics. Statistical significance was set at *p*< 0.05 (two-tailed). Temporal effect size trends were visualized using GraphPadPrism 10.5.0.

## 6 Results

The systematic search yielded 71 studies meeting the inclusion criteria. A further 15 relevant studies were identified through manual screening of reference lists. Thus, from an initial 4,796 records, 86 studies were selected for final analysis, with 46 pertaining to SCI and 41 to PNI. One study (Marzullo et al., 2002) was applicable to both categories and was consequently included in both totals (Tables 1 and 2).

**Table 1.** Use in Spinal Cord Injury.

| № | Authors/Year | Title | Fusogen | Subjects | Injury model | Max. duration | Scales | Results | Histology |
| --- | --- | --- | --- | --- | --- | --- | --- | --- | --- |
| 1 | Shi R. and Borgens R.B., 1999 | Acute repair of crushed guinea pig spinal cord by polyethylene glycol | PEG-1800 50% w/w in distilled water, topical application for 2 min + oxygenated Krebs solution + 4-aminopyridine for 15 min | Guinea pigs.<br>Exp: $n = 10$<br>Ctrl: $n = 10$<br>Total: $n = 20$ | Compression of extracted SC at the mid-thoracic level, *ex vivo* | 1 hour | CAP | Exp.: 100% recovery; earliest recorded CAP within 1–2 min after PEG application.<br>Ctrl: 0% | Exp.: A greater number of preserved myelinated axons myelinated axons. Less vacuolization, myelin dispersion, and degenerative changes. Electron microscopy showed preserved membrane structure and partial fusion of severed axons, consistent with the effects of PEG on membrane fusion |
| 2 | Shi R. et al., 1999 | Functional reconnection of severed mammalian spinal cord axons with polyethylene glycol | PEG-1400, PEG-2000, PEG-3500, and PEG-1800 50% w/w in distilled water were used, but data are presented only for PEG-1800, topical application for 2 min + oxygenated Krebs solution | Guinea pigs.<br>Exp.: $n = 20$;<br>Ctrl: $n = 5$;<br>Total: $n = 25$ | Compression of extracted SC at the mid-thoracic level, *ex vivo* | 1 hour | CAP | Recovery of evoked potentials was often observed within 5-15 min after PEG application and lasted up to 60-80 min, after which physiological recordings ceased. Recovered CAPs exhibited approximately 5% of the peak amplitude of the original pre-transection CAPs.<br>AP amplitude: $4.61 \pm 2.83\%$;<br>peak latency: $114.5 \pm 8.71\%$; $\frac{1}{2}$ height duration: $85.5 \pm 6.45\%$ | Exp.: Intracellular diffusion of two different fluorescent dyes across the injury site was observed. Axons filled with different dyes were examined independently. In Ctrl, dye introduced into axons of one segment was never observed in axons of the opposite SC segment |
| 3 | Borgens R.B., and Shi R., 2000 | Immediate recovery from spinal cord injury through molecular repair of nerve membranes with polyethylene glycol | PEG-400 or PEG-1800 50% w/w in distilled water (data combined), topical application for 2 min + Krebs solution | Guinea pigs.<br>15-min compression group:<br>Exp.: $n = 14$;<br>Ctrl: $n = 11$.<br>8 hours post-injury group:<br>Exp.: $n = 11$;<br>Ctrl: $n = 11$;<br>Total: $n = 47$ | Compression of SC at the mid-thoracic level, *in vivo* | First group - 4 days;<br>Second group - 1 month | CTM | 15 min: Exp.: 10/14; Ctrl: 2/11.<br>8 hours: Exp.: 10/11; Ctrl: 1/11.<br>No statistically significant difference between the timing of PEG application | No |
| 4 | Shi R. and Borgens R.B., 2000 | Anatomical repair of nerve membranes in crushed mammalian spinal cord with polyethylene glycol | PEG-1800 50% w/w in distilled water, topical application for 2 min + oxygenated Krebs solution + 4-aminopyridine for 15 min | Guinea pigs.<br>Exp.: $n = 11$;<br>Ctrl: $n = 13$;<br>Total: $n = 24$ | Compression of extracted SC at the mid-thoracic level, *ex vivo* | 1 hour | CAP | Exp.: After 15 min ~65%;<br>Ctrl after 15 min ~25% of baseline values | No |
| 5 | Borgens R.B., and Bohnert D., 2001 | Rapid recovery from spinal cord injury after subcutaneously administered polyethylene glycol | PEG-2000 30% w/w in sterile lactated Ringers solution, subcutaneous, 1 mL single dose.<br>For tracing: FITC-PEG-2500 (PEG-2000, FITC-labeled) – FITC-PEG 50% w/w in sterile lactated Ringers solution, topical or 30% subcutaneous 1 mL single dose or 30% IV or intraperitoneal | Guinea pigs.<br>Exp.: $n = 10$;<br>Ctrl: $n = 10$;<br>Total: $n = 20$ | SC compression in the thoracic region, *in vivo* | 4 weeks | CTM, SEP | CTM: recovery<br>Exp.: 7/10;<br>Ctrl: 0/10.<br>SSEP: recovery<br>Exp.: 10/10;<br>Ctrl: 0/10 | FITC-labeled PEG accumulated in the area of hemorrhagic contusion but did not penetrate into unaffected areas of the spinal cord |
| 6 | Borgens R.B. et al., 2002 | Behavioral recovery from spinal cord injury following delayed application of polyethylene glycol | PEG-1800 50% w/v in distilled water, topical application for 2 min at 7 hours post-injury + isotonic Krebs solution | Guinea pigs.<br>Exp.: $n = 15$;<br>Ctrl: $n = 13$;<br>Total: $n = 28$ | SC compression at level Th10-Th11, *in vivo* | 1 month | SSEP, CTM | SEEP:<br>Exp.: 100%;<br>Ctrl: 0%.<br>CTM: Recovery in the Exp. group within the first 24 h was 73%. After one month, CTM recovery was observed in 14 Exp. animals and spontaneous recovery was detected in 3 Ctrl animals | No |
| 7 | Duerstock B.S. and Borgens R.B., 2002 | Three-dimensional morphometry of spinal cord injury following polyethylene glycol treatment | PEG-1800 50% w/v in distilled water, topical for 2 min immediately or at 7 hours post-injury + isotonic Krebs solution | Guinea pigs.<br>PEG immediate: $n = 5$;<br>PEG at 7 h: $n = 7$;<br>Saline: $n = 5$;<br>Total: $n = 17$ | SC compression at level Th10-Th11, *in vivo* | 1 month | None | Statistically significant changes were observed in the PEG-treated groups, including an increase in the volume of intact SC parenchyma, a decrease in the number of cystic cavities, and overall lesions that were smaller and less diffuse within the injured area of the SC compared with the Ctrl. A significant inverse relationship was found between the volume of intact parenchyma and cyst volume in | PEG application increased the amount of preserved parenchyma in the injured SC and significantly reduced the extent of cavitation and cyst formation. The average volume of normal-appearing parenchyma post-injury in Saline was 24.4%, while 35.8% of neural tissue survived in the immediate PEG group and 49.7% in the delayed PEG group. On average, 46.9% of the total volume of SC segments in Saline consisted of cysts compared to 32.4% and 21.7% in segments treated with |
|  |  |  |  |  |  |  |  | the delayed PEG treatment group (P = 0.03) | PEG immediately and delayed, respectively |
| 8 | Luo J. et al., 2002 | Polyethylene glycol immediately repairs neuronal membranes and inhibits free radical production after acute spinal cord injury | PEG-2000 50% w/w in distilled water, topical for 3 min + oxygenated Krebs solution | Guinea pigs. Number of animals not specified | Compression of extracted SC at the mid-thoracic level, *ex vivo* | 1 hour | None | The percentage reduction in horseradish peroxidase permeability induced by PEG exposure was 33% at 1 min, 46% at 15 min, and 46% at 60 min. Similarly, lactate dehydrogenase accumulation in extracellular fluids decreased by 45% at 60 min post-compression and PEG treatment. PEG application reduced the level of lipid peroxidation to $14.6 \pm 1.2$ nmol/100 mg, a 24.3% decrease. PEG at two concentrations (50% and 5%) had almost no effect on free radical scavenging. PEG at two concentrations (50% and 5%) had a minor effect on xanthine oxidase activity (4.5% and 0.05% inhibition, respectively) | PEG exposure resulted in a 47% and 48% reduction in ethidium bromide fluorescence at 15 and 60 min post-compression, respectively, compared to Ctrl. Superoxide production, indicated by oxidized erythrosine (ethidium) fluorescence, increased by 118% (218% of control) at 1 hour post-injury. PEG application significantly reduced superoxide production at 1 hour post-injury, as indicated by a decrease in fluorescence from 218% to 178% compared to the intact group |
| 9 | Marzullo T.C. et al., 2002 | Cooling enhances in vitro survival and fusion-repair of severed axons taken from the peripheral and central nervous systems of rats | PEG-2000 50% w/w in distilled water, topical for 3 min. + oxygenated Krebs solution with $\text{Ca}^{2+}$ + hypothermia | Rats, Sprague-Dawley. Number of animals not specified | Crush/compression of SC at the mid-thoracic level, *ex vivo* | 48 hours | CAP | SC stored *in vitro* at 6–9°C prior to crush successfully underwent PEG-mediated fusion for up to 1.5 days compared to 3 hours at 37-38°C | No |
| 10 | Borgens R.B.<br>et al., 2004 | Subcutaneous tri-block copolymer produces recovery from spinal cord injury | P188 subcutaneously, different dosages: 150 mg/kg, 15 mg/kg, 1.5 mg/kg. Administered 30 min post-injury | Guinea pigs.<br>Exp. (dose-response study): $n = 20$;<br>Ctrl: $n = 21$;<br>Total: $n = 41$ | SC compression at level Th12-L1, *in vivo* | 6 weeks | SSEP, CTM | Best results with the high dose (150 mg/kg).<br>SSEP recovered in 6/7 animals at 150 mg/kg, 2/7 at 15 mg/kg, and 1/7 at 1.5 mg/kg.<br>SSEP recovery at 6 weeks: Exp.: 91%; Ctrl: 10%.<br>CTM: Recovery of measurable areflexic field area was observed only in the Exp. group—$40.8 \pm 5.3\%$ | 3D reconstruction: In the P188 group, cysts in spinal cord segments were smaller and distributed more chaotically on both sides of the central lesion, unlike in Ctrl. Lesions were more localized in the P188 group. The average volume of normal SC parenchyma in Ctrl was 41.1%, compared to 56.5% in the P188 group. On average, 24.7% of the total volume of spinal cord segments in Ctrl consisted of cysts, compared to 14% in the P188 group. The cavitation volume was also smaller in the P188 group |
| 11 | Laverty P.H.<br>et al., 2004 | A preliminary study of intravenous surfactants in paraplegic dogs: polymer therapy in canine clinical spinal cord injury | 1. PEG-3500 30% w/w in sterile saline, 2 mL/kg IV over 15 min.<br>2. P188 – 150 mg/mL, 2 mL/kg + MPSS 30 mg/kg IV. First fusogen injection during radiographic examination, second injection 4–6 hours later | Dogs.<br>PEG: $n = 19$;<br>P188: $n = 16$;<br>Ctrl: $n = 24$;<br>Total: $n = 59$ | Acute intervertebral disc herniation with paraplegia and myelopathy at level T3–L3, decompressive laminectomy, *in vivo* | 6–8 weeks | TNS, SSEP | TNS:<br>PEG: 9.1;<br>P188: 9.3;<br>Ctrl: 5.4.<br>SSEP recovery:<br>PEG: 7/11;<br>Ctrl: 0/10 | None |
| 12 | Luo J. et al., 2004 | Polyethylene glycol improves function and reduces oxidative stress in synaptosomal preparations following spinal cord injury | PEG-2000 50% w/v, topical for 2 min + Krebs solution | Guinea pigs. 5–9 per time point per group.<br>Ctrl (sham operation); Injury; Injury + PEG; PEG without injury | SC compression at level Th10-Th11, *in vivo* | 72 hours | None | PEG administration immediately after injury caused a noticeable reduction in synaptosomal oxidative stress and calcium, associated with improved synaptosomal function | FITC-labeled PEG penetrated damaged spinal cord cells better than intact cells, especially in gray matter. In damaged cells, FITC-PEG entered the cytoplasm and was evenly distributed in the cytosol |
| 13 | Luo J. and Shi R., 2004 | Diffusive oxidative stress following acute spinal cord injury in guinea pigs and its inhibition by polyethylene glycol | PEG-2000 50% w/w, topical for 5 min + Krebs solution | Guinea pigs. 5–9 per time point per group.<br>Ctrl (sham operation); Injury; Injury + PEG; PEG without injury | SC compression at level Th10-Th11, *in vivo* | 72 hours | None | Oxidative stress was assessed via lipid peroxidation, protein carbonyl content, and glutathione levels. PEG administration immediately after injury led to a noticeable reduction in oxidative stress both at the injury site and in adjacent segments | None |
| 14 | Cole A. and Shi R., 2005 | Prolonged focal application of polyethylene glycol induces conduction block in guinea pig spinal cord white matter | PEG-2000 50% w/w in distilled water, 2.5 mL, topical for 25 min + Krebs solution | Guinea pigs. Number of animals not specified | Compression of extracted SC at the mid-thoracic level, *ex vivo* | 1 hour | CAP | In the Ctrl group, by the end of the 25-min continuous PEG exposure, amplitude significantly decreased to $64 \pm 4\%$ of pre-PEG levels. After a 30-min wash with normal Krebs solution, amplitude only partially recovered to $70 \pm 5\%$ of pre-PEG levels. In the Exp. group, by the end of the 25-min continuous PEG exposure, amplitude significantly decreased to $64 \pm 7\%$ of pre-PEG levels. After a 30-min wash with normal Krebs solution, amplitude only partially recovered to $88 \pm 11\%$ | None |
|  |  |  |  |  |  |  |  | of pre-PEG levels. Despite lacking statistical significance, the compressed SC recovered slightly better than the uninjured SC after PEG application (88 ± 11% vs. 70 ± 5%) |  |
| 15 | Ditor D.S. et al., 2007 | Effects of polyethylene glycol and magnesium sulfate administration on clinically relevant neurological outcomes after spinal cord injury in the rat | PEG-3500 30% w/w in sterile saline, 1 g/kg IV + MgSO<sub>4</sub> 300 mg/kg, administered at 15 min and 6 hours post-injury | Rats, Wistar.<br>PEG: *n* = 11;<br>MgSO<sub>4</sub>: *n* = 5;<br>PEG+MgSO<sub>4</sub>: *n* = 6;<br>Ctrl: *n* = 10;<br>Total: *n* = 32 | SC compression at level Th4, *in vivo* | 6 weeks | BBB | BBB:<br>PEG: 7.3 ± 0.2;<br>MgSO<sub>4</sub>: 7.7 ± 0.4;<br>Ctrl: 6.4 ± 0.6;<br>PEG+MgSO<sub>4</sub>: 7.6 ± 0.2 | Greater volume of preserved white matter in the PEG+MgSO<sub>4</sub> group compared to Ctrl (0.68 mm<sup>2</sup> vs. 0.41 mm<sup>2</sup>).<br>Myelin volume:<br>PEG+MgSO<sub>4</sub>: 4.86 ± 0.52 mm<sup>2</sup>;<br>PEG-treated: 4.80 ± 0.72 mm<sup>2</sup>;<br>MgSO<sub>4</sub>-treated: 5.02 ± 0.41 mm<sup>2</sup>;<br>Ctrl: 4.38 ± 0.48 mm<sup>2</sup> |
| 16 | Baptiste D.C. et al., 2009 | Systemic polyethylene glycol promotes neurological recovery and tissue sparing in rats after cervical spinal cord injury | PEG-2000 30% w/w in sterile Ringer's lactate solution, IV, 1 g/kg at 15 min post-injury | Rats, Wistar.<br>Ctrl: *n* = 39;<br>Exp.: *n* = 36;<br>PEG [FITC]: *n* = 2;<br>Total: *n* = 77 | SC compression at level C7-Th1, *in vivo* | 6 weeks | BBB, IP* | BBB:<br>Ctrl: 4.5 ± 0.86;<br>Exp.: 7.2 ± 0.56.<br>IP:<br>Ctrl: 37.9 ± 1.89;<br>Exp.: 40.0 ± 2.59 | At the injury epicenter, the cross-sectional area was ~0.9 in Exp. and ~0.75 in Ctrl. Retrogradely labeled neurons in the reticular formation of animals showed that systemic PEG treatment is associated with greater axonal preservation in the injured spinal cord relative to the Ctrl group |
| 17 | Kwon B.K.<br>et al., 2009 | Magnesium chloride<br>in a polyethylene<br>glycol formulation as<br>a neuroprotective<br>therapy for acute<br>spinal cord injury:<br>preclinical refinement<br>and optimization | Exp. #1:<br>1. MgSO<sub>4</sub> in PEG<br>(500 mmol/kg MgSO<sub>4</sub><br>in 1 g/kg PEG and<br>0.5% saline);<br>2. PEG (1 g/kg PEG in<br>0.5% saline)<br>IV, first injection at<br>15 min post-injury,<br>second at 6 h.<br>Exp. #2:<br>1. MgCl<sub>2</sub> low-dose<br>(127 mmol/kg MgCl<sub>2</sub><br>in 1 g/kg PEG);<br>2. MgCl<sub>2</sub> high-dose<br>(254 mmol/kg MgCl<sub>2</sub><br>in 2 g/kg PEG);<br>3. MgSO<sub>4</sub> low-dose<br>(127 mmol/kg MgSO<sub>4</sub><br>in 1 g/kg PEG)<br>IV, different number<br>of infusions, at 8 h.<br>Exp. #3:<br>MgCl<sub>2</sub><br>(190 mmol/kg MgCl<sub>2</sub><br>in 1.65 g/kg PEG)<br>IV, different number<br>of infusions and timing | Rats,<br>Sprague-Dawley.<br>Exp. #1:<br>MgSO<sub>4</sub> in PEG:<br>*n* = 10;<br>MgSO<sub>4</sub>: *n* = 10;<br>PEG: *n* = 10;<br>Saline: *n* = 10;<br>Sham: *n* = 10;<br>Total: *n* = 50.<br>Exp. #2:<br>MgCl<sub>2</sub> low-dose,<br>2 inf: *n* = 5;<br>MgCl<sub>2</sub> low-dose,<br>4 inf: *n* = 10;<br>MgCl<sub>2</sub> low-dose,<br>6 inf: *n* = 10;<br>MgCl<sub>2</sub> high-dose,<br>2 inf: *n* = 5;<br>MgSO<sub>4</sub> low-dose,<br>2 inf: *n* = 5;<br>MgCl<sub>2</sub> low-dose,<br>6h, 2 inf: *n* = 10;<br>Saline: *n* = 10;<br>Sham: *n* = 5;<br>Total: *n* = 60.<br>Exp. #3:<br>MgCl<sub>2</sub> in PEG<br>15min, 5 inf: *n* = 5;<br>MgCl<sub>2</sub> in PEG<br>2h, 5 inf: *n* = 10;<br>MgCl<sub>2</sub> in PEG<br>4h, 5 inf: *n* = 10; | Contusion SCI at<br>the T10 level,<br>*in vivo* | 6 weeks | BBB | Exp. #1:<br>MgSO<sub>4</sub> in PEG: 8.78 ± 0.32;<br>MgSO<sub>4</sub>: 7.78 ± 0.32;<br>PEG: 8.3 ± 0.65;<br>Saline: 7.5 ± 0.11;<br>Sham: 12.<br>Exp. #2:<br>MgCl<sub>2</sub> low-dose, 2 inf:<br>8.6 ± 0.68;<br>MgCl<sub>2</sub> low-dose, 4 inf:<br>8.5 ± 0.45;<br>MgCl<sub>2</sub> low-dose, 6 inf:<br>9.4 ± [value missing];<br>MgCl<sub>2</sub> high-dose, 2 inf:<br>10 ± 0.71;<br>MgSO<sub>4</sub> low-dose, 2 inf:<br>8.2 ± 0.37;<br>MgCl<sub>2</sub> low-dose, 6h, 2 inf:<br>9.2 ± 0.58;<br>Saline: 7.2 ± 0.54;<br>Sham: 12.<br>Exp. #3:<br>MgCl<sub>2</sub> in PEG 15min, 5 inf:<br>9.2 ± 0.37;<br>MgCl<sub>2</sub> in PEG 2h, 5 inf:<br>10 ± 0.24;<br>MgCl<sub>2</sub> in PEG 4h, 5 inf:<br>8.8 ± 0.29;<br>MgCl<sub>2</sub> in PEG 8h, 5 inf:<br>8.2 ± 0.25;<br>MP* 2h, 5 inf: 8.6 ± 0.51;<br>MP+MgCl<sub>2</sub> in PEG 2h, 5 inf:<br>9.5 ± 0.27;<br>MgCl<sub>2</sub> in PEG 2h, 5 inf:<br>9.4 ± 0.26; | A strong negative correlation<br>was observed between lesion<br>volume and neurological<br>recovery. Neurological<br>recovery in treatment groups<br>was likely due to reduced<br>secondary damage and,<br>consequently, smaller lesion<br>volume compared to control<br>groups, which showed<br>significant cavitation |
|  |  |  |  | MgCl<sub>2</sub> in PEG 8h, 5 inf: $n = 10$;<br>MP* 2h, 5 inf: $n = 5$;<br>MP + MgCl<sub>2</sub> in PEG 2h, 5 inf: $n = 10$;<br>MgCl<sub>2</sub> in PEG 2h, 5 inf: $n = 5$;<br>MgCl<sub>2</sub> in PEG 2h, 2 inf: $n = 5$;<br>Saline: $n = 10$;<br>Sham: $n = 5$;<br>Total: $n = 75$.<br>Summary:<br>$n = 185$ |  |  |  | MgCl<sub>2</sub> in PEG 2h, 2 inf: $8.6 \pm 0.26$;<br>Saline: $7.8 \pm 0.36$;<br>Sham: 12 |  |
| 18 | Cho Y. et al., 2010 | Chitosan produces potent neuroprotection and physiological recovery following traumatic spinal cord injury | Chitosan, subcutaneous single dose | Guinea pigs.<br>Exp.: $n = 8$;<br>Ctrl: $n = 6$;<br>Total: $n = 14$ | Forceps compression at the mid-thoracic SC level, *in vivo* | 2 weeks | SSEP | Recovery in Exp.: 6/8 (two animals died after start of recording);<br>Ctrl: 0/6 | Chitosan seals membranes, inhibits intracellular lactate dehydrogenase leakage, and provides neuroprotection by preventing progressive reactive oxygen species formation and lipid peroxidation |
| 19 | Lee J.H.T. et al., 2010 | Magnesium in a polyethylene glycol formulation provided neuroprotection after unilateral cervical spinal cord injury | 190 mmol/kg MgCl<sub>2</sub> in 1 g/kg PEG, IV at 2 h over 10 min, 5 infusions every 6 h | Rats, Sprague-Dawley.<br>Exp.: $n = 11$;<br>Ctrl: $n = 11$;<br>Total: $n = 22$ | Contusion, hemisection at the C4-C5 level, *in vivo* | 6 weeks | Horizontal ladder test;<br>Cylinder rearing test;<br>Catwalk gait analysis | Horizontal Ladder Test:<br>At 6 weeks, the Exp. group had significantly fewer forelimb errors compared to Ctrl (3.84% vs. 11.02%)<br>Cylinder Rearing Test:<br>At 2, 4 and 6 weeks post-injury, use of the left forelimb<br>did not differ significantly between groups<br>Catwalk Gait Analysis:<br>At 6 weeks post-injury, no significant differences were found between groups for any gait parameters | The degree of preservation of white and gray matter was higher in Exp. than Ctrl. None of these differences were statistically significant at any individual site. However, cumulative white matter preservation over 1600 mm caudal and rostral sections showed significantly greater white matter sparing in Exp. vs. Ctrl |
| 20 | Kouhzaei S. et al., 2013a | The neuroprotective ability of polyethylene glycol is affected by temperature in ex vivo spinal cord injury model | 1. PEG-400;<br>2. PEG-1000;<br>3. PEG-2000;<br>50% w/w in Krebs solution, topical, at different temps (25, 37, 40°C). | Rats, Wistar.<br>5–6 animals per group<br>Total: $n = 47$ | SC compression at L1 level, *ex vivo* | 5 minutes | CAP | Best results at 40°C:<br>PEG-400: $65.46 \pm 5\%$ ($n=6$);<br>PEG-1000: $41.49 \pm 2.4\%$ ($n=5$);<br>PEG-2000: $37.36 \pm 1.6\%$ ($n=5$) | No |
| 21 | Kouhzaei S. et al., 2013b | Protective effect of low molecular weight polyethylene glycol on the repair of experimentally damaged neural membranes in rat's spinal cord | 1. PEG-200;<br>2. PEG-400;<br>3. PEG-600;<br>4. PEG-1000;<br>5. PEG-2000;<br>50% w/w in ddH<sub>2</sub>O for SCEP* or 10%-70% w/w in serum-free DMEM for cell cultures, topical for 5 min + Krebs solution | Rats, Wistar.<br>PEG-200: $n = 7$;<br>PEG-400: $n = 6$;<br>PEG-600: $n = 6$;<br>PEG-1000: $n = 6$;<br>PEG-2000: $n = 6$;<br>Total: $n = 31$ | SC compression at levels Th1-2, Th11, and L1-2, *in vivo* | 25 minutes | SCEP, CAP | SCEP:<br>PEG-200: 49.5%;<br>PEG-2000: 16.3%. Lower molecular weight PEG showed higher recovery.<br>CAP:<br>PEG-200: $49.5 \pm 3.9\%$;<br>PEG-400: $38.9 \pm 5.3\%$;<br>PEG-600: $24.4 \pm 3.4\%$;<br>PEG-1000: $17.6 \pm 4.5\%$;<br>PEG-2000: $16.3 \pm 2.3\%$ | MTT assay of mechanically damaged cell cultures showed smaller-molecular-weight PEGs provide higher degree of membrane sealing: $77.8 \pm 3.5$ for PEG-400 (20% w/w) compared to $32.1 \pm 6.9$ for PEG-2000 (20% w/w). Large-molecular-weight PEGs (PEG-1000, -2000), which provided minor regeneration at low concentrations, showed no significant sealing effect at high concentrations (>50% w/w) |
| 22 | Estrada V. et al., 2014 | Long-lasting significant functional improvement in chronic severe spinal cord injury following scar resection and polyethylene glycol implantation | PEG-600, topical, after scar resection or Matrigel, or alginate hydrogel (ALG) | Rats, Wistar.<br>No exact animal numbers.<br>PEG: $n = 14$;<br>PEG re-transection: $n = 3$;<br>Ctrl: $n = 14$ | Chronic SCI: Dorsal hemisection at T9, observation for 5 weeks, scar excision with fusogen application, re-hemisection at 8 weeks, *in vivo* | 8 months | mBBB | PEG: Mean 8.5, Max 20.<br>Ctrl: ~3.<br>PEG vs. Ctrl, reached certain mBBB thresholds, gradually increasing from 5, 10, 15, to 20. Motor improvement in the PEG group (mean score 8.5) was significantly reduced to a mean score of 1.8 after re-hemisection, indicating significant contribution of regenerated/myelinated axons to motor improvement in PEG-treated animals | Matrigel showed little/no axonal ingrowth into biopolymer matrix and was excluded from further analysis. The ALG group showed few axonal profiles. Quantification of NF+ axonal profiles at 5 weeks post-op showed no significant differences between ALG and Ctrl groups. The PEG group showed the highest axon density. Axons from all analyzed fiber populations grew into the PEG bridge. Ctrl and ALG groups had only few small-diameter blood vessels, whereas the PEG group showed dense distribution of large-diameter vessels throughout the matrix area |
| 23 | Oda Y. et al., 2014 | Effects of polyethylene glycol administration and bone marrow stromal cell transplantation therapy in spinal cord injury mice | PEG-4000 50% w/v in PBS, 10 μL injected at six points rostral and caudal to the injury site | Mice, ICR.<br>BMSC+PEG: $n = 8$;<br>PEG: $n = 8$;<br>BMSC: $n = 8$;<br>Ctrl: $n = 8$;<br>Total: $n = 36$ | Complete transection at Th10 with BMSC* transplantation, with PEG, *in vivo* | 4 weeks | BBB | BMSC: ~10;<br>BMSC+PEG: ~11;<br>PEG: ~11;<br>Ctrl: ~8 | In BMSC and BMSC+PEG groups, neuronal cells aggressively migrated toward the glial scar from areas rostral to the lesion. In Ctrl and PEG groups, the border of damaged areas was covered by astrocytes and few neuronal cells migrated toward the glial scar |
| 24 | Bittner G.D.<br>et al., 2015 | Application and implications of polyethylene glycol-fusion as a novel technology to repair injured spinal cords | PEG-3250, 500 mM in ddH<sub>2</sub>O*, topical 2 min + Ca<sup>2+</sup>-free Krebs with EGTA + MB* + phys. sol. with Ca<sup>2+</sup> | Rats, Lewis.<br>Exp.: $n = 6$;<br>Ctrl: $n = 6$;<br>Total: $n = 12$ | Blunt SCI at the Th9-Th10 level, *in vivo* | 5 weeks | BBB | Exp.: ~12.2;<br>Ctrl: ~8.2 | No |
| 25 | Kim C-Y.,<br>2016 | PEG-assisted reconstruction of the cervical spinal cord in rats: effects on motor conduction at 1 h | PEG-400, topical on SC and blade prior to transection | Rats, Sprague-Dawley.<br>Exp.: $n = 7$;<br>Ctrl: $n = 7$;<br>Total: $n = 14$ | Complete SC transection at C5, *in vivo* | 1 hour | MEP | Exp.: $156.7 \pm 7.6 \mu\text{V}$;<br>Ctrl.: $81.1 \pm 12.4 \mu\text{V}$ | No |
| 26 | Kim C-Y.<br>et al., 2016 | Accelerated recovery of sensorimotor function in a dog submitted to quasi-total transection of the cervical spinal cord and treated with PEG | PEG-400, topical | Dog, Beagle.<br>Total: $n = 1$ | SC transection at the C5 level, *in vivo* | 3 weeks | MFS for dogs | Exp.: 4–5 | No |
| 27 | Olby N.J.<br>et al., 2016 | A placebo-controlled, prospective, randomized clinical trial of polyethylene glycol and methylprednisolone sodium succinate in dogs with | PEG-3500 30% w/w in sterile saline, 2 mL/kg IV over 15 min immediately and at 4 h | Dogs, various breeds.<br>PEG: $n = 24$;<br>MPSS: $n = 21$;<br>Ctrl: $n = 18$; | Traumatic IVDD (Intervertebral Disc Disease in Dogs) in thoracic region (<24 h old), paraplegia, no deep pain sensation, | 12 weeks | Open field score, Ambulation ability | Open field score:<br>PEG: 5.75 (0–9.5);<br>MPSS: 5 (0–11.5);<br>Ctrl: 7.5 (1–11.5).<br>Ambulation ability:<br>PEG: 12/24;<br>MPSS: 11/21;<br>Ctrl: 7/18 | No |
| | | intervertebral disk herniation | | Total: $n = 63$ | hemilaminectomy, *in vivo* | | | | |
| 28 | Streijger F. et al., 2016 | The evaluation of magnesium chloride within a polyethylene glycol formulation in a porcine model of acute spinal cord injury | ACP105 (MgSO<sub>4</sub> + PEG-3350) at 4 h post-injury, IV over 30 min every 6 h for 5 doses | Pigs, Yucatan minipigs.<br>ACP105: $n = 6$;<br>MgSO<sub>4</sub>: $n = 6$;<br>Saline: $n = 6$;<br>Sham: $n = 4$;<br>Total: $n = 22$ | Contusion with 50g weight + additional 100g compression, laminectomy, transpedicular fixation, *in vivo* | 12 weeks | PTIBS | ACP105: $3.33 \pm 0.42$;<br>MgSO<sub>4</sub>: $3.50 \pm 0.43$;<br>Saline: $3.33 \pm 0.21$.<br>No significant differences in PTIBS scores between treatment groups at any time point | Significant destruction of gray/white matter was observed at the epicenter. The lesion core length at T10 was ~6.4 mm in the Saline group, and extended to 8.0 mm and 9.6 mm in the AC105 and MgSO<sub>4</sub> groups, respectively. Quantification of spared tissue area showed no differences between the Saline and MgSO<sub>4</sub> groups. Although edema was less severe in the AC105 group, the difference was not significant. White/gray matter at the epicenter was destroyed in all the groups, with gradual recovery with increasing distance. Small areas of gray/white matter were observed 4.0–7.2 mm rostral to the epicenter after AC105 treatment |
| 29 | Ye Y. et al., 2016 | Fusogen-assisted rapid reconstitution of anatomophysiologic continuity of the transected spinal cord | 1. PEG-1500;<br>2. PEG-4000;<br>50% in distilled water, topical | Mice, C57 wild-type.<br>PEG-1500: $n = 8$;<br>PEG-4000: $n = 8$;<br>Ctrl: $n = 8$; | Dorsal column transection at T10, *in vivo* | 4 weeks | BBB, SSEP | BBB:<br>PEG-1500: 14;<br>PEG-4000: 10;<br>Ctrl: 2<br>SSEP:<br>PEG-1500: 8/8;<br>PEG-4000: 8/8;<br>Ctrl: 0/8 | No |
| | | | | Total: $n = 24$ | | | | | |
| 30 | Huang Z. et al., 2017 | AC105 increases extracellular magnesium delivery and reduces excitotoxic glutamate exposure within injured spinal cords in rats | ACP105 (MgCl<sub>2</sub> + PEG-3350) at 2 h post-injury, IV over 30 min single or 5× every 6 h | Rats, Long-Evans. 3 groups of 12-15 animals: ACP105, MgSO<sub>4</sub>, Saline, Sham | SC compression at T10, *in vivo/in vitro* | 32 hours | None | Microdialysis showed AC105 reduced extracellular glutamate exposure parallel to increased Mg<sup>2+</sup> delivery post-SCI. Repeated AC105 fully/significantly restored extracellular Mg<sup>2+</sup> concentration post-injury to sham levels, whereas saline did not. Multiple infusions of equimolar MgSO<sub>4</sub> only slightly increased extracellular Mg<sup>2+</sup> concentration | No |
| 31 | Kim C-Y. et al., 2017 | Immunohistochemical evidence of axonal regrowth across polyethylene glycol-fused cervical cords in mice | PEG-400, topical | Mice, ICR. No animal numbers provided | SC transection at C5, *in vivo* | 4 weeks | None | No | Longitudinal sections showed clear signs of axonal sprouting across the gap |
| 32 | Ren S. et al., 2017 | Polyethylene glycol-induced motor recovery after total spinal transection in rats | PEG-600 100%, 0.5 mL, topical | Rats, strain unknown.<br>Exp.: $n = 9$;<br>Ctrl: $n = 6$;<br>Total: $n = 15$ | SC transection at Th10, *in vivo* | 4 weeks | BBB | Exp.: 12;<br>Ctrl: 4.4 | No |
| 33 | Liu Z. et al., 2018 | Restoration of motor function after operative reconstruction of the | PEG-600, 2 mL, topical | Dogs, Beagle.<br>Exp.: $n = 7$; | SC transection at Th10, *in vivo* | 60 days | cBBB | Exp.: 8 (range 5 to 18);<br>Ctrl: 3 (2–4) | No |
| | | acutely transected spinal cord in the canine model | | Ctrl: $n = 5$;<br>Total: $n = 12$ | | | | | |
| 34 | Kim C.-Y. et al., 2018 | Effect of Graphene Nanoribbons (TexasPEG) on locomotor function recovery in a rat model of lumbar spinal cord transection | "TexasPEG" - PEG-600 in pegylated graphene nanoribbons PEG-GNR, topical | Rats, Sprague-Dawley.<br>Exp.: $n = 10$;<br>Ctrl: $n = 10$;<br>Total: $n = 20$ | SC transection at L1, *in vivo* | 5 weeks | BBB | Exp.: 8 (max 9);<br>Ctrl: <4 | Increased number of NF200-positive fibers in the injury zone, reflecting axonal regeneration. GFAP expression was reduced in Exp., indicative of limited glial scar formation and reduced reactive gliosis vs. Ctrl |
| 35 | Ren S. et al., 2019 | Reconstruction of the spinal cord of spinal transected dogs with polyethylene glycol | PEG-600, 2 mL, topical | Dogs, Beagle.<br>Exp.: $n = 7$;<br>Ctrl: $n = 5$;<br>Total: $n = 12$ | SC transection at Th10, *in vivo* | 6 months | cBBB | Exp.: 11;<br>Ctrl: 4 | A significant reduction in vacuolization and cyst formation was observed in the experimental group compared with the Ctrl, which exhibited extensive glial cavities. The Exp. group showed abundant myelin preservation above, at, and below the injury site, accompanied by extensive axonal growth spanning the lesion gap. In Ctrl, myelin was almost completely destroyed in the lesion area |
| 36 | Ren S. et al., 2021 | Transplantation of a vascularized pedicle of hemisected spinal cord to establish spinal cord continuity after removal of a segment of the | PEG-600, 2 mL, topical for 5 min | Dogs, Beagle.<br>Exp.: $n = 4$;<br>Ctrl: $n = 4$; | SC transection at Th10 + transplantation of vascular-pedicle of hemisected SC, *in vivo* | 6 months | cBBB | Graft+PEG: 11.5;<br>Graft: 4 | The best results were noted in the Graft+PEG group, which displayed a greater number of axons, myelin, and healthy oligodendrocytes at the transection site. Vacuolization was minimal at |
| | | thoracic spinal cord:<br>A proof-of-principle study in dogs | | Total: $n = 8$ | | | | | the suture site in the PEG group. Many axons traversed the connective tissue area in the injury zone |
| 37 | Ren X. et al., 2022a | Partial restoration of spinal cord neural continuity *via* sural nerve transplantation using a technique of spinal cord fusion | PEG-600, topical + sural nerve transplantation | Humans.<br>PEG: $n = 12$ | Chronic complete SCI, *in vivo* | 6 months | ISNCSCI, ASIA | Neurological improvement in 5/12 patients.<br>1: ISNCSCI improved 0 to 16, ASIA A to C.<br>2: Improved 81 to 98, ASIA A to B.<br>3: Improved 79 to 84, ASIA A to B.<br>4: Improved 68 to 72, ASIA A to B.<br>5: ISNCSCI LL motor score 0 to 4, ASIA remained A | No |
| 38 | Ren X. et al., 2022b | Partial restoration of spinal cord neural continuity via vascular pedicle hemisected spinal cord transplantation using spinal cord fusion technique | PEG-600, 10 mL topical + transplantation of a vascular-pedicle of hemisected SC | Humans.<br>PEG: $n = 8$ | Chronic complete SCI, *in vivo* | 6 months | ISNCSCI, ASIA | One patient improved to ASIA C, ISNCSCI – 16; others showed improvement in autonomic functions | No |
| 39 | Lebenstein-Gumovski M. et al., 2023a | Chitosan/PEG-mediated spinal cord fusion after complete dorsal transection in rabbits - functional results at 30 days | PEG-chitosan, 2 mL topical | Rabbits.<br>Exp.: $n = 10$;<br>Ctrl: $n = 10$;<br>Total: $n = 20$ | Complete dorsal transection at Th9, *in vivo* | 40 days | mBBB | Exp.: 17;<br>Ctrl: 3 | No |
| 40 | Lebenstein-Gumovski M. et al., 2023b | PEG-chitosan (Neuro-PEG) induced restoration of motor function after complete transection of the dorsal spinal cord in swine. A pilot study | PEG-chitosan (Neuro-PEG), 2 mL topical + PEG 600 25% w/w in saline IV 50 mL for 7 days | Pigs, Mangalica.<br>Exp.: $n = 3$;<br>Ctrl: $n = 2$;<br>Total: $n = 5$ | Complete dorsal transection at Th9, laminectomy, transpedicular fixation, *in vivo* | 60 days | ILMS, PTIBS, mBBB | ILMS: Exp.: 6.6; Ctrl: 1. PTIBS: Exp.: 8; Ctrl: 1. mBBB: Exp.: 20.3; Ctrl: 2 | Ctrl: Clear transection site with axonal loss in Wallerian degeneration areas.<br>Exp: Despite cysts in gray matter, twisted/thickened axons were visualized in white matter. DiI tracing showed dye distribution across the injury site and axons crossing the transection gap |
| 41 | Lebenstein-Gumovski M. et al., 2023c | Recovery of spinal cord functions after experimental complete crossection under the effect of chitosan polymeric compounds | PEG-chitosan, 2 mL topical | Rabbits.<br>Exp.: $n = 10$;<br>Ctrl: $n = 10$;<br>Total: $n = 20$ | Complete transection at Th9, *in vivo* | 40 days | mBBB | Exp.: 20;<br>Ctrl: 0 | Ctrl: Fibro-glial transformation, glial cell proliferation, multiple cysts, focal necrosis, macrophage/fibroblast infiltration at the injury site. Disrupted axon course. Fibro-glial scar occupied entire gap, interrupting axon course. Exp: Degeneration significantly less severe. Few cysts, axon tortuosity/thickening. No significant glial proliferation/macrophage infiltration. Curved/enlarged axons crossed the injury site, visualized caudally. DiI vital dye spread in axons revealed connections between cranial/caudal axons |
| 42 | Lee J. et al., 2023 | Protection of the vascular system by polyethylene glycol reduces secondary injury following | PEG-600 30% in PBS, 20 μL, topical subdural | Rats, Sprague-Dawley.<br>Exp.: $n = 33$; | Contusion with 10g rod at Th10, *in vivo* | 6 weeks | BBB | Exp.: $15 \pm 1.5$;<br>Ctrl: $\sim 10 \pm 1.2$ | Significant reduction in dye extravasation vs. saline Ctrl, indicating restored vascular barrier integrity. Preserved expression of CD31 |
| | | spinal cord injury in rats | | Ctrl: $n = 33$;<br>Total: $n = 66$ | | | | | (endothelial marker) and ZO-1 (key tight junction protein) at epicenter, indicating support for vascular structure/endothelial tight junctions. Spared tissue area significantly larger in the Exp. group than in the Ctrl |
| 43 | Nourbakhsh A. et al., 2024 | A novel reconstruction model for thoracic spinal cord injury in swine | PEG-600, 30%, topical 0.5 mL+ sural nerve graft into SC defect | Pigs, Yucatan.<br>Exp.: $n = 3$;<br>Ctrl: $n = 2$;<br>Total: $n = 5$ | Contusion with 100g rod held for 5 min, excision of 5mm damaged SC segment at Th10, *in vivo* | 12 weeks | Sensory and motor function assessment score | PEG alone: No significant improvement.<br>PEG+sural nerve graft: Standing at week 7, reaction to physical manipulation, active knee/hip movement weeks 9–10 | Inflammation/necrosis observed in both groups, no details on axonal growth; authors relied on neuroimaging methods |
| 44 | Shen T. et al., 2024 | Developing preclinical dog models for reconstructive severed spinal cord continuity via spinal cord fusion technique | PEG-600, topical | Dogs, Beagle.<br>SCT: $n = 7$;<br>vSCT: $n = 4$;<br>vASCT: $n = 12$;<br>Total: $n = 23$ | SCT: Complete transection at Th10.<br>vSCT: 1-cm segment removal at Th10 + transposition of half SC on vascular pedicle.<br>vASCT: SC transplantation on vascular pedicle using PEG in all groups, *in vivo* | 6 months | cBBB | SCT+PEG: 11;<br>vSCT+PEG: 11.5;<br>vASCT+PEG: 10 | No |
| 45 | Zhang W. et al., 2024a | A novel strategy for spinal cord reconstruction via vascularized | PEG-600 topical + Tacrolimus 0.1 mg/kg/day + | Dogs, Beagle. Paired donor/recipient. | SC transection at Th10, transplantation of vascular-pedicle | 6 months | cBBB | Exp.: 10.63;<br>Ctrl: 0 | No |
| | | allogeneic spinal cord transplantation<br>combine spinal cord fusion | methylprednisolone<br>1 mg/kg/day | Tac+MP: $n = 8$;<br>Ctrl: $n = 4$;<br>Total: $n = 24$ | hemisected SC,<br>transplantation of<br>vascularized<br>allograft SC,<br>*in vivo* | | | | |
| 46 | Zhang W.<br>et al., 2024b | Recovery of<br>independent<br>ambulation after<br>complete spinal cord<br>transection in the<br>presence of the<br>neuroprotectant<br>polyethylene glycol in<br>monkeys | PEG-600, 5 mL topical<br>+ tacrolimus | Monkeys,<br>*Macaca fascicularis*.<br>Exp.: $n = 1$;<br>Ctrl: $n = 1$;<br>Total: $n = 2$ | Complete SC<br>transection at<br>Th10, *in vivo* | 18<br>months | Tarlov<br>scale,<br>Pritchard<br>21-point<br>scale | Exp:<br>Tarlov: 4;<br>Pritchard: 16 right, 18 left.<br>Ctrl:<br>Tarlov: 0;<br>Pritchard: 0 | Exp: Preserved structure—<br>intact neurons, axonal<br>pathways, myelin.<br>Ctrl: SC at transection level<br>with no residual neural tissue |

**Table 2.**
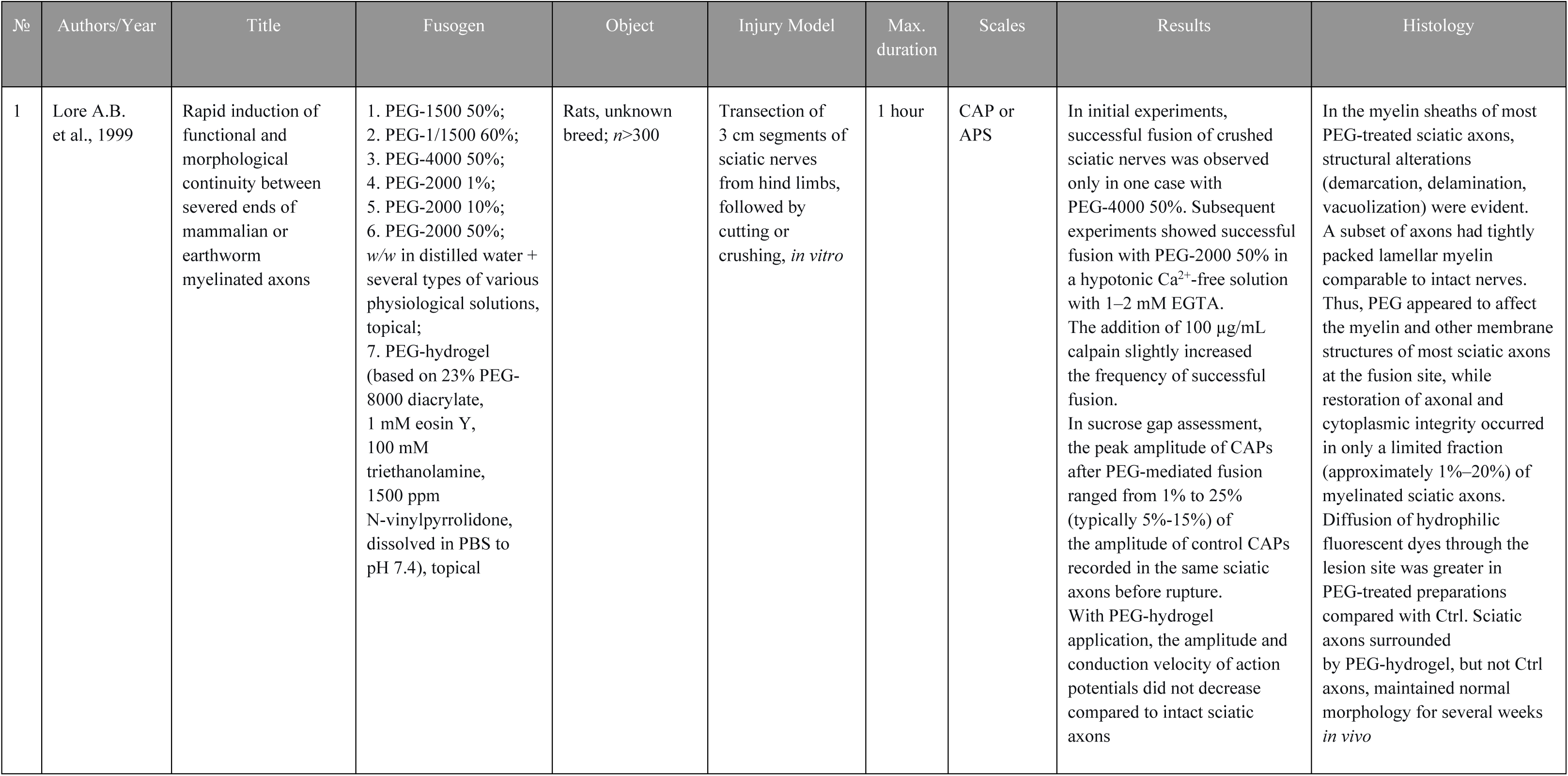
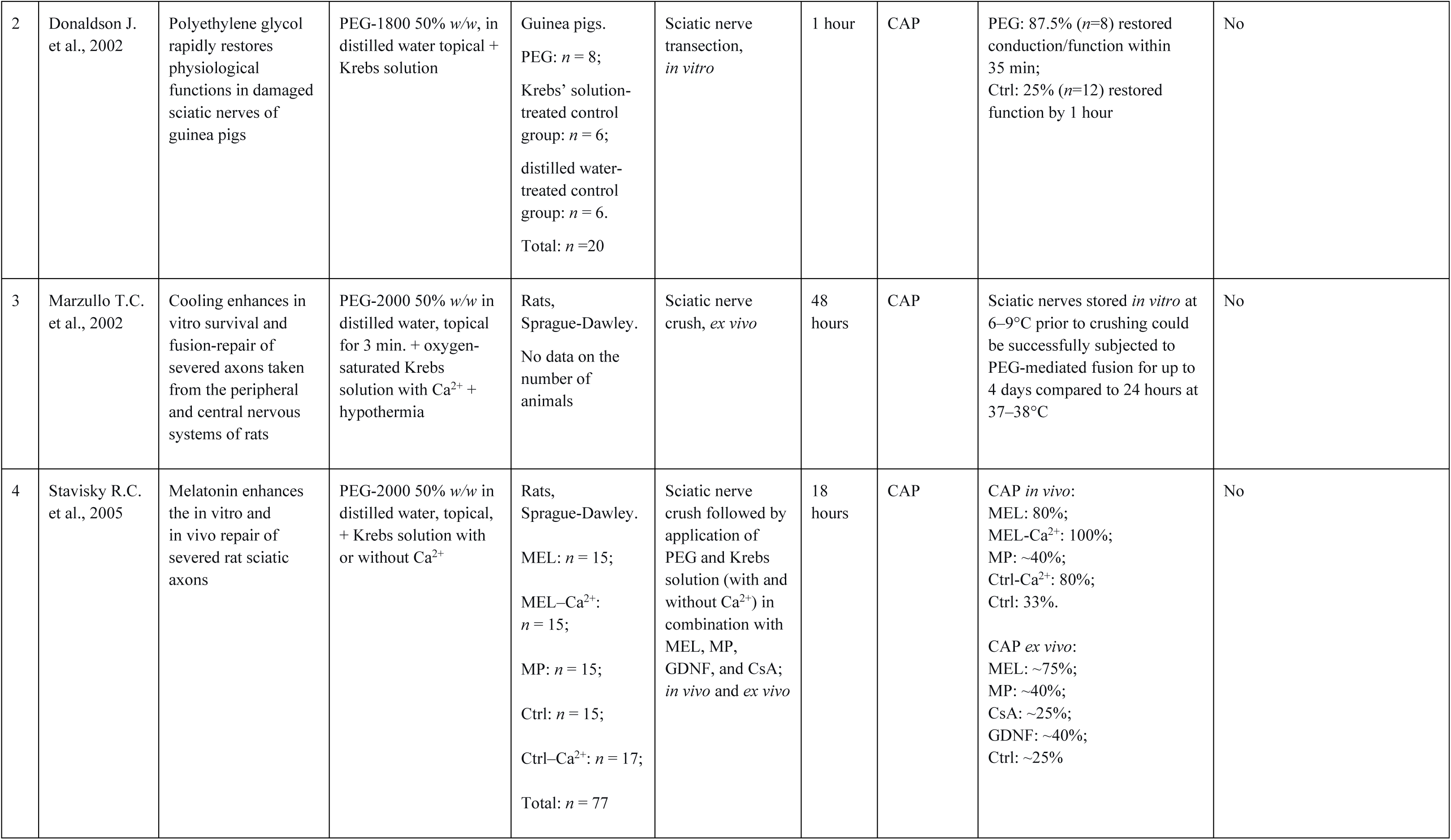
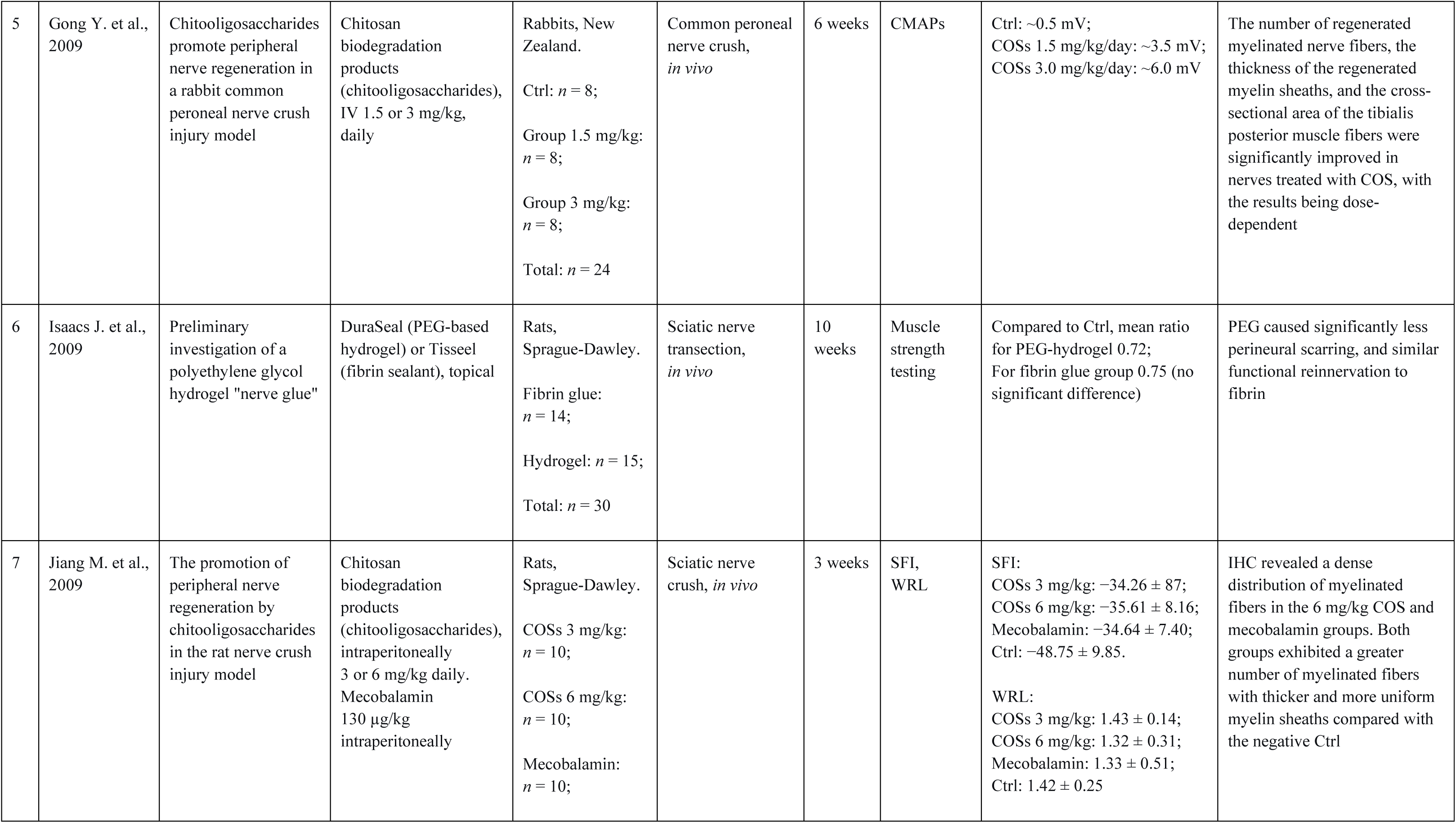
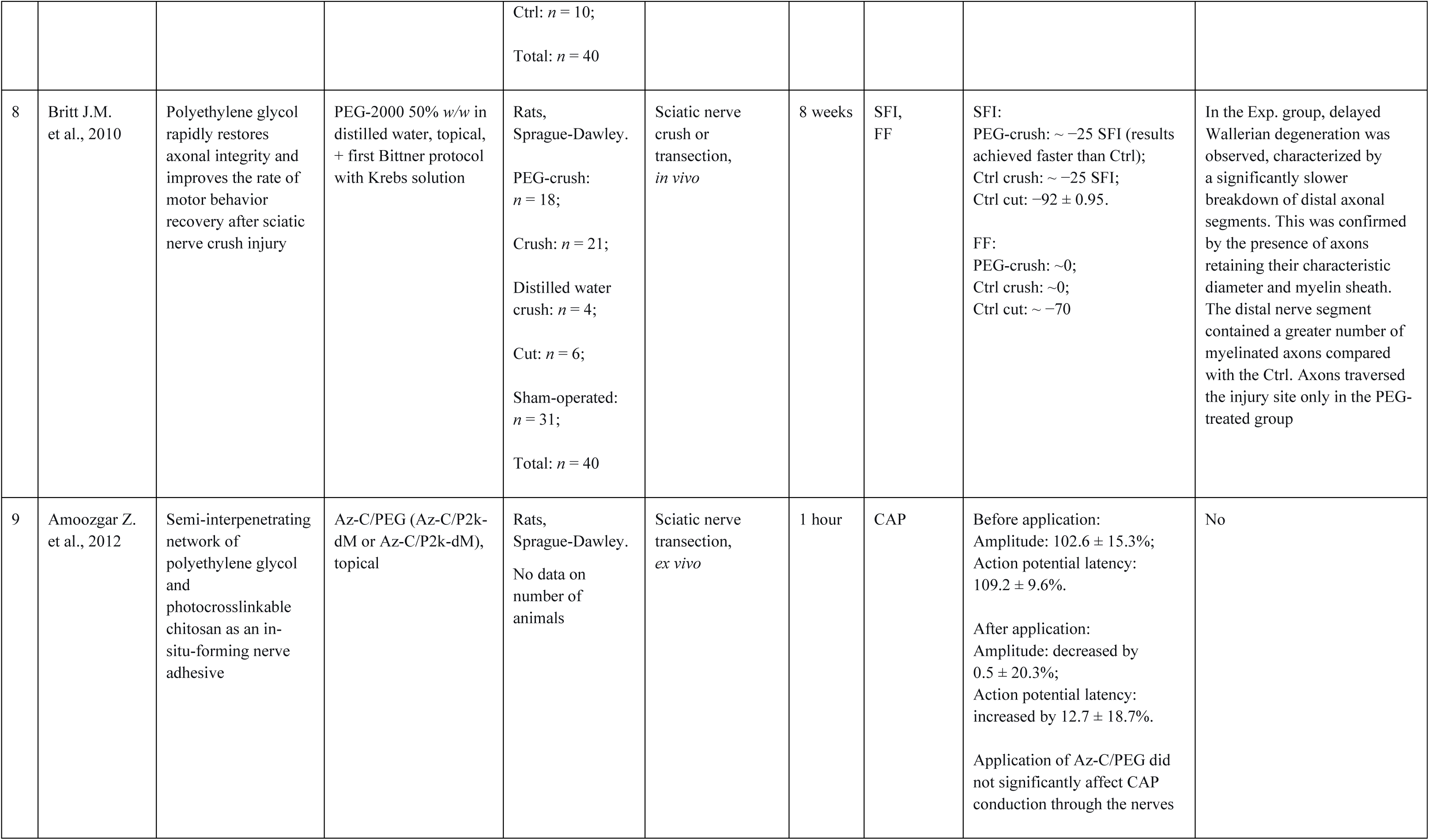
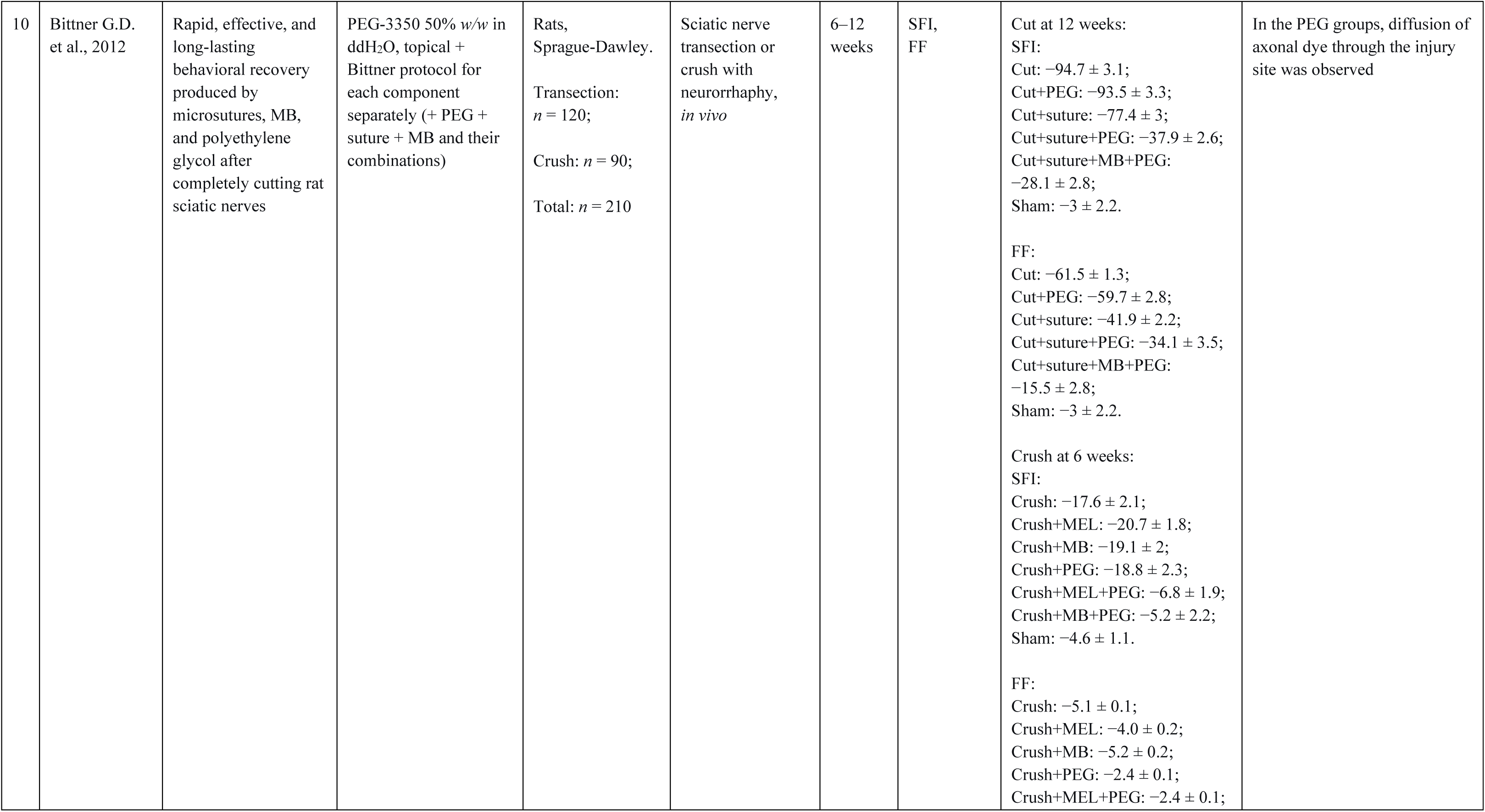
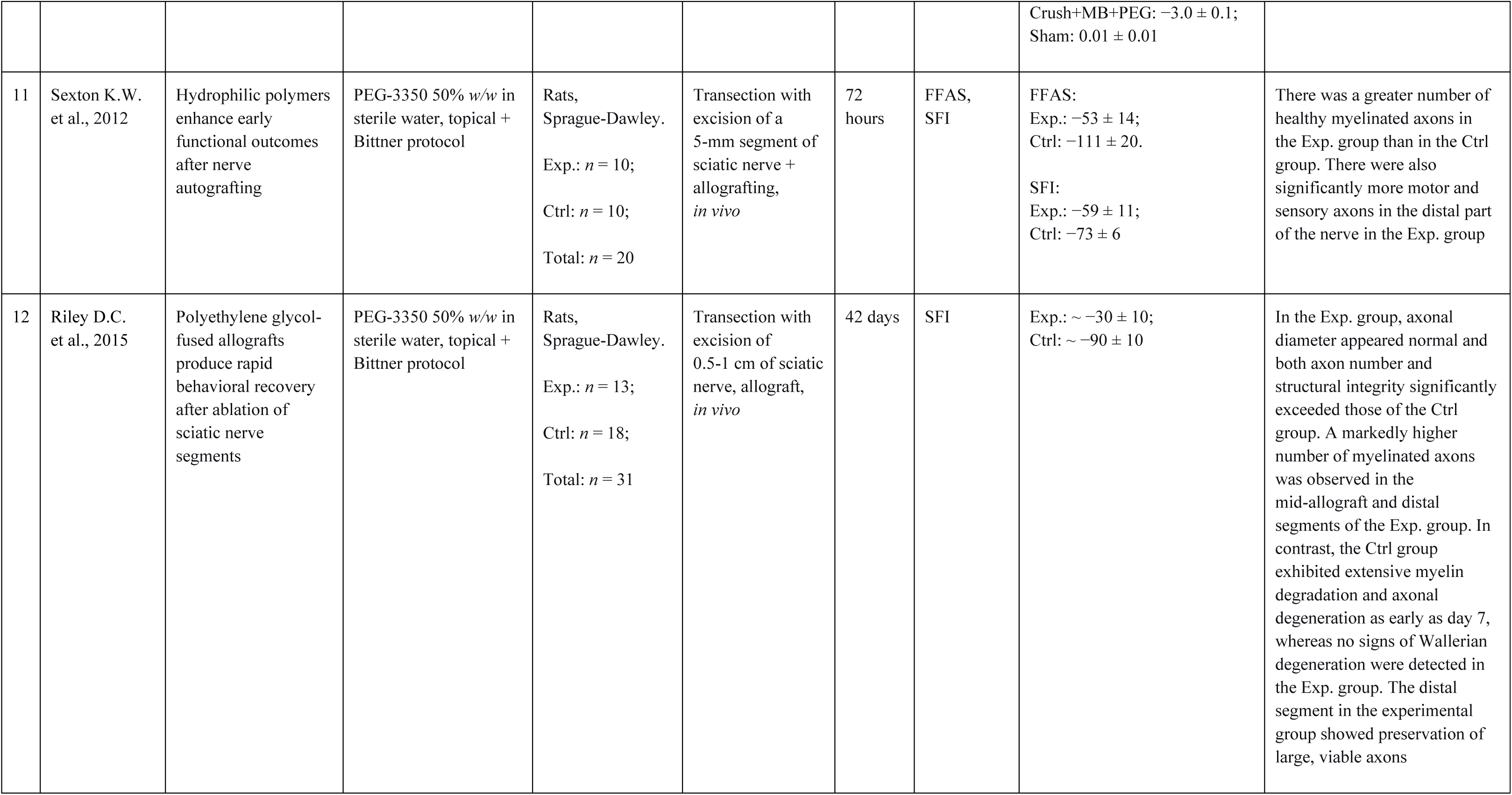
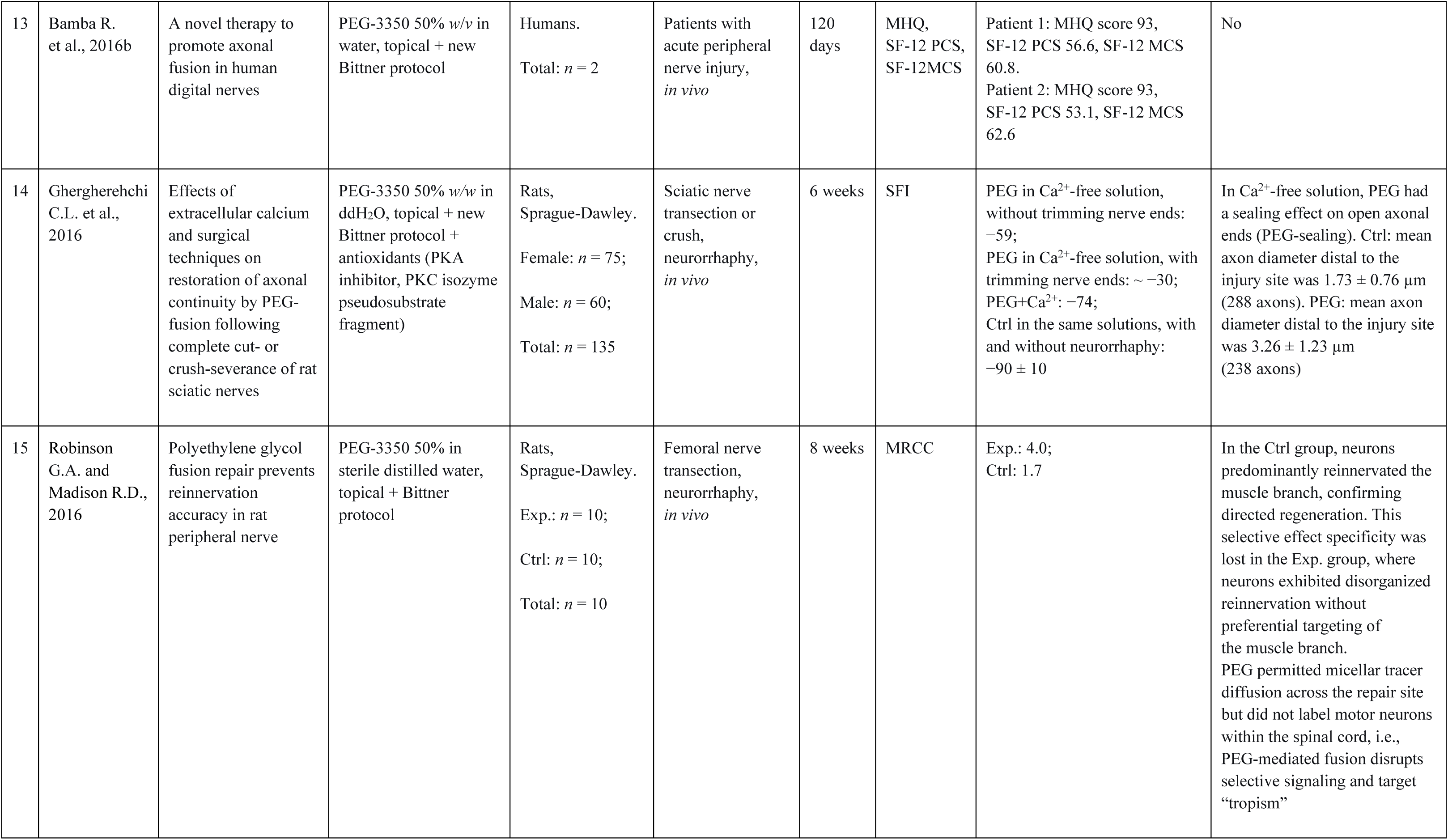
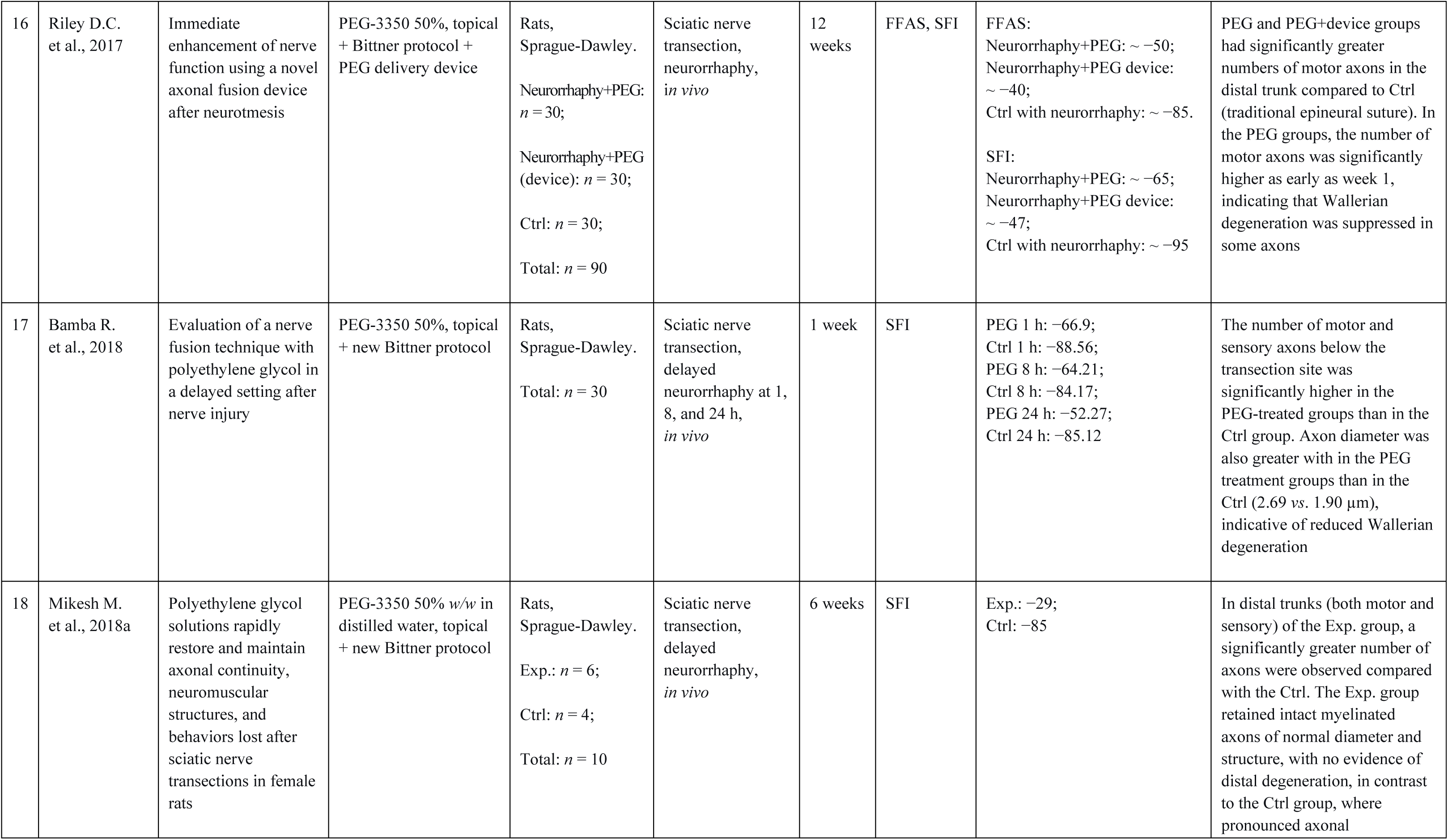
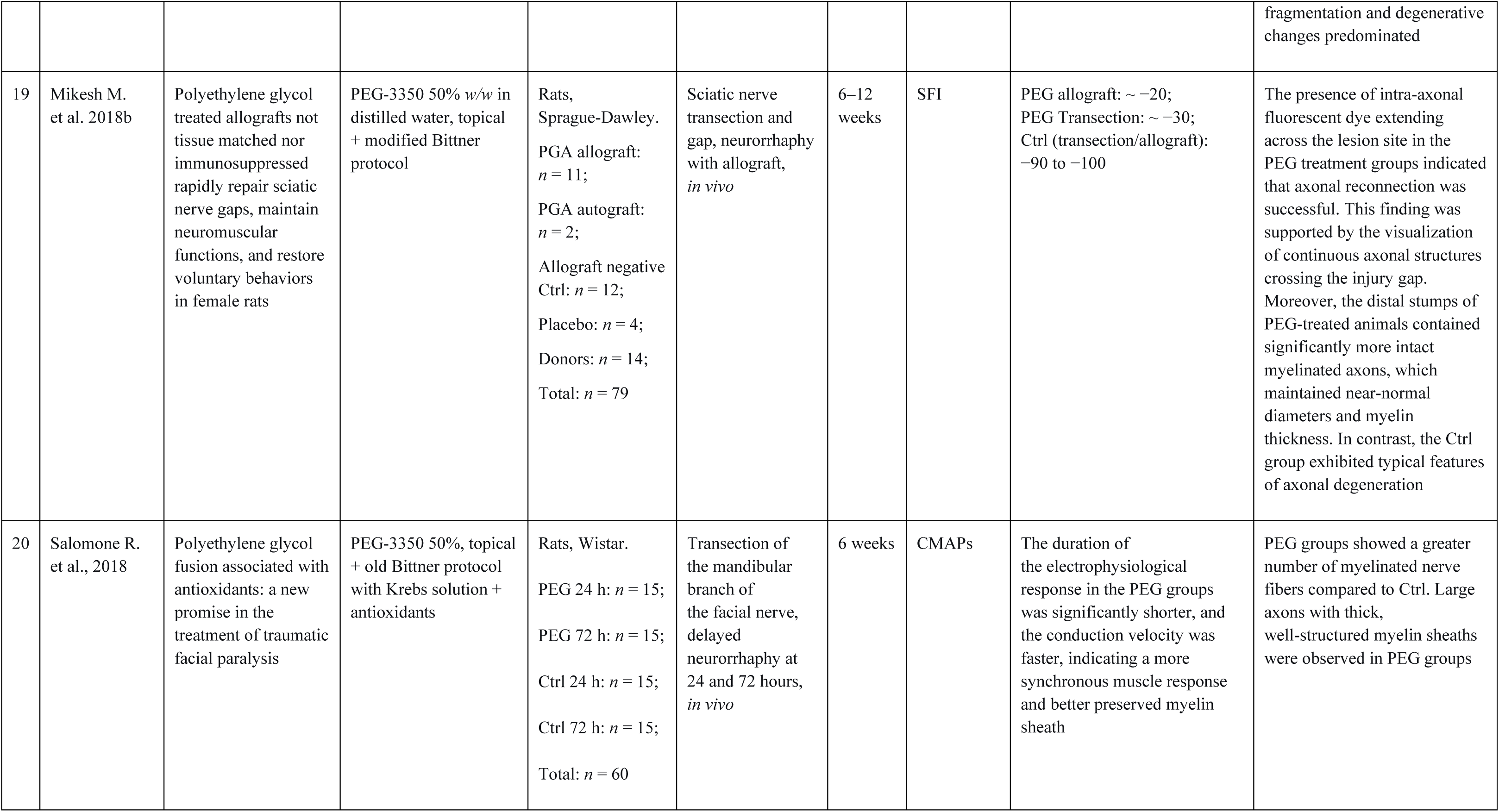
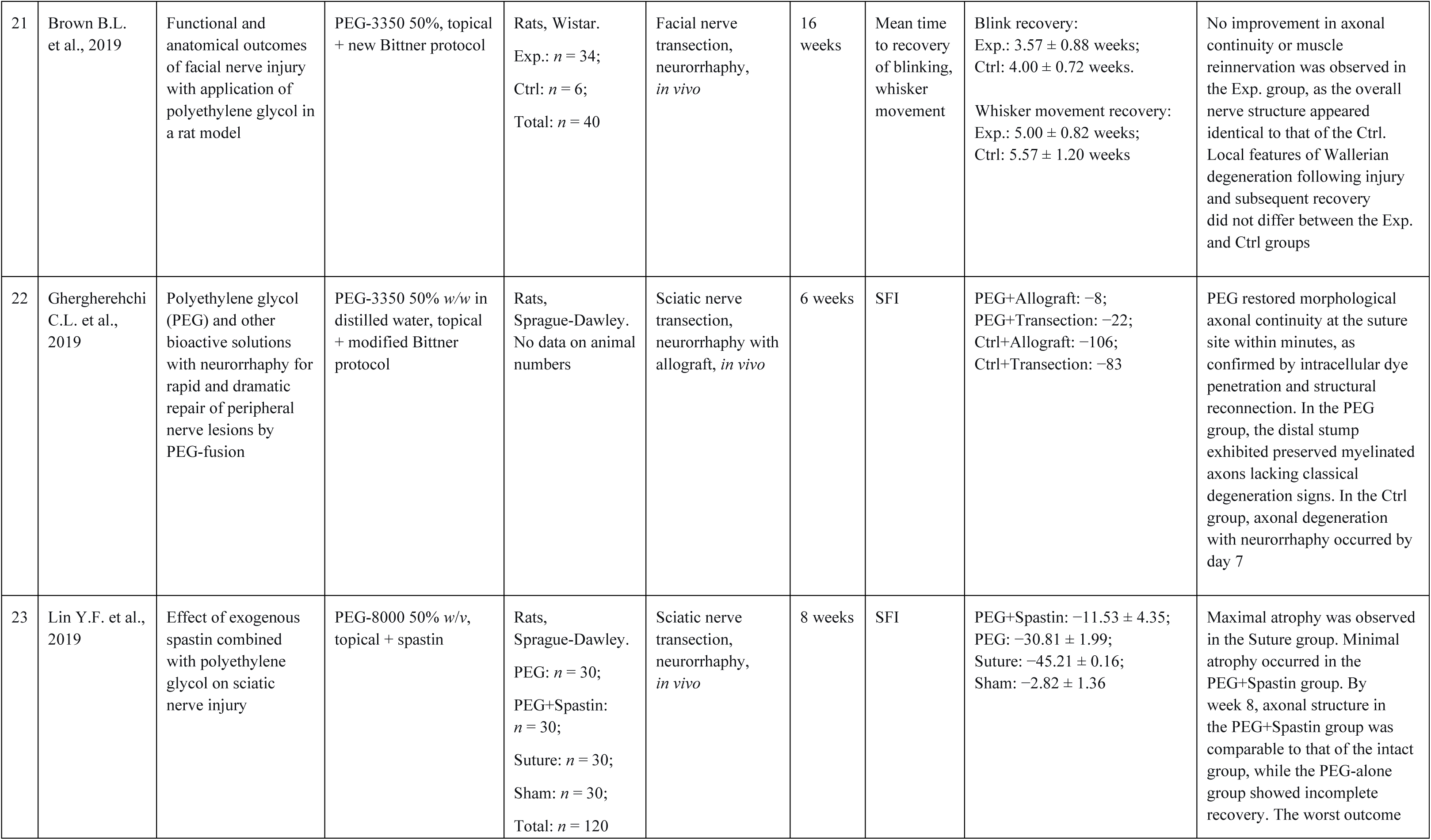
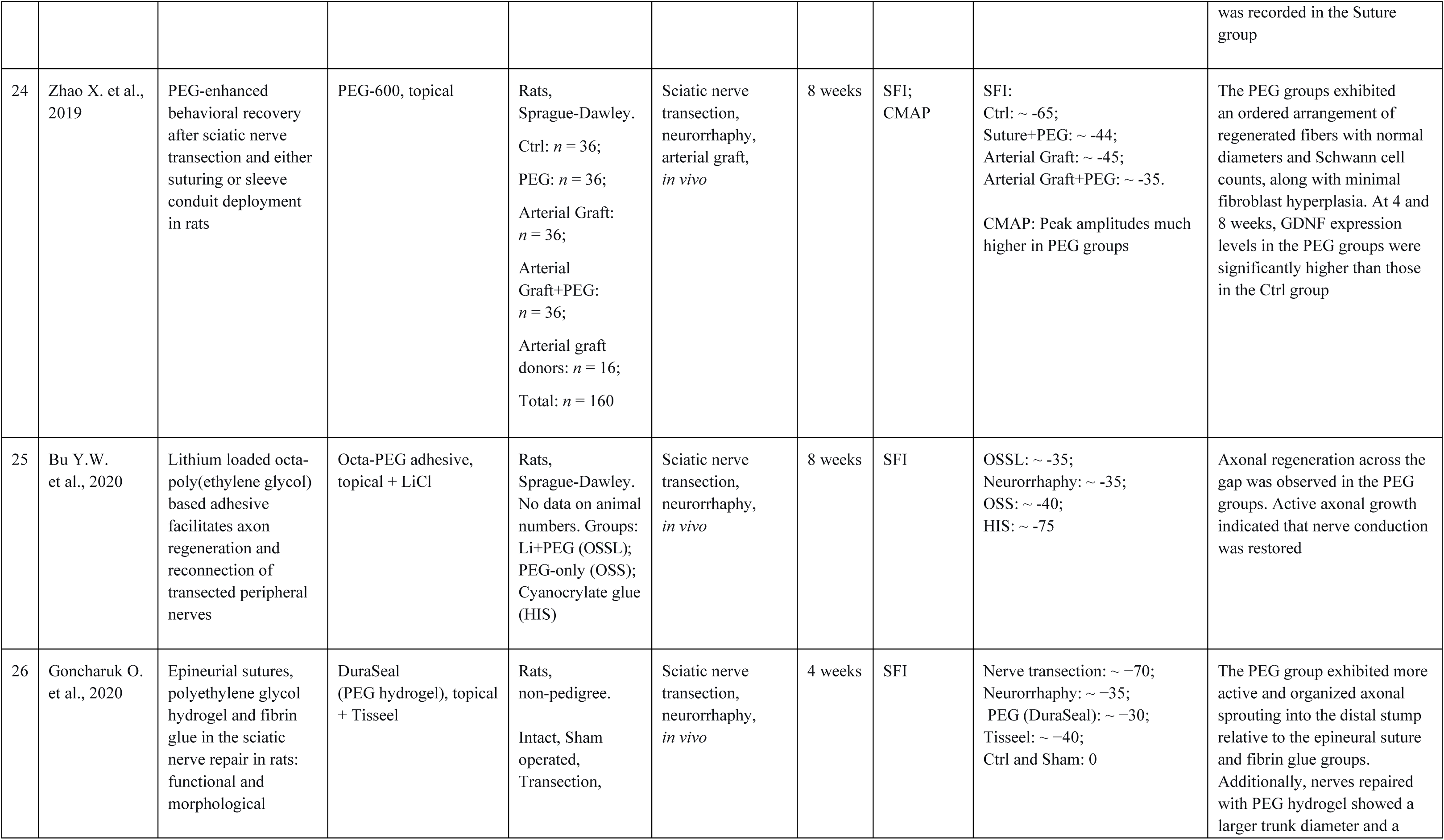
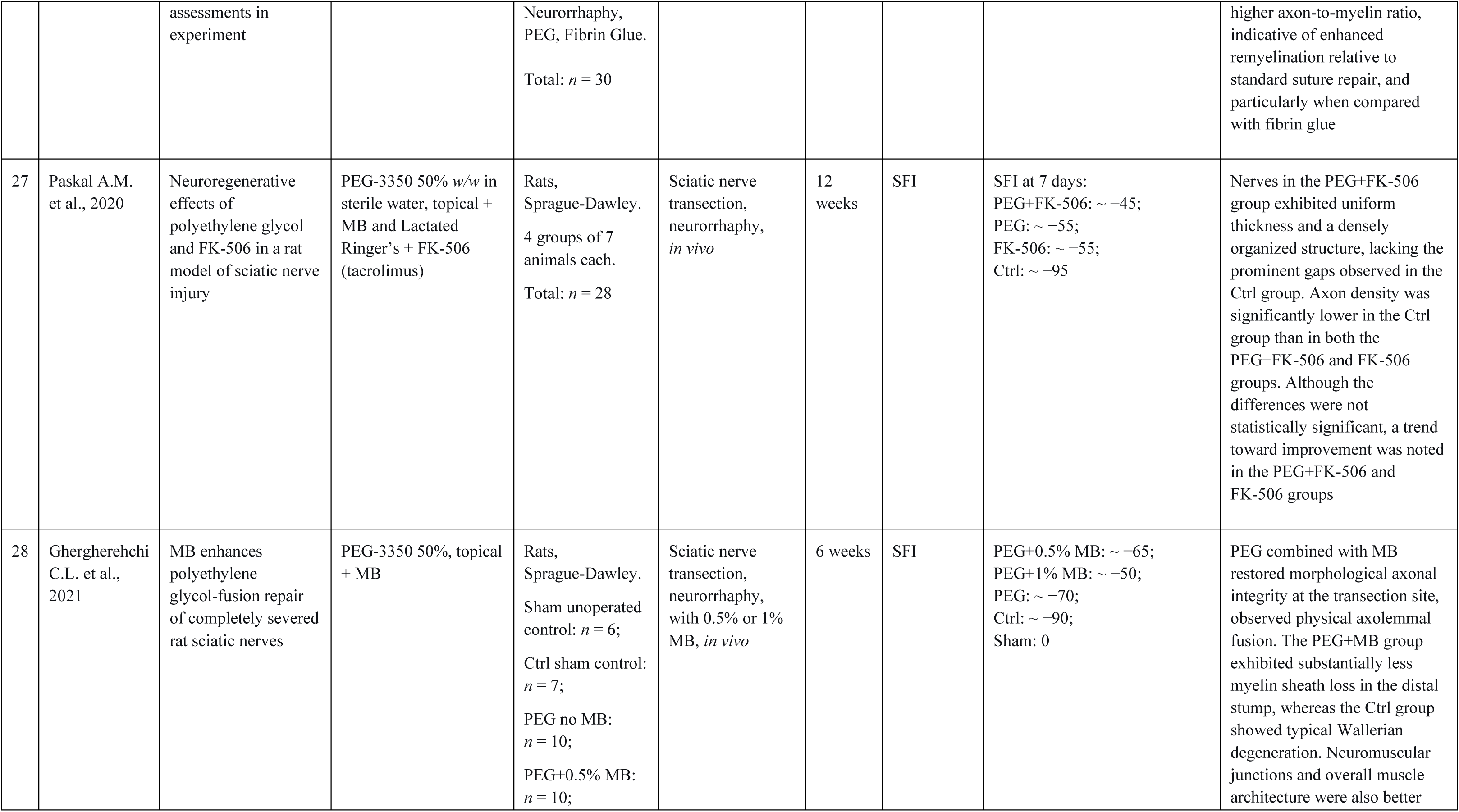
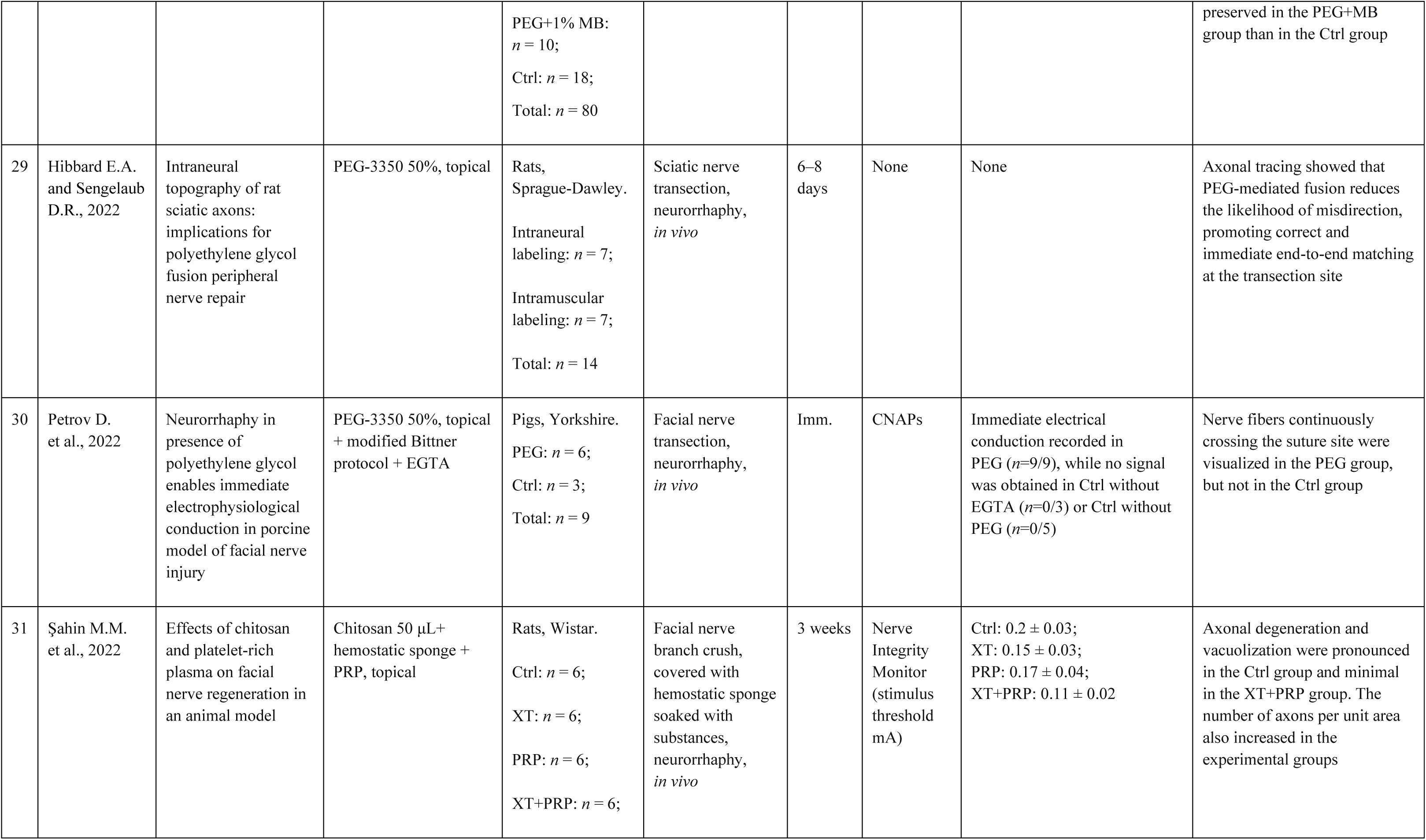
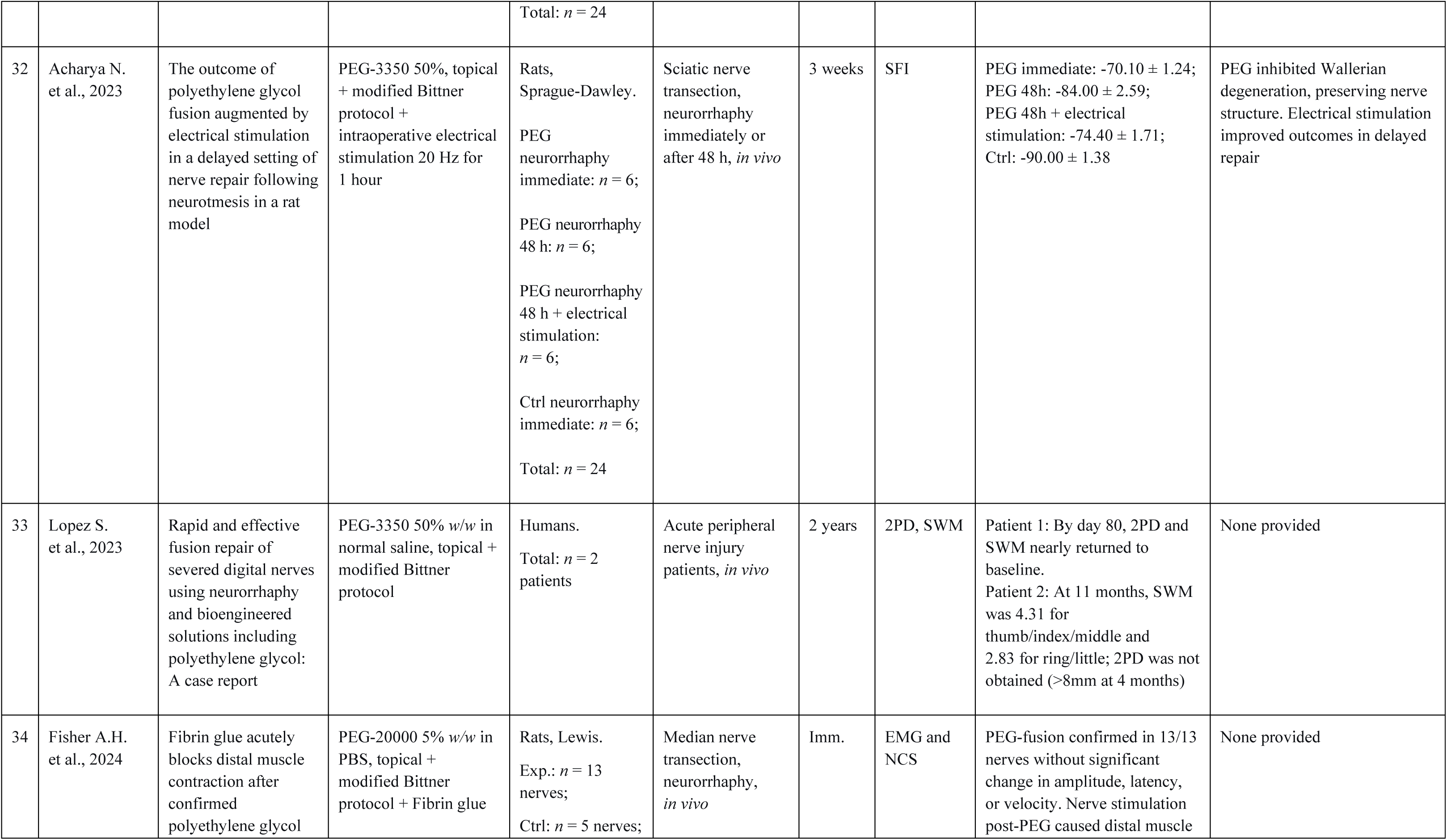
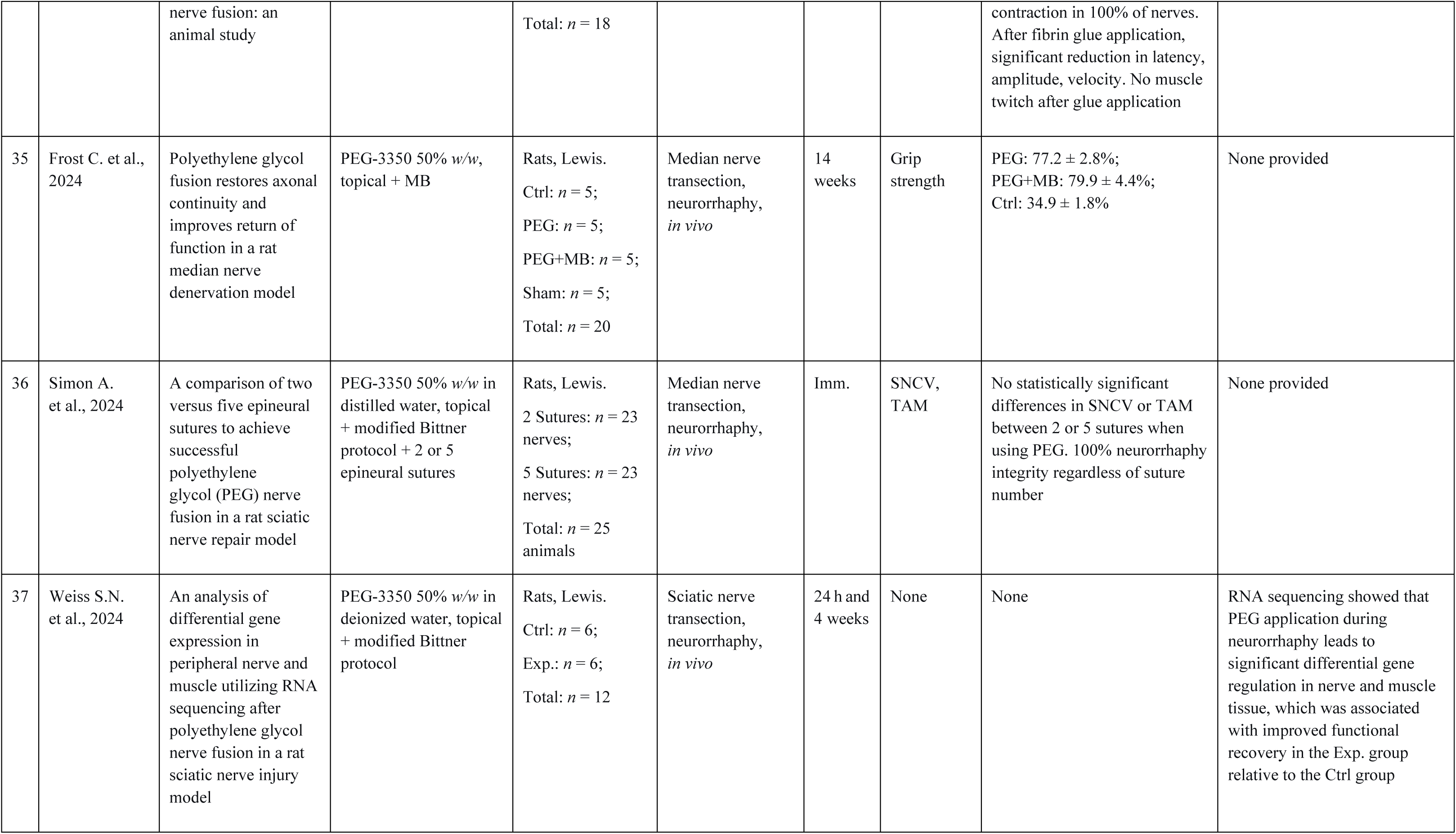
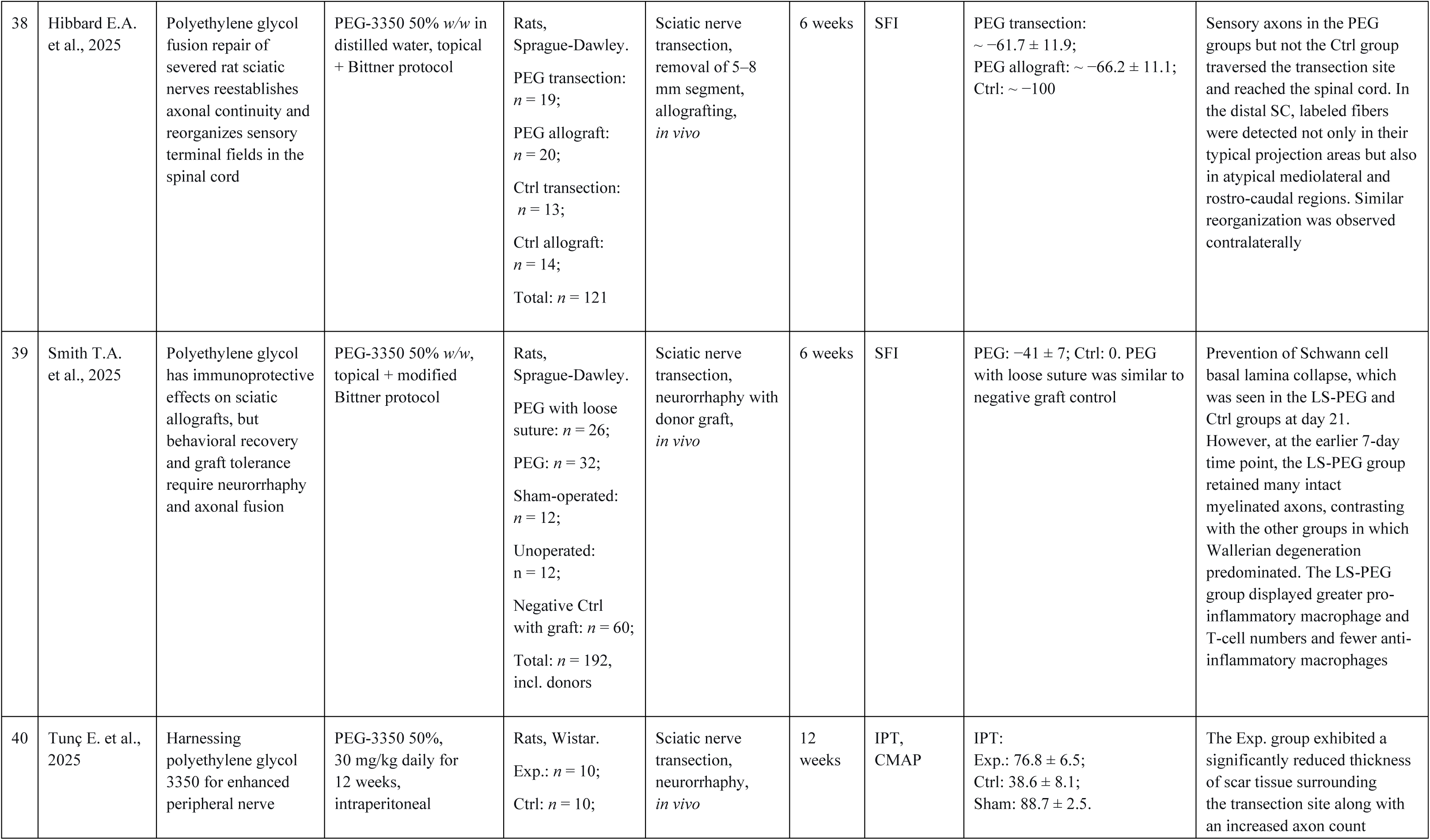
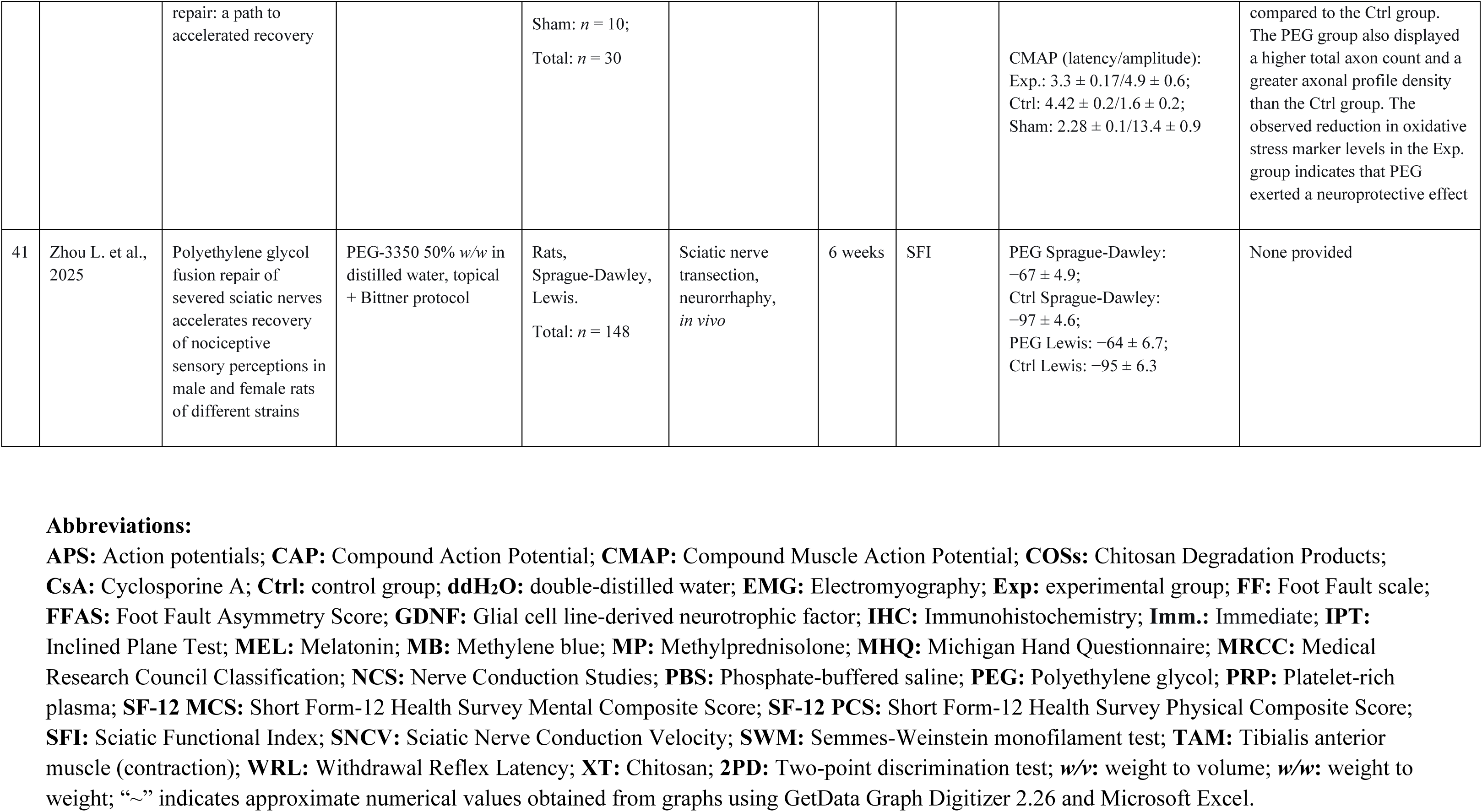
Application for Peripheral Nerve Injury.

### 6.1 Research Subjects (Models)

Studies on SCI predominantly employed animal models. The species distribution was as follows: rats, 14 studies; guinea pigs, 13 studies; dogs, 8 studies; mice, 3 studies; pigs, 3 studies; rabbits, 2 studies; and non-human primates, 1 study. Two studies involved human participants. Among the studies using rats, the strain distribution comprised the following: Sprague-Dawley, 6 studies; Wistar, 5 studies; Lewis, 1 study; and Long-Evans, 1 study; one study used non-pedigreed animals (designated as N/A) (Figure 3).

**Figure 3.**
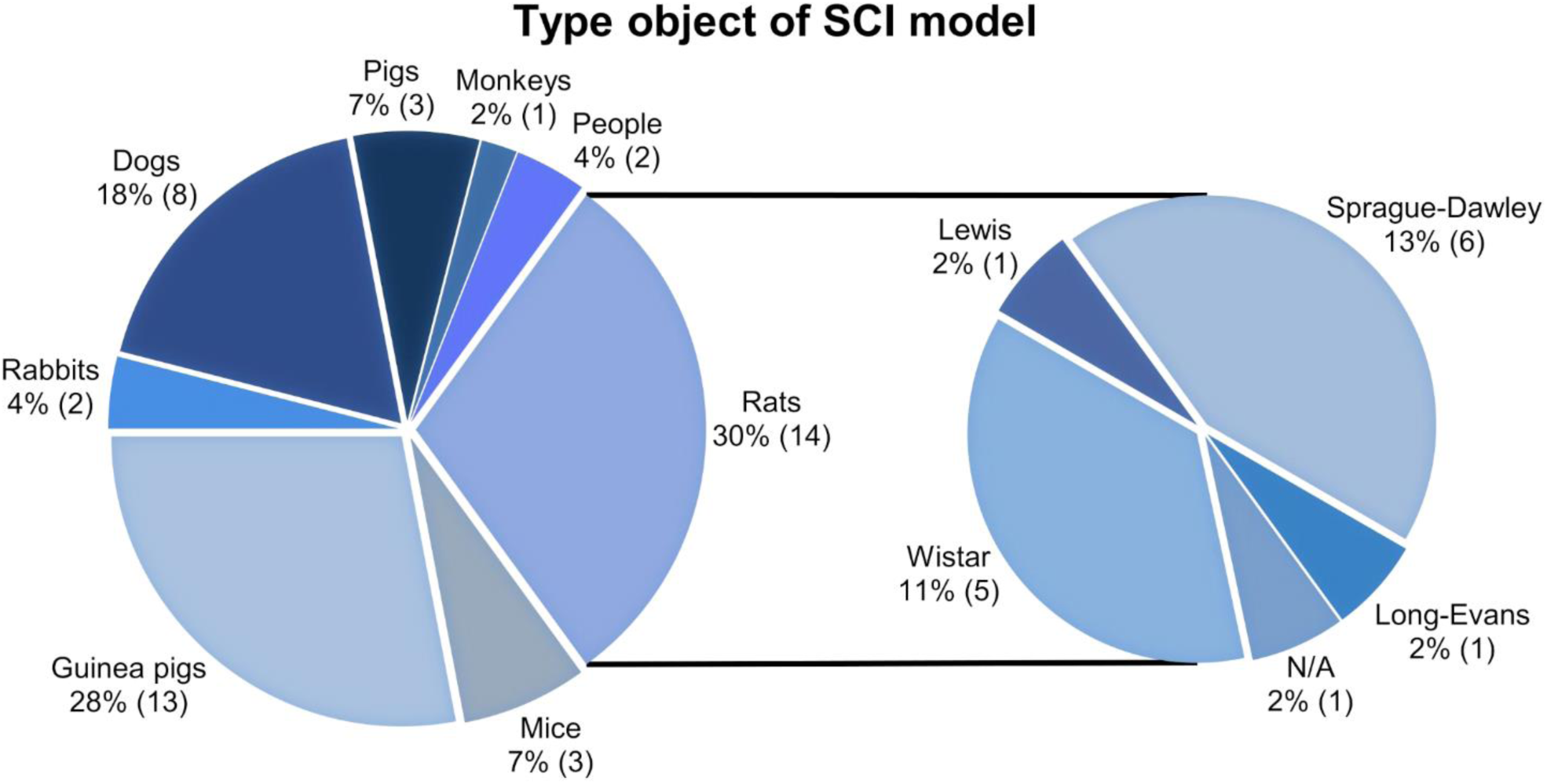
Diagram of the distribution of studies by the type of research object in a model of spinal cord injury. The secondary pie chart shows the distribution of rat studies by breed. The proportion of studies is indicated as a percentage (%), with the number of studies for each research object given in parentheses. N/A: non-pedigree animals or no data available.

Studies on PNI were similarly dominated by preclinical models, with 36 using rats, and one each employing guinea pigs, pigs, and rabbits. Two clinical studies involved human participants. Among the rat studies, 26 involved Sprague-Dawley rats, 4 involved Wistar rats, 5 used Lewis rats, and 2 employed non-pedigreed animals (designated N/A) (Figure 4). One study used both Sprague-Dawley and Lewis rats concurrently. Additionally, the studies by Zhou et al. (2025) and Hibbard et al. (2025) incorporated sex-based subgroup comparisons.

**Figure 4.**
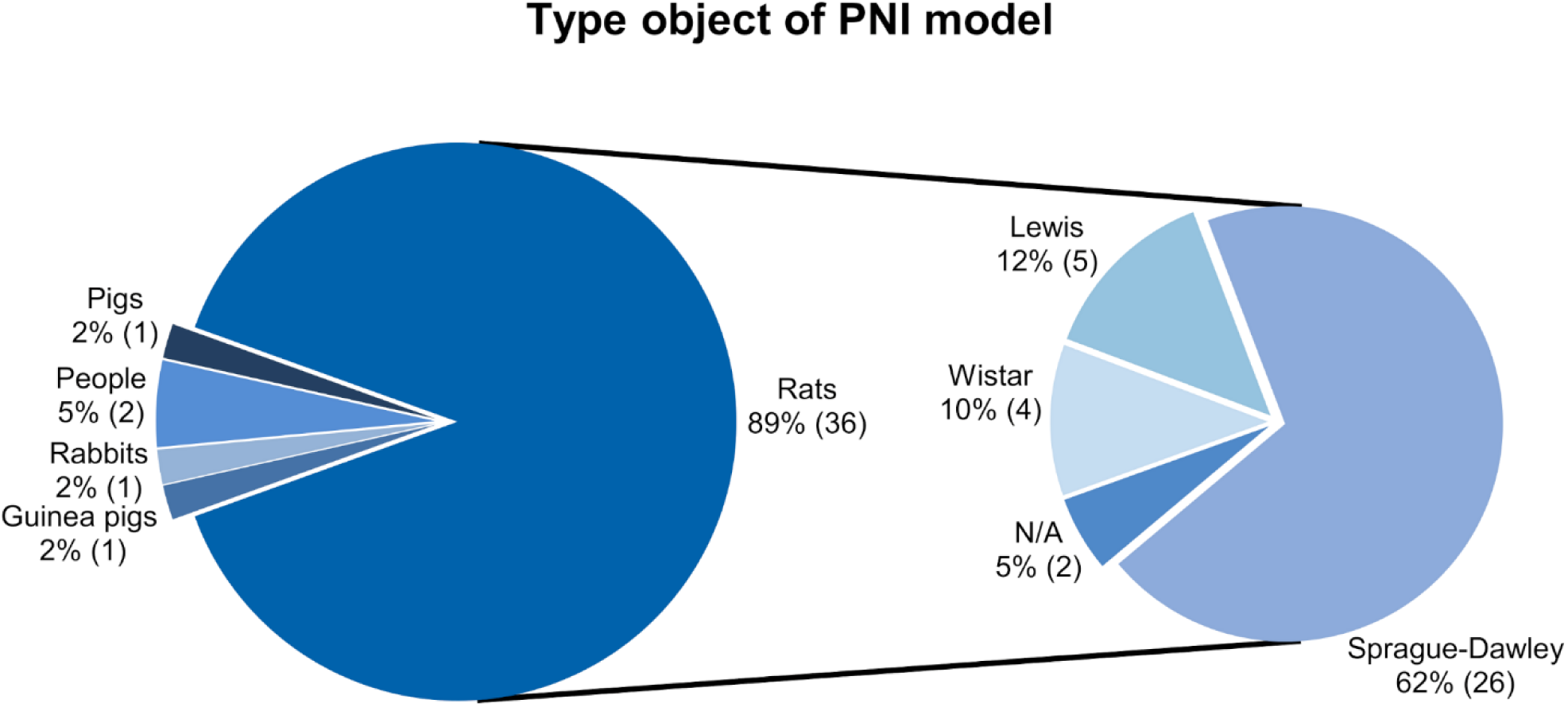
Diagram of the distribution of studies by the type of research object in a model of peripheral nerve injury. The secondary pie chart shows the distribution of rat studies by breed. The proportion of studies is indicated as a percentage (%), with the number of studies for each research object given in parentheses. N/A: non-pedigree animals or no data available.

### 6.2 Spinal Cord Injury and Peripheral Nerve Injury Models

In preclinical SCI research, the most frequently employed model was compression/contusion induced by various weights (24 studies), followed by complete transection (13 studies). Among the latter, Ye et al. (2016) specifically sectioned the dorsal column. Three studies undertook a transplantation of an intact vascular pedicle of hemisected spinal cord (Ren S. et al., 2021; Shen T. et al., 2024; Zhang et al., 2024a). Estrada et al. (2014) resected scar tissue at 5 weeks post-contusion, while Nourbakhsh et al. (2024) performed sural nerve transplantation after excising the contused segment. Additionally, two canine studies employed a clinically relevant model of acute intervertebral disc herniation with laminectomy (Laverty et al., 2004; Olby et al., 2016) (Figure 5A).

**Figure 5.**
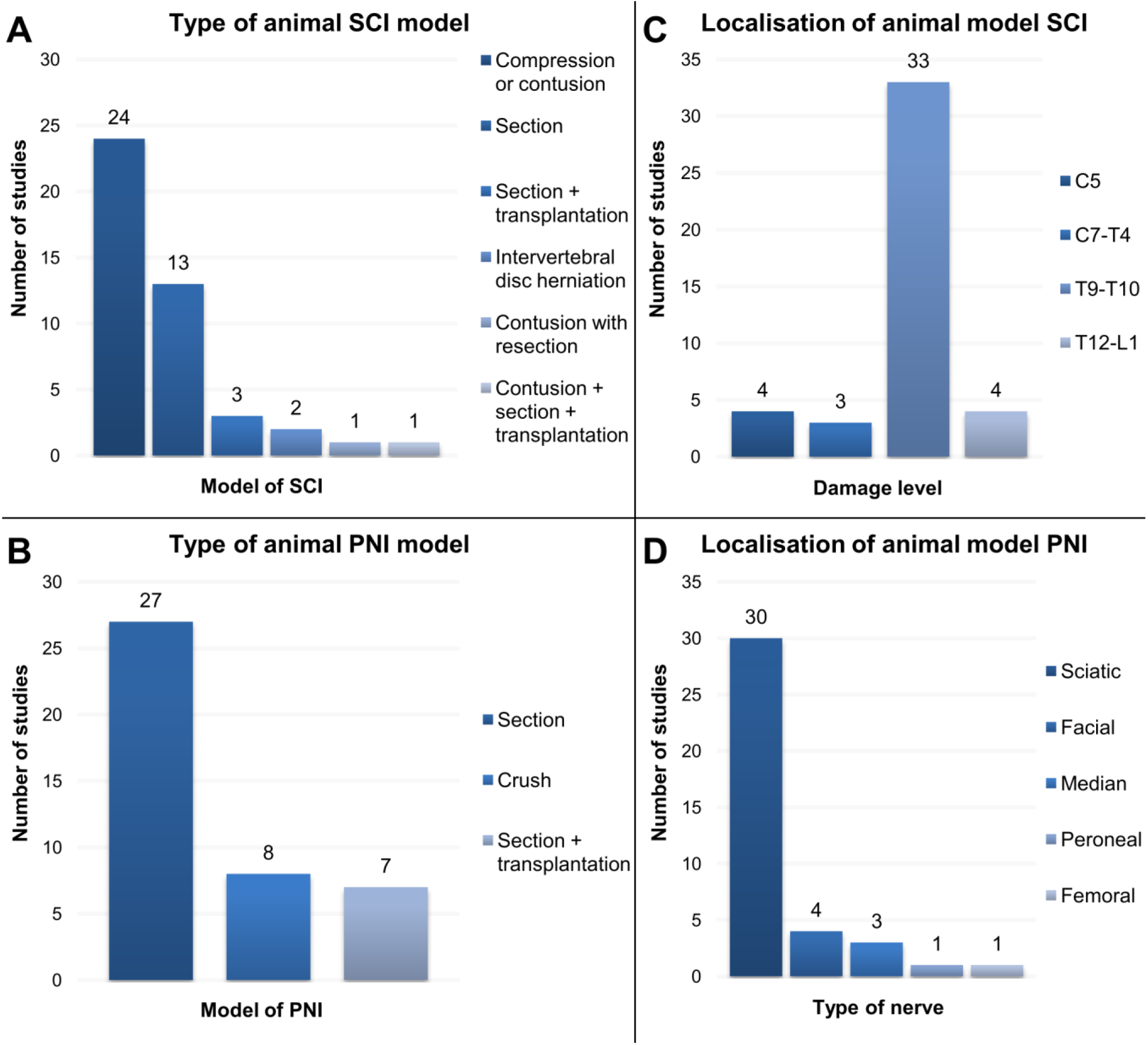
A. - Diagram of the distribution of preclinical studies according to the type of spinal cord injury (SCI) model; **B** - Diagram of the distribution of preclinical studies according to the type of peripheral nerve injury (PNI) model; **C** - Diagram of the distribution of preclinical studies according to the level of spinal cord injury; **D** - Diagram of the distribution of preclinical studies according to the type of damaged nerve.

Clinical human trials exclusively enrolled patients with chronic complete SCI, classified as ASIA A (Ren X. et al., 2022a; Ren X. et al., 2022b).

In preclinical PNI research, nerve transection served as the primary model (27 studies), followed by nerve crush injury (8 studies). Four investigations directly compared both sciatic nerve transection and crush models (Lore et al., 1999; Britt et al., 2010; Bittner et al., 2012; Ghergherehchi et al., 2016). Seven additional studies involved auto- or allograft transplantation to bridge a segmental defect following nerve transection and excision, one of which used an arterial graft (Zhao X. et al., 2019) (Figure 5B).

Clinical studies enrolled patients with acute traumatic peripheral nerve injuries, predominantly affecting distal digital nerve branches (Bamba et al., 2016b; Lopez et al., 2023).

### 6.3 Injury Location

In experimental SCI models, the mid-thoracic region (predominantly T9–T11) was the most commonly targeted site, featuring in 33 studies. Four studies focused on the thoracolumbar junction (T12–L1) (Borgens et al., 2004; Kouhzaei et al., 2013a; Kouhzaei et al., 2013b; Kim C-Y. et al., 2018), while another four involved cervical-level injuries (C5) (Lee J. H. T. et al., 2010; Kim C-Y. et al., 2016; Kim C-Y., 2016; Kim C-Y. et al., 2017). Three studies focused on the cervicothoracic or upper thoracic region (C7–T4) (Ditor et al., 2007; Baptiste et al., 2009; Kouhzaei et al., 2013b) (Figure 5C). Notably, Kouhzaei et al. (2013b) conducted a single experiment spanning three distinct levels (T1–T2, T11, L1–L2). No studies were identified that systematically compared outcomes across different injury levels.

In models of disc herniation-induced SCI, the resulting myelopathy either exhibited a broad anatomical distribution (T3 to L3) (Laverty et al., 2004) or was described as a generalized thoracolumbar localization without precise level specification (Olby et al., 2016).

Clinical studies enrolled patients with chronic, complete paraplegia secondary to SCI at levels T1–T12 (12 patients; Ren X. et al., 2022a) or T4–T10 (8 patients; Ren X. et al., 2022b).

In preclinical PNI research, the sciatic nerve was the predominant model used (30 studies). Other investigated nerves included the facial nerve and its branches (4 studies; Salomone et al., 2018; Brown et al., 2019; Petrov et al., 2022; Şahin et al., 2022), the median nerve (3 studies; Simon et al., 2024; Fisher et al., 2024; Frost et al., 2024), the common peroneal nerve (1 study; Gong et al., 2009), and the femoral nerve (1 study; Robinson and Madison, 2016) (Figure 5D).

Clinical studies involved four patients with acute peripheral nerve injuries, one presenting with a single digital nerve injury and two with triple digital nerve lesions (Bamba et al., 2016b); a separate study reported one patient with a digital nerve injury and another with a median nerve injury (Lopez et al., 2023).

### 6.4 Types of Fusogens

The primary fusogen in most studies was PEG of various MWs and in various combinations with other substances (78 out of 86 studies, of which 42 focused on SCI and 36 on PNI). In SCI models, PEG-600 was used most frequently (*n* = 14), while in PNI models, PEG-3350 was the predominant PEG type employed (*n* = 26). Of all the PEG types, only five were used in both spinal cord and PNI models (PEG-600, PEG-1500, PEG-1800, PEG-2000, and PEG-4000). Only four studies on SCI involved a comparison of PEGs of varying MWs (Borgens and Shi, 2000; Kouhzaei et al., 2013a; Kouhzaei et al., 2013b; Ye et al., 2016). Notably, Shi and Borgens (1999) used four types of PEG but only reported results for PEG-1800. Only one study on PNI compared PEGs of multiple MWs (Lore et al., 1999); however, it was not very informative, as results are only provided for PEG-2000 and PEG-diacrylate.

In most spinal cord studies, the preference for PEG-600 lacked justification. Meanwhile, in nerve fusion research, PEG-3350 was used in 26 out of 41 studies, most often as a 50% solution. As with PEG-600, none of the cited publications offered a rationale for the selection of this PEG MW based on a comparison with others. Interestingly, PEG-3350, which is widely used for nerve fusion, was not applied in any spinal cord-related study. G. D. Bittner, who applied PEG-3350 on nerves, employed PEG-3250, a compound of comparable MW, in his spinal cord experiments, also without explaining this choice (Bittner et al., 2015).

An important characteristic influencing experimental outcomes is the concentration of PEG used (e.g., dilution in distilled water, saline solutions, or phosphate-buffered saline, by mass or volume). In total, 14 studies on SCI models used undiluted, pure PEG (without the addition of other substances) by mass. Interestingly, in PNI models, undiluted PEG (PEG-600) was used in only one study (Zhao X. et al., 2019); all other studies reported using a 50% dilution by mass. A 30% concentration was employed exclusively in SCI studies, administered via several routes: subcutaneous (Borgens and Bohnert, 2001, PEG-2000), intravenous (Laverty et al., 2004, PEG-3500; Ditor et al., 2007, PEG-3500; Baptiste et al., 2009, PEG-2000; Olby et al., 2016, PEG-3500), and subdural (Lee J. et al., 2023, PEG-600). The same concentration was applied in a contusion model involving the excision of the damaged section and its replacement with a sural nerve graft (Nourbakhsh et al., 2024, PEG-600). Furthermore, one study reported a 25% dilution of PEG-600 for intravenous administration (Lebenstein-Gumovski M. et al., 2023b). However, no studies—either in SCI or PNI research—directly compared the effects of different PEG concentrations, except for Kouhzaei et al. (2013b), who evaluated PEG concentrations ranging from 10% to 70% *w*/*w* in serum-free Dulbecco′s Modified Eagle′s Medium (DMEM) in cell culture, and Lore et al. (1999), who presented comparative data for PEG-2000 and PEG-diacrylate.

Chitosan or its degradation products were also applied as a fusogen (4 studies—1 on SCI and 3 on PNI). Notably, pure chitosan was used in only two studies, one employing a model of compressive SCI (Cho et al., 2010) and one a model of facial nerve injury (Şahin et al., 2022). Chitosan degradation products were restricted to two PNI studies (Gong et al., 2009; Jiang et al., 2009). The use of a PEG-chitosan conjugate (Neuro-PEG)—which combines the properties oftwo fusogens—was reported in three research papers relating to a transection model of SCI in various animal models (Lebenstein-Gumovski et al., 2023a; Lebenstein-Gumovski et al., 2023b; Lebenstein-Gumovski et al., 2023c). A different conjugate—azidobenzoyl-chitosan-PEG (Az-C/PEG) (Amoozgar et al., 2012)—was studied solely in peripheral nerve research (Figure 6).

**Figure 6.**
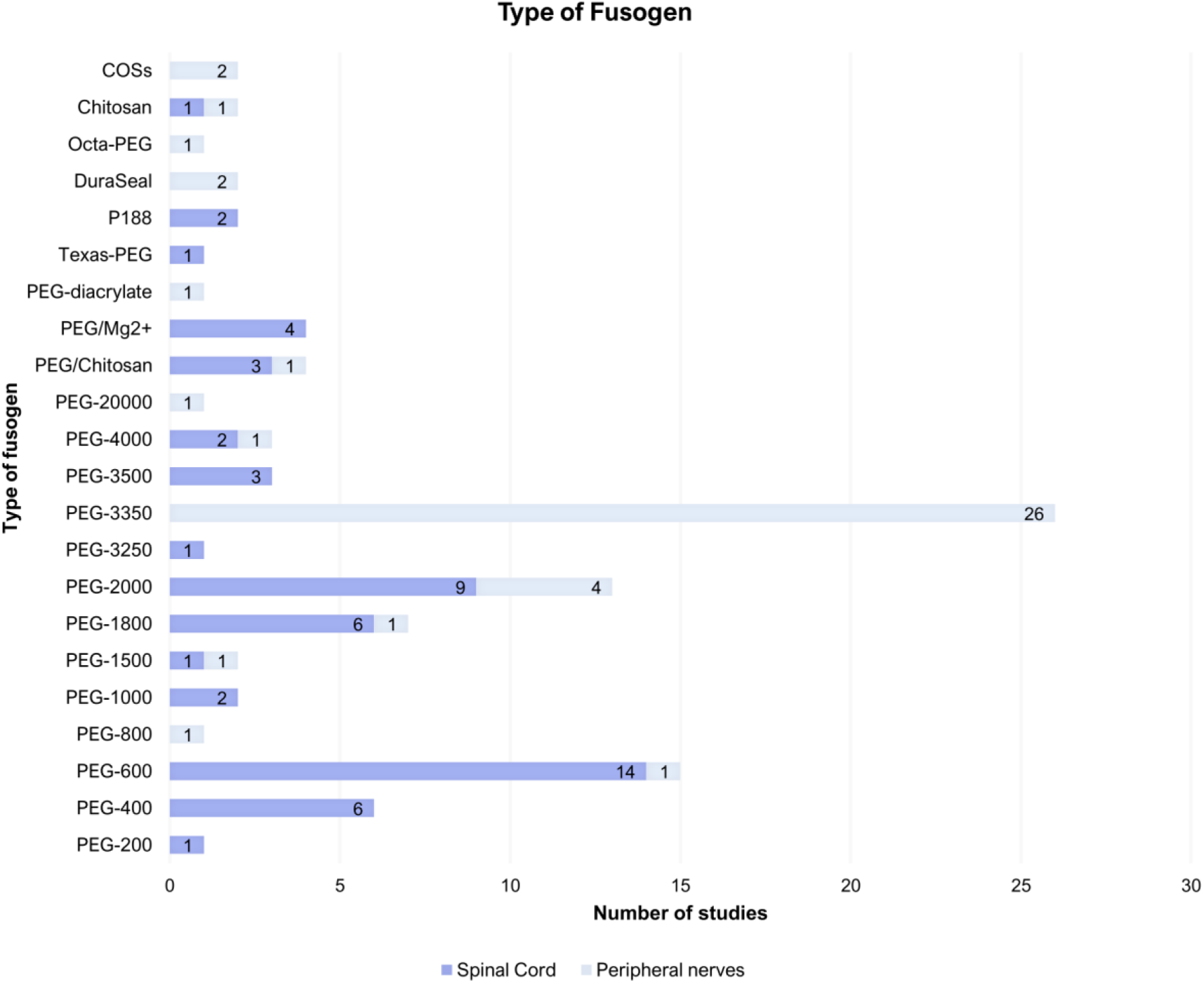
Diagram of the distribution of studies according to the type of fusogen used.

### 6.5 Method of Fusogen Administration

The primary method of fusogen administration in most studies was intraoperative local application at the injury site. Notably, Kim C-Y. et al. (2018) used pegylated graphene nanoribbons (“Texas-PEG”) as a conduit, while Zhao C. et al. (2022) employed a gelatin sponge. The development of a specific device for more effective intraoperative PEG application by Riley et al. (2017) is also of interest.

Exclusively intravenous administration was used in eight studies on SCI (Laverty et al., 2004; Ditor et al., 2007; Baptiste et al., 2009; Kwon et al., 2009; Lee J. H. T. et al., 2010; Olby et al., 2016; Streijger et al., 2016; Huang et al., 2017) and in one study on PNI (Gong et al., 2009).

Combined administration (local and intravenous) was performed in only one study on spinal cord trauma (Lebenstein-Gumovski et al., 2023b). Furthermore, subcutaneous administration was used in three studies involving SCI models (Borgens and Bohnert, 2001; Borgens et al., 2004; Cho et al., 2010). Intraperitoneal administration was employed in two studies on PNI (Jiang et al., 2009; Tunç et al., 2025).

### 6.6 Observation Periods

Follow-up durations in SCI research ranged from 25 minutes to 18 months. In experimental studies, the average observation period was 4–8 weeks in small animals and 6 months in large animals. In studies on PNI, the observation periods ranged from 1 hour to 16 weeks in animals and up to 2 years in humans.

### 6.7 Assessment of Neurological Functions

In animal models of SCI, functional recovery was most commonly assessed using various modifications of the BBB scale, while in PNI models, the SFI was the predominant measure applied. However, other scales, functional tests, and instrumental methods were also employed for evaluating neurological function (Table 1). In the present analysis, we relied on the BBB and SFI scales as they represent the most widely used and accepted tools for assessing functional outcomes in laboratory animals.

### 6.8 Histological Studies

A variety of methods were used for histological assessment, including light and electron microscopy and immunohistochemical and fluorescence analyses, as well as investigations performed on cell cultures (Tables 1 and 2).

### 6.9 Meta-Analysis of Quantitative Data

Of the 75 articles selected for the systematic review, only 20—11 on SCI and 9 on PNI—were included in the quantitative meta-analysis (Tables 3 and 4). The primary criteria for selecting articles for the meta-analysis were the availability of neurological function assessment results based on the BBB and SFI scales for spinal cord and peripheral nerve injuries, respectively. BBB scores were evaluated over 4 weeks, and SFI scores over 6 weeks, to include the largest possible number of studies.

**Table 3.** List and parameters of studies included in the meta-analysis on spinal cord injury.

| № | Author/Year | Fusogen | Subject | Injury model | Injury level | Administration method | Exp Group Size | Ctrl Group Size | Mean at 1 wk (Exp) | Mean at 1 wk (Ctrl) | Mean at 2 wk (Exp) | Mean at 2 wk (Ctrl) | Mean at 3 wk (Exp) | Mean at 3 wk (Ctrl) | Mean at 4 wk (Exp) | Mean at 4 wk (Ctrl) |
| --- | --- | --- | --- | --- | --- | --- | --- | --- | --- | --- | --- | --- | --- | --- | --- | --- |
| 1 | Ditor D.S. et al., 2007 | PEG-3500 | Wistar rat | Compression/contusion | T4 | Intravenous | 11 | 10 | $\sim 2.5 \pm 1.32$ | $\sim 2 \pm 0.94$ | $\sim 4 \pm 1.65$ | $\sim 3.8 \pm 1.26$ | $\sim 5.8 \pm 1.32$ | $\sim 5 \pm 1.58$ | $\sim 6.7 \pm 0.99$ | $\sim 5.7 \pm 1.26$ |
| 2 | Baptiste D.C. et al., 2009 | PEG-2000 | Wistar rat | Compression/contusion | C7-T1 | Intravenous | 11 | 12 | $1.2 \pm 1.82$ | $1.2 \pm 1.83$ | $2.7 \pm 2.08$ | $3.5 \pm 2.49$ | $4.3 \pm 2.42$ | $3.0 \pm 2.35$ | $4.5 \pm 2.18$ | $3.5 \pm 2.56$ |
| 3 | Oda Y. et al., 2014 | PEG-4000 | ICR mice | Section | T10 | Local | 8 | 8 | $\sim 9.2 \pm 1.41$ | $\sim 6 \pm 1.41$ | $\sim 10.3 \pm 1.97$ | $\sim 8.2 \pm 1.69$ | $\sim 11 \pm 1.69$ | $\sim 8.3 \pm 1.69$ | $\sim 12 \pm 1.41$ | $\sim 8.4 \pm 1.69$ |
| 4 | Bittner G.D. et al., 2015 | PEG-3250 | Lewis rat | Compression/contusion | T9-T10 | Local | 6 | 6 | $2.8 \pm 2.22$ | $3.17 \pm 3.06$ | $\sim 9 \pm 1.71$ | $\sim 6 \pm 1.46$ | $\sim 11.3 \pm 1.95$ | $\sim 8 \pm 1.95$ | $\sim 13 \pm 1.95$ | $\sim 9.3 \pm 1.71$ |
| 5 | Ye Y. et al., 2016 | PEG-1500 | C57 mice | Section | T10 | Local | 8 | 8 | $\sim 6 \pm 1.41$ | $\sim 1 \pm 1.41$ | $\sim 11.5 \pm 1.41$ | $\sim 1.5 \pm 1.41$ | $\sim 13.5 \pm 0.84$ | $\sim 1.8 \pm 1.41$ | $\sim 14 \pm 0.84$ | $\sim 2 \pm 1.41$ |
| 6 | Ye Y. et al., 2016 | PEG-4000 | C57 mice | Section | T10 | Local | 8 | 8 | $\sim 3.5 \pm 0.28$ | $\sim 1 \pm 1.41$ | $\sim 6.5 \pm 1.41$ | $\sim 1.5 \pm 1.41$ | $\sim 9.8 \pm 1.41$ | $\sim 1.8 \pm 1.41$ | $\sim 10 \pm 1.13$ | $\sim 2 \pm 1.41$ |
| 7 | Ren S. et al., 2017 | PEG-600 | Rat N/A | Section | T10 | Local | 9 | 5 | $\sim 6 \pm 3$ | $\sim 2.5 \pm 1.11$ | $\sim 9.5 \pm 2.7$ | $\sim 3.2 \pm 1.11$ | $\sim 11 \pm 2.7$ | $\sim 4 \pm 1.11$ | $\sim 12 \pm 3$ | $\sim 4.4 \pm 2.23$ |
| 8 | Kim C-Y. et al., 2018 | Texas-PEG | Rat SD | Section | L1 | Local | 10 | 10 | $\sim 2 \pm 0.94$ | $\sim 1.8 \pm 0.94$ | $\sim 4.2 \pm 1.89$ | $\sim 2.6 \pm 1.26$ | $\sim 5.6 \pm 2.21$ | $\sim 3 \pm 1.26$ | $\sim 7.2 \pm 2.52$ | $\sim 3.5 \pm 1.58$ |
| 9 | Lee J. et al., 2023 | PEG-600 | Rat SD | Compression/contusion | T10 | Intravenous | 6 | 6 | $\sim 7.5 \pm 2.4$ | $\sim 4.5 \pm 1.71$ | $\sim 12.5 \pm 2.2$ | $\sim 9 \pm 1.95$ | $\sim 13 \pm 2.44$ | $\sim 9.5 \pm 1.95$ | $\sim 13 \pm 2.44$ | $\sim 10 \pm 1.95$ |
| 10 | Lebenstein-Gumovski M.V. et al., 2023a | Neuro-PEG | Rabbit | Section | T9 | Local | 10 | 10 | 11 ± 1.58 | 1 ± 0.63 | 16 ± 1.58 | 1.5 ± 1.58 | 18 ± 1.58 | 2 ± 0.94 | 20 ± 1.58 | 2.5 ± 1.58 |
| 11 | Lebenstein-Gumovski M.V. et al., 2023b | Neuro-PEG | Pig | Section | T9 | Local + intravenous | 3 | 2 | 7.3 ± 1.73 | 1 ± 0.7 | 10 ± 1.21 | 1.5 ± 0.7 | 14 ± 1.73 | 1.5 ± 0.28 | 17 ± 1.73 | 2 ± 0.28 |
| 12 | Lebenstein-Gumovski M.V. et al., 2023c | Neuro-PEG | Rabbit | Section | T9 | Local | 10 | 10 | 8 ± 1.26 | 1 ± 0.63 | 12 ± 1.89 | 2 ± 0.94 | 14 ± 0.06 | 2 ± 0.94 | 15 ± 1.58 | 2 ± 1.89 |

Data are presented as means with standard deviation (SD). The SD was calculated from the standard error of the mean (SEM) using the formula described in Methods.
“~” indicates approximate numerical values obtained from graphs using GetData Graph Digitizer 2.26 and Microsoft Excel

**Table 4.** List and parameters of studies included in the meta-analysis on peripheral nerve injury.

| № | Author/Year | Fusogen | Subject | Injury model | Exp Group Size | Ctrl Group Size | 1-Week Mean (Exp) | 1-Week Mean (Ctrl) | 2-Week Mean (Exp) | 2-Week Mean (Ctrl) | 3-Week Mean (Exp) | 3-Week Mean (Ctrl) | 4-Week Mean (Exp) | 4-Week Mean (Ctrl) | 5-Week Mean (Exp) | 5-Week Mean (Ctrl) | 6-Week Mean (Exp) | 6-Week Mean (Ctrl) |
| --- | --- | --- | --- | --- | --- | --- | --- | --- | --- | --- | --- | --- | --- | --- | --- | --- | --- | --- |
| 1 | Britt. J.K. et al., 2010 | PEG-2000 | SD rat | Crush | 13 | 18 | $\sim -90 \pm 9.01$ | $\sim -95 \pm 12.72$ | $\sim -85 \pm 10.81$ | $\sim -90 \pm 16.97$ | $\sim -40 \pm 14.42$ | $\sim -50 \pm 16.97$ | $\sim -20 \pm 10.81$ | $\sim -25 \pm 16.97$ | $\sim -20 \pm 9.01$ | $\sim -22 \pm 16.97$ | $\sim -15 \pm 9.01$ | $\sim -20 \pm 14.84$ |
| 2 | Bittner G.D. et al., 2012 | PEG-3350 | SD rat | Crush | 20 | 20 | $\sim -95 \pm 8.04$ | $\sim -96 \pm 10.28$ | $\sim -93 \pm 12.52$ | $\sim -95 \pm 13.41$ | $\sim -33.7 \pm 8.04$ | $\sim -47.9 \pm 17.88$ | $\sim -23 \pm 8.04$ | $\sim -21.5 \pm 8.04$ | $\sim -20.9 \pm 11.62$ | $\sim -20.1 \pm 8.49$ | $\sim -19.1 \pm 8.94$ | $\sim -17.6 \pm 9.39$ |
| 3 | Bittner G.D. et al., 2012 | PEG-3350 | SD rat | Cut | 4 | 10 | $\sim -81.7 \pm 5.6$ | $\sim -88.7 \pm 6.95$ | $\sim -73.1 \pm 6.4$ | $\sim -87.4 \pm 9.8$ | $\sim -61.5 \pm 3.2$ | $\sim -89.1 \pm 6$ | $\sim -55.8 \pm 5.8$ | $\sim -87.8 \pm 7.9$ | $\sim -50.9 \pm 3$ | $\sim -87.4 \pm 9.17$ | $\sim -48.7 \pm 2.4$ | $\sim -84.2 \pm 10.75$ |
| 4 | Ghergherehchi C.L. et al., 2016 | PEG-3350 | SD rat | Cut | 5 | 5 | $\sim -95 \pm 11.18$ | $\sim -105 \pm 11.18$ | $\sim -70 \pm 11.18$ | $\sim -105 \pm 17.88$ | $\sim -45 \pm 15.62$ | $\sim -105 \pm 0.22$ | $\sim -40 \pm 15.65$ | $\sim -95 \pm 11.18$ | $\sim -40 \pm 11.18$ | $\sim -92 \pm 6.7$ | $\sim -35 \pm 11.18$ | $\sim -96 \pm 6.7$ |
| 5 | Ghergherehchi C.L. et al., 2016 | PEG-3350 | SD rat | Crush | 4 | 5 | $\sim -103 \pm 12$ | $\sim -105 \pm 11.18$ | $\sim -90 \pm 12$ | $\sim -92 \pm 11.18$ | $\sim -85 \pm 0.22$ | $\sim -85 \pm 11.18$ | $\sim -35 \pm 6.7$ | $\sim -37 \pm 4.47$ | $\sim -25 \pm 2.23$ | $\sim -25 \pm 8.94$ | $\sim -24 \pm 10$ | $\sim -25 \pm 11.18$ |
| 6 | Riley D.C. et al., 2017 | PEG-3350 | SD rat | Cut | 6 | 6 | $\sim -103 \pm 7.34$ | $\sim -78 \pm 4.89$ | $\sim -75 \pm 2.44$ | $\sim -105 \pm 4.89$ | $\sim -77 \pm 12.24$ | $\sim -100 \pm 7.34$ | $\sim -72 \pm 4.89$ | $\sim -95 \pm 7.34$ | $\sim -62 \pm 4.89$ | $\sim -97 \pm 12.24$ | $\sim -60 \pm 4.89$ | $\sim -92 \pm 12.24$ |
| 7 | Mikesh M. et al., 2018a | PEG-3350 | SD rat | Cut | 6 | 4 | $\sim -97 \pm 7.34$ | $\sim -95 \pm 6$ | $\sim -80 \pm 7.34$ | $\sim -90 \pm 4$ | $\sim -70 \pm 6$ | $\sim -87 \pm 6$ | $\sim -38 \pm 7.34$ | $\sim -90 \pm 10$ | $\sim -40 \pm 4$ | $\sim -87 \pm 6$ | $\sim -29 \pm 7.34$ | $\sim -85 \pm 4$ |
| 8 | Lin Y.F. et al., 2019 | PEG-800 | SD rat | Cut | 30 | 30 | $\sim -89.83 \pm 0.73$ | $\sim -98.29 \pm 0.68$ | $\sim -62.55 \pm 0.38$ | $\sim -78.15 \pm 1.03$ | $\sim -53.28 \pm 2.39$ | $\sim -73.16 \pm 2.19$ | $\sim -44.01 \pm 4.40$ | $\sim -68.17 \pm 3.35$ | $\sim -40.71 \pm 3.8$ | $\sim -62.43 \pm 2.55$ | $\sim -37.41 \pm 3.20$ | $\sim -56.69 \pm 1.76$ |
| 9 | Zhao X. et al., 2019 | PEG-600 | SD rat | Cut | 36 | 36 | $\sim -92.5 \pm 12$ | $\sim -94 \pm 12$ | $\sim -90 \pm 12$ | $\sim -92 \pm 12$ | $\sim -80 \pm 12$ | $\sim -84 \pm 12$ | $\sim -70 \pm 12$ | $\sim -76 \pm 12$ | $\sim -66 \pm 12$ | $\sim -73 \pm 12$ | $\sim -63 \pm 12$ | $\sim -71 \pm 12$ |
| 10 | Ghergherehchi C.L. et al., 2021 | PEG-3350 | SD rat | Cut | 10 | 18 | $\sim -95 \pm 5$ | $\sim -96 \pm 6$ | $\sim -87 \pm 5$ | $\sim -92 \pm 5$ | $\sim -84 \pm 5$ | $\sim -90 \pm 5$ | $\sim -73 \pm 2.39$ | $\sim -89 \pm 5$ | $\sim -68 \pm 5$ | $\sim -88 \pm 5$ | $\sim -63 \pm 5$ | $\sim -86 \pm 5$ |
| 11 | Zhou L. et al.,<br>2025 | PEG-3350 | SD rat | Cut | 35 | 21 | ~ -90±<br>11.83 | ~ -100±<br>9.16 | ~ -93±<br>11.83 | ~ -91±<br>9.16 | ~ -95±<br>11.83 | ~ -94±<br>9.16 | ~ -91±<br>11.83 | ~ -94±<br>9.16 | ~ -74±<br>14.79 | ~ -97±<br>13.74 | ~ -67±<br>23.07 | ~ -97±<br>21.07 |
| 12 | Zhou L. et al.,<br>2025 | PEG-3350 | Lewis<br>rat | Cut | 20 | 13 | ~ -90±<br>8.94 | ~ -87±<br>7.21 | ~ -86±<br>8.94 | ~ -90±<br>7.21 | ~ -85±<br>8.94 | ~ -87±<br>7.21 | ~ -82±<br>13.41 | ~ -88±<br>10.81 | ~ -76±<br>15.65 | ~ -92±<br>10.81 | ~ -64±<br>25.49 | ~ -95±<br>22.71 |

Data are presented as means with standard deviation (SD). The SD was calculated from the standard error of the mean (SEM) using the formula described in Methods.
“~” indicates approximate numerical values obtained from graphs using GetData Graph Digitizer 2.26 and Microsoft Excel.

Among the studies selected for the systematic review (Tables 1 and 2), the BBB scale was used to assess functional outcomes in 16 articles. However, five studies were excluded: four (Liu et al., 2018; Ren S. et al., 2019; Ren S. et al., 2021; and Shen T. et al., 2024) because they reported medians without confidence intervals, and one (Zhang et al., 2024b) because the results accounted for the use of an additional substance, tacrolimus. The study by Ye et al. (2016) was included twice, as it compared two types of PEG—PEG-1500 and PEG-4000—in different groups.

The SFI scale was used for assessing functional outcomes in 20 articles. Six of these—Jiang et al. (2009), Sexton et al. (2012), Bamba et al. (2018), Bu et al. (2020), Pascal et al. (2020) and Acharya et al. (2023)—were excluded for diverse reasons (observation periods shorter than 8 weeks [ranging from 3 days to 3 weeks], single-time SFI assessments without dynamic monitoring, the absence of a group using pure fusogen without other drugs). Studies by Riley et al. (2015), Mikesh et al. (2018a), Mikesh et al. (2018b), Ghergherehchi et al. (2019), Smith et al. (2025) and Hibbard et al. (2025) were also excluded, as results were only provided for groups with grafts. Studies by Bittner et al. (2012), and Ghergherehchi et al. (2021) were included twice, as they compared two types of PNI models—crush and transection—in different groups. The study by Zhou et al. (2025) was also included twice, as it compared two breeds of laboratory rats—Sprague-Dawley and Lewis—in different groups.

### 6.10 Overall Analysis of Fusogen Effects

Four weeks post-SCI, animals administered fusogens exhibited significantly improved neurological function, as measured by the BBB scale, compared to controls (SMD = 3.30, 95% CI: 2.06 to 4.54; *p*< 0.00001), reflecting a robust therapeutic benefit (Figure 7A).

**Figure 7.**
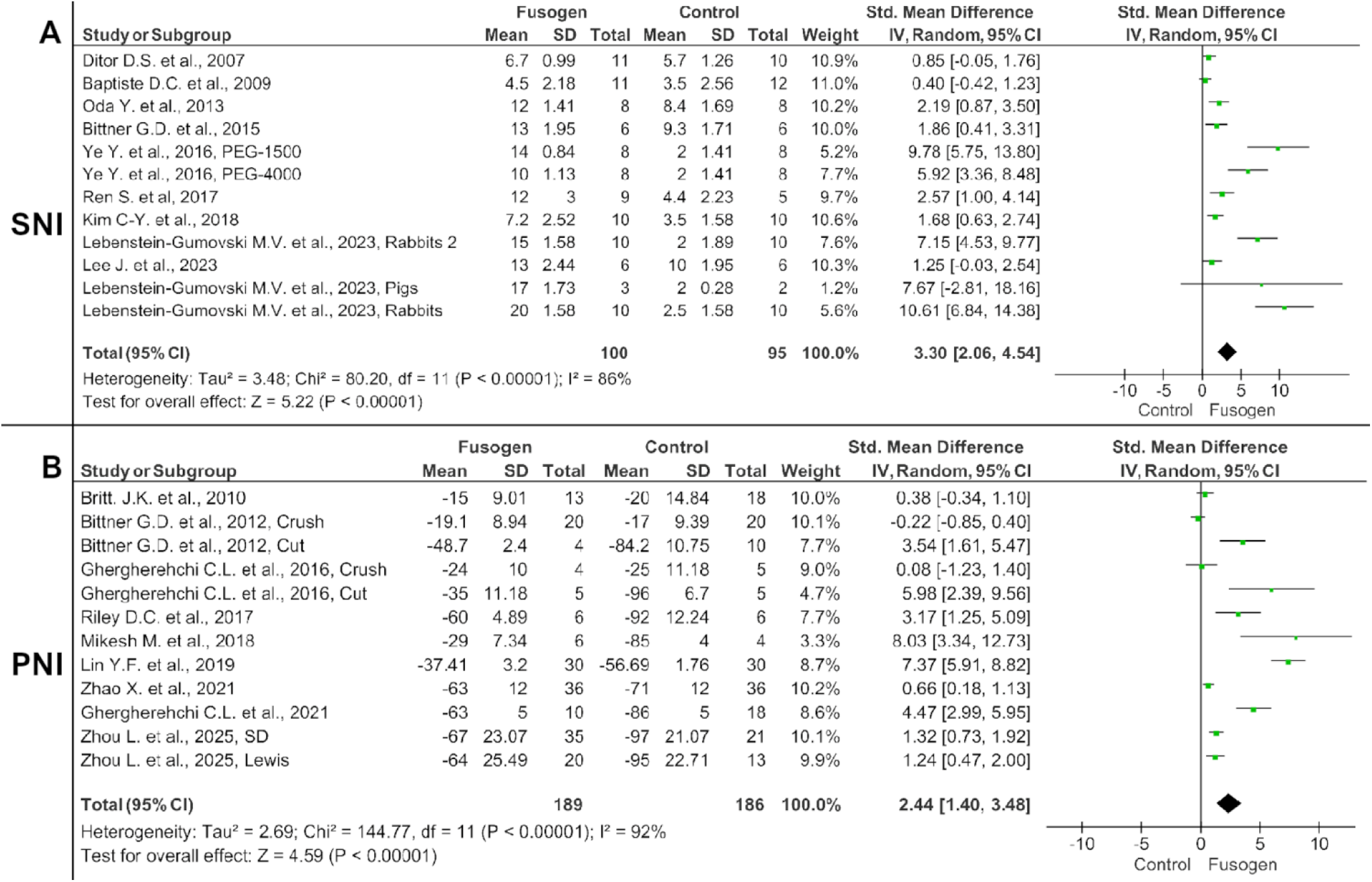
A. - Forest plot and meta-analysis of Basso-Beattie-Bresnahan (BBB) score results at 4 weeks after spinal cord injury; **B** - Forest plot and meta-analysis of Sciatic Functional Index (SFI) score results at 6 weeks after peripheral nerve injury.

Efficacy was similarly demonstrated following PNI. By week 6, fusogen-treated animals demonstrated significant functional recovery on the SFI scale relative to controls (SMD = 2.44, 95% CI: 1.40 to 3.48; *p*< 0.00001) (Figure 7B).

Notably, we observed that there was considerable heterogeneity across the studies for both spinal cord (*χ*² = 80.20, *I*² = 86%; Figure 7A) and peripheral nerve injuries (*χ*² = 144.77, *I*² = 92%; Figure 7B). This variability in outcomes is likely attributable to differences in model organisms, injury paradigms, fusogen types, administration routes, functional assessment methodologies, and cohort sizes. The significant heterogeneity was further corroborated by funnel plot analysis (Figure 8).

**Figure 8.**
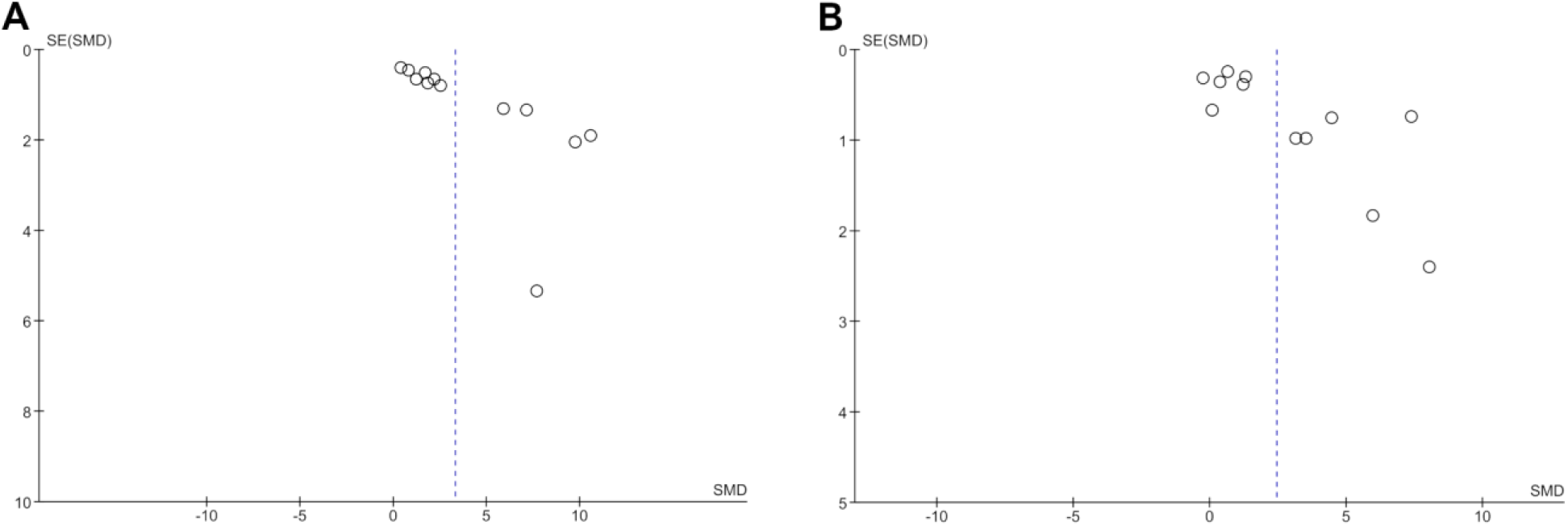
Funnel plot. **A** - The distribution of studies included in the meta-analysis of spinal cord injury; **B** - The distribution of studies included in the meta-analysis of peripheral nerve injury. Significant heterogeneity is observed, likely due to differences in study characteristics.

### 6.11 Subgroup Analysis

Subgroup analysis of SCI studies identified significant heterogeneity between different injury models (transection *vs*. compression/contusion; *χ*² = 15.91, *I*² = 93.7%, *p*< 0.00001). The treatment effect was substantially more pronounced in the transection model (SMD = 5.23, 95% CI: 3.17 to 7.28; *p*< 0.00001) than in the contusion or compression models (SMD = 0.89, 95% CI: 0.34 to 1.44; *p* = 0.002) (Figure 9A).

**Figure 9.**
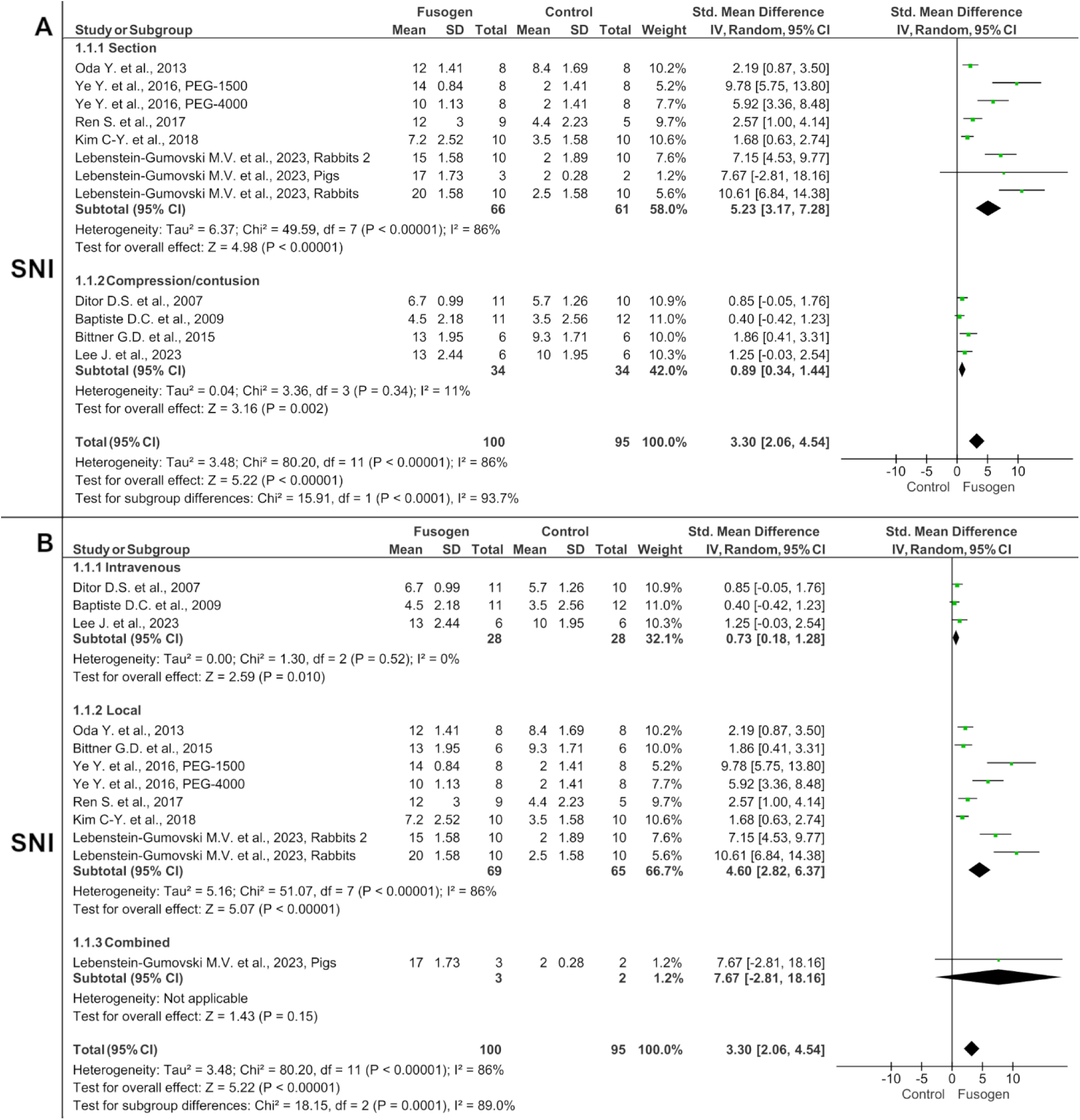
A. - Forest plot and subgroup meta-analysis of Basso-Beattie-Bresnahan (BBB) score results at 4 weeks after spinal cord injury. Comparison by injury model – transection or compression/contusion spinal cord injury; **B** - Forest plot and subgroup meta-analysis of Basso-Beattie-Bresnahan (BBB) score results at 4 weeks after spinal cord injury. Comparison of fusogens administration methods – intravenous, local, and combined.

Furthermore, subgroup analysis based on the route of fusogen administration (intravenous, local, or combined) also revealed significant differences (*χ*² = 18.15, *I*² = 89%, *p* = 0.0001). Local administration of PEG yielded a substantially greater treatment effect (SMD = 4.60, 95% CI: 2.82 to 6.37; *p*< 0.00001) than intravenous delivery (SMD = 0.73, 95% CI: 0.18 to 1.28; *p* = 0.52). Notably, a combined administration approach was associated with the highest effect size (SMD = 7.67, 95% CI: −2.81 to 18.16). However, these data should be interpreted with caution, as this strategy was evaluated in only one study with a small sample size (3 experimental and 2 control animals) (Figure 9B).

In PNI studies, subgroup analysis also showed significant divergence between injury models (crush vs. transection; χ² = 21.59, I² = 95.7%, p< 0.00001). The therapeutic effect was more marked following nerve transection (SMD = 3.48, 95% CI: 2.10 to 4.86; p< 0.00001) than after crush injury (SMD = 0.04, 95% CI: −0.40 to 0.48; p = 0.46) (Figure 10A).

**Figure 10.**
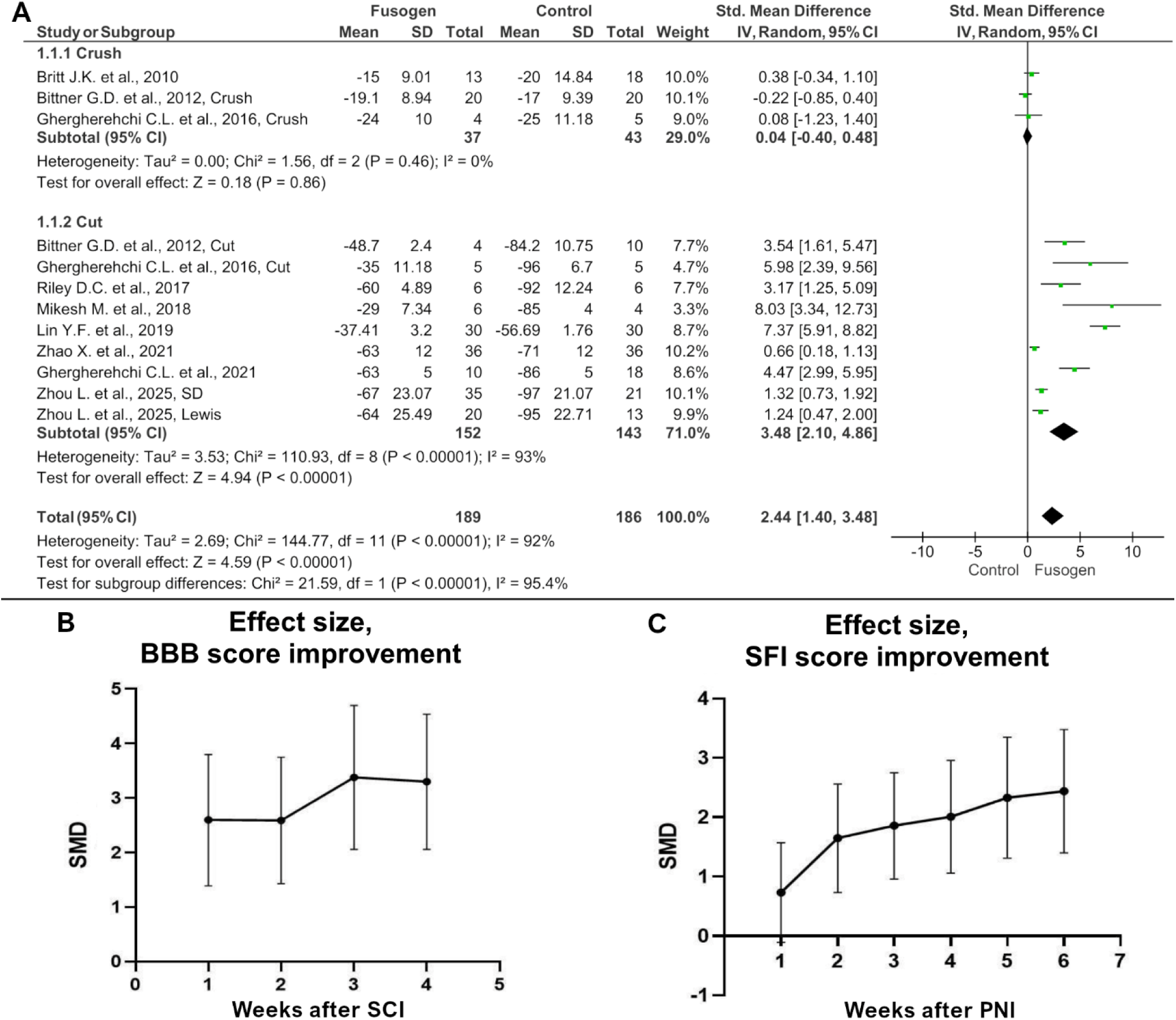
Forest plot and subgroup meta-analysis of Sciatic Functional Index (SFI) scores at 6 weeks after peripheral nerve injury **(A)**. Comparison by injury model – crush or transection of peripheral nerves; Effect sizes (SMD with error bars, calculated as 95% CI) on the Basso-Beattie-Bresnahan (BBB) **(B)** and Sciatic Functional Index (SFI) **(C)**; scales over time (by week). Effect size values for the BBB scale are shown at 4 weeks, and for the SFI scale at 6 weeks.

### 6.12 Sensitivity Analysis

Sensitivity analysis was conducted by sequentially excluding studies with limited cohort sizes or by removing individual studies one at a time. Moderate to high heterogeneity persisted across all exclusion methods. The improvements in BBB and SFI scores, however, remained largely consistent, except for the first week—a deviation likely attributable to the confounding effects of the immediate postoperative period. These results indicate that the beneficial effect of fusogen treatment on motor function recovery following both spinal cord and peripheral nerve injuries was robust and stable.

### 6.13 Limitations of the Meta-Analysis

The findings of this meta-analysis should be interpreted as approximate owing to constraints inherent to the data extraction methodology, the limited number of eligible studies, and their small sample sizes. Primary studies frequently reported outcomes graphically without providing raw numerical values, necessitating the use of third-party software for data digitization, a process that may have introduced measurement inaccuracies. The small number of included studies precluded further subgroup analyses that could have elucidated the sources of heterogeneity. For SCI research, we were unable to stratify results by animal species, injury level, or specific PEG variant. Similarly, for PNI studies, it was not feasible to undertake subgroup analyses based on animal species, fusogen type, administration route, or injury location. This is particularly relevant for non-sciatic nerve injuries, as functional recovery was assessed using the SFI, which is specific to sciatic nerve deficits.

Future investigations designed to compare fusogen effects across these variables are needed to validate these findings and definitively identify the factors modulating therapeutic outcomes. Despite these limitations, the present meta-analysis reveals a compelling positive trend for fusogen therapy in neural repair, as evidenced by the temporal progression of aggregate effect sizes (Figure 10B).

## 7 Discussion

### 7.1 Methodology of Axonal Fusion

The classic Bittner protocol outlines specific steps for axon preparation, both before and after fusogen application. The author of the protocol based it on a calcium-dependent mechanism of axon sealing (Bittner et al., 2016).

For PEG-mediated fusion to occur, the axonal tube must be kept open and gaping (Figure 11). To achieve this, transected axons are “opened” by exposure to a hypotonic calcium-free solution of a specific osmolarity. This increases axoplasmic volume and opens the axonal ends by removing intracellular membrane-bound vesicles/organelles (Figure 12 A, B).

**Figure 11.**
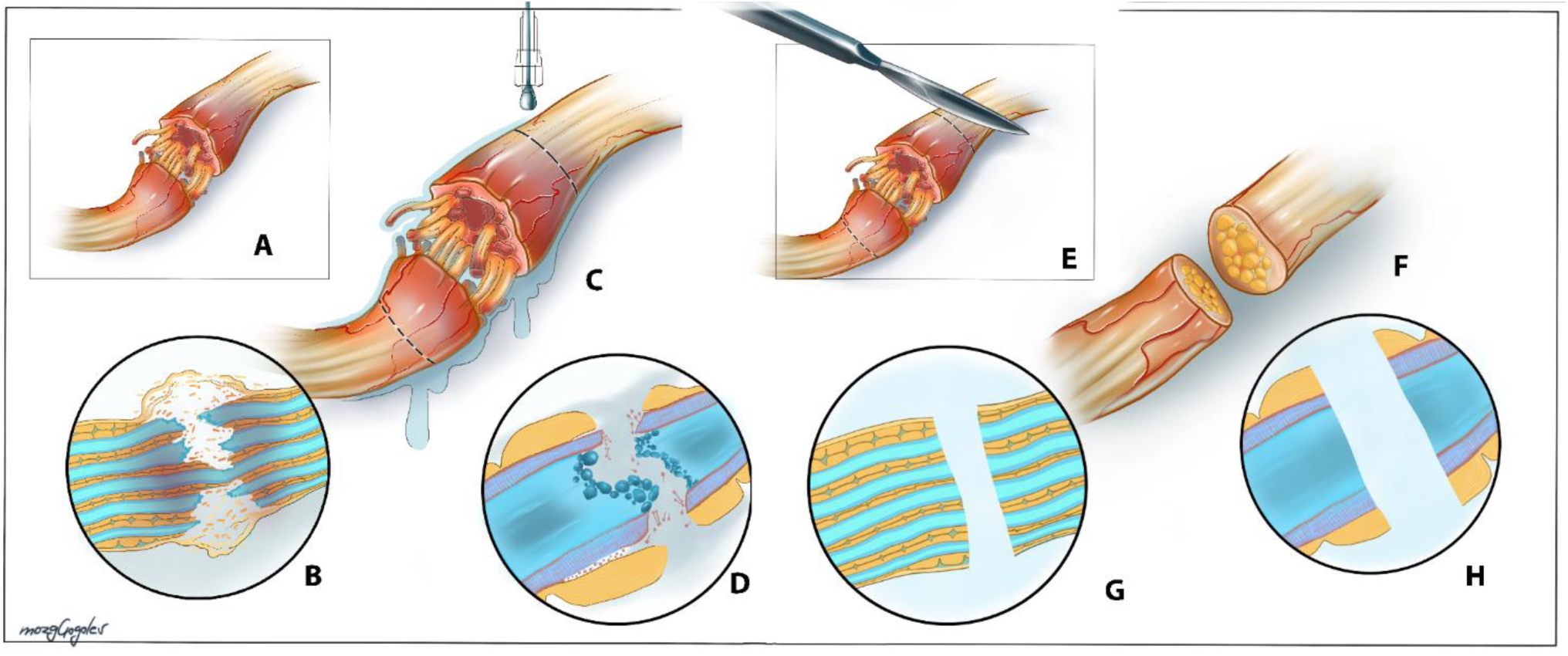
Preparation of nerve ends for neurorrhaphy using a fusogenic protocol. **A** - injured nerve; **B** - schematic representation of axons in an injured nerve. Some axons are destroyed, others are preserved. Vesicles and cell debris are formed; **C** - irrigation with a hypotonic calcium-free solution; **D** - schematic representation of an individual injured axon with a damaged myelin sheath and self-sealing of the ends due to calcium-bound vesicles; **E** - excision of nerve ends within viable tissue in a hypotonic solution; **F** - excised nerve ends; **G** - schematic representation of axons after excision; **H** - diagram of an individual axon. Components that prevent fusion are absent.

**Figure 12.**
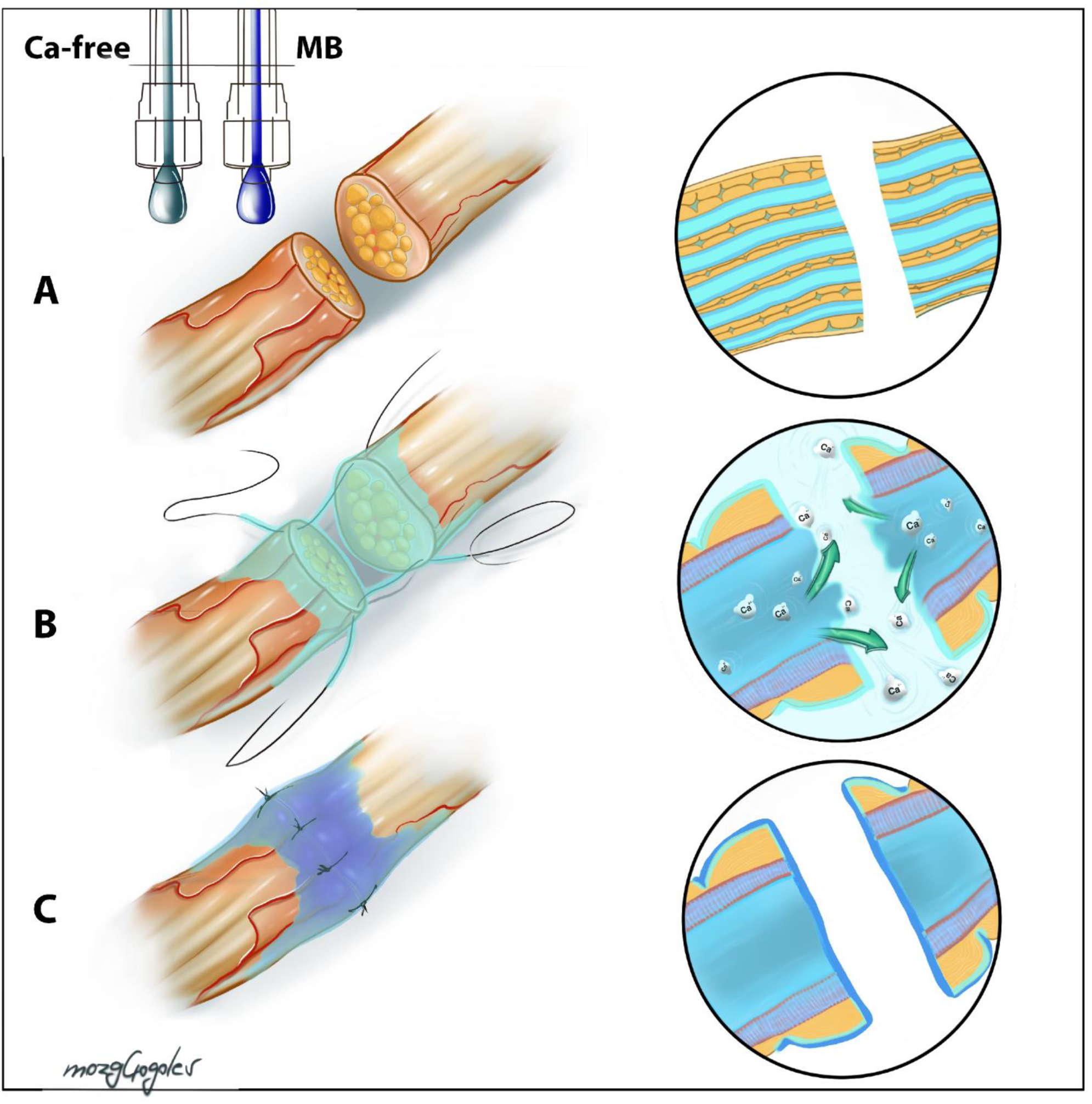
Preparing the excised nerve ends for fusion. **A** - Nerve ends. Macroscopic view is on the left, microscopic view is in the circle on the right; **B** - Copious application of a hypotonic calcium-free solution and placement of the first sutures. Macroscopic view is on the left, microscopic view is in the circle on the right. Active release of axoplasm with increased levels of intracellular calcium, proinflammatory aggregates, and calcium-bound vesicles occurs; **C** - After all sutures are placed, the suture line is treated with 1% methylene blue (MB) solution to achieve an antioxidant cytoprotective effect and membrane stabilization. Macroscopic view is on the left, microscopic view is in the circle on the right.

In the second step, methylene blue solution is applied, which prevents the formation of intracellular vesicles that promote the sealing of the cut axonal ends (Figure 12 C).

Subsequently, nerve microsuture is performed with maximum precision, ensuring end-to-end approximation without tension. This allows the applied solutions to penetrate the transection site into the axons and achieve repair.

The next step involves irrigating the anastomosis site with a 50% PEG solution. According to G. D. Bittner, PEG removes bound cellular water, leading to the fusion of closely apposed open axonal membranes by folding and merging of membrane lipids, and sealing small holes in the plasmalemma (Bittner et al., 2016) (Figure 13 A-F).

**Figure 13.**
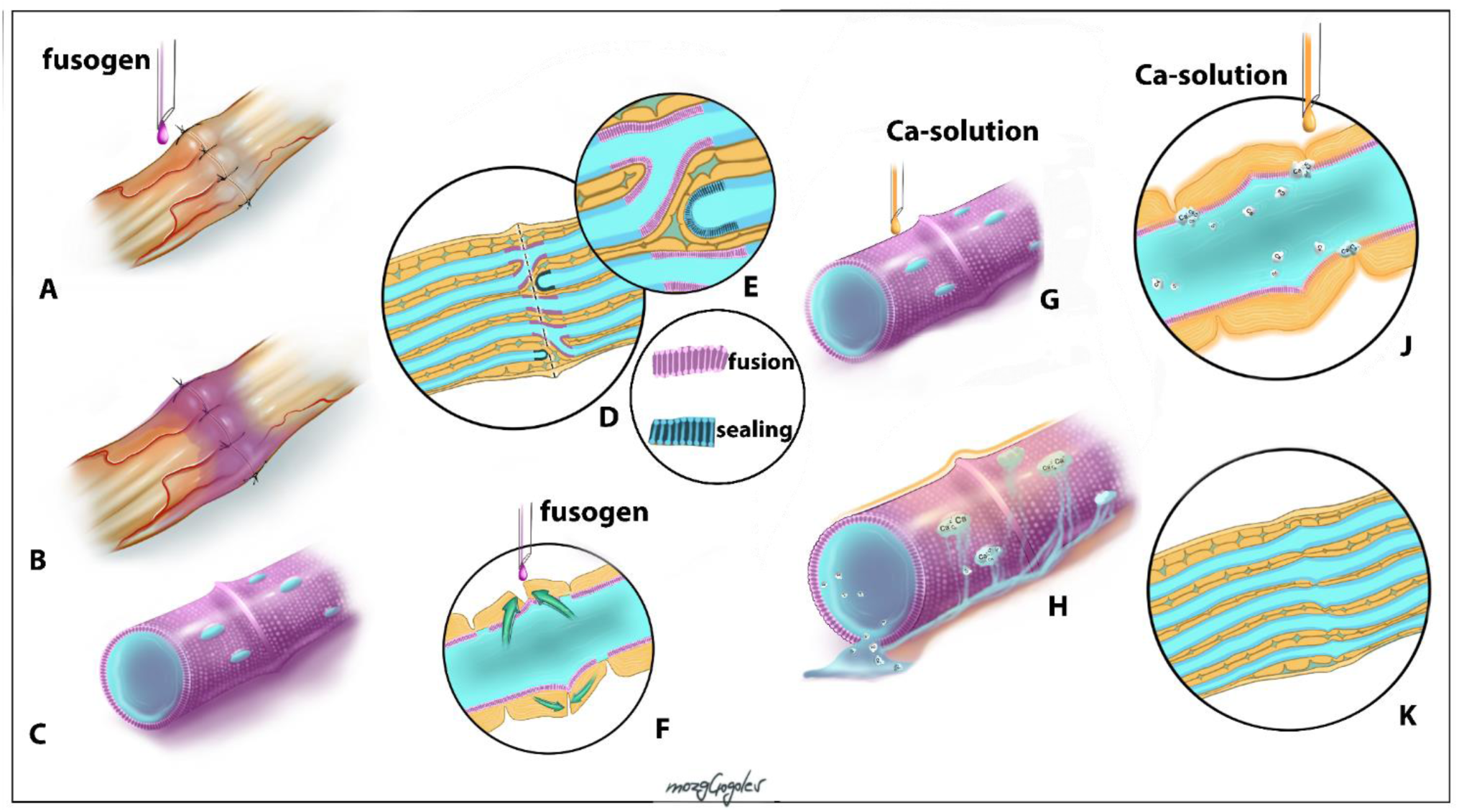
Axonal fusion after neurorrhaphy. **A** - Fusogen (polyethylene glycol, according to the Bittner protocol) is applied to the prepared and sutured nerve; **B** - Fusogen spreads evenly over the suture site, penetrating the intercellular space and remaining in place for approximately 1 minute; **C** - View of an individual axon after fusogen treatment. Two axons have fused, but there are unsealed holes; **D** - Schematic representation of the fusogen’s action area at the nerve end junction. Chaotic axon fusion is shown in purple. Sealing of the axon end that has not fused is shown in turquoise; **E** - Enlarged area of membrane fusion/sealing; **F** - Schematic diagram of the fusion using a single axon as an example. Green arrows indicate the mutual attraction of membranes. **G** - Treatment of the fusion area with an isotonic calcium-containing solution using a single axon as an example; **H** - after treatment with a calcium-containing solution, calcium-bound vesicles are presumably formed, sealing the axolemmal holes that arose after axon injury and sealing the fusion site; **J** - schematic micrograph of a single axon after treatment with a calcium solution; **K** - final appearance of the nerve completely subjected to the fusion protocol. Scheme.

After fusogen application, the nerve is irrigated with a calcium-containing solution, which induces vesicle formation and the final sealing of any axolemmal damage following PEG-mediated fusion of the open axonal ends (Bittner et al., 2016). Emphasis is subsequently placed on washing away any residual PEG after fusion induction (Figure 13 G-K).

It is evident that axonal ends not in close contact with each other do not undergo fusion. Instead, membrane changes lead to axon closure and sealing, resulting in the formation of a stump. This distinction is crucial for successful axonal fusion as defined by the Bittner protocol. Thus, given this sealing effect of PEG, merely irrigating the surface of nerve or spinal cord segments with fusogens should theoretically lead to the formation of non-functional axonal stumps.

Importantly, there are conflicting reports regarding the use of a calcium-free solution. For instance, Shi and Borgens (1999) found no statistically significant difference in outcomes when the sciatic nerve fusion protocol was applied to rats with or without a calcium-free solution (Shi and Borgens, 1999). This suggests that the injury does not necessarily have to be inflicted in a Ca^2+^-free environment.

Furthermore, a separate comparative study showed that the PEG fusion protocol using methylene blue yielded better results compared to the protocol where it was omitted (Bittner et al., 2012).

G. D. Bittner himself only applied his methodology to the spinal cord in a contusion injury setting, obtaining a moderate positive result in the delayed phase (a 3–4 point improvement on the BBB scale after 5 weeks); however, no significant difference relative to the control group were detected during the first week post-injury (Control 3.17±1.25 SEM, PEG-group 2.83±0.91 SEM) (Bittner et al., 2015).

Despite the documented success of the Bittner protocol when applied to sharply transected peripheral nerves, most fusogen studies on the spinal cord do not incorporate his methodology, disregarding his successes in peripheral nerve surgery (Shi and Borgens, 1999; Ren X. et al., 2019; Ren X. et al., 2022a; Lebenstein-Gumovski et al., 2023c; Shen T. et al., 2024; Zhang et al., 2024b).

Indeed, a number of authors reported first irrigating the stumps with PEG and then approximating and coapting the segments (Ren S. et al., 2017; Kim C-Y. et al., 2017; Liu et al., 2018; Ren S. et al., 2019; Zhang et al., 2024b), thereby omitting the preliminary preparation of axons with solutions of varying ionic composition and osmolarity. Such application of fusogens should presumably seal the axonal ends (Figure 14). Nevertheless, these authors report a positive effect, including the recovery of motor function in animals with complete spinal cord transection.

**Figure 14.**
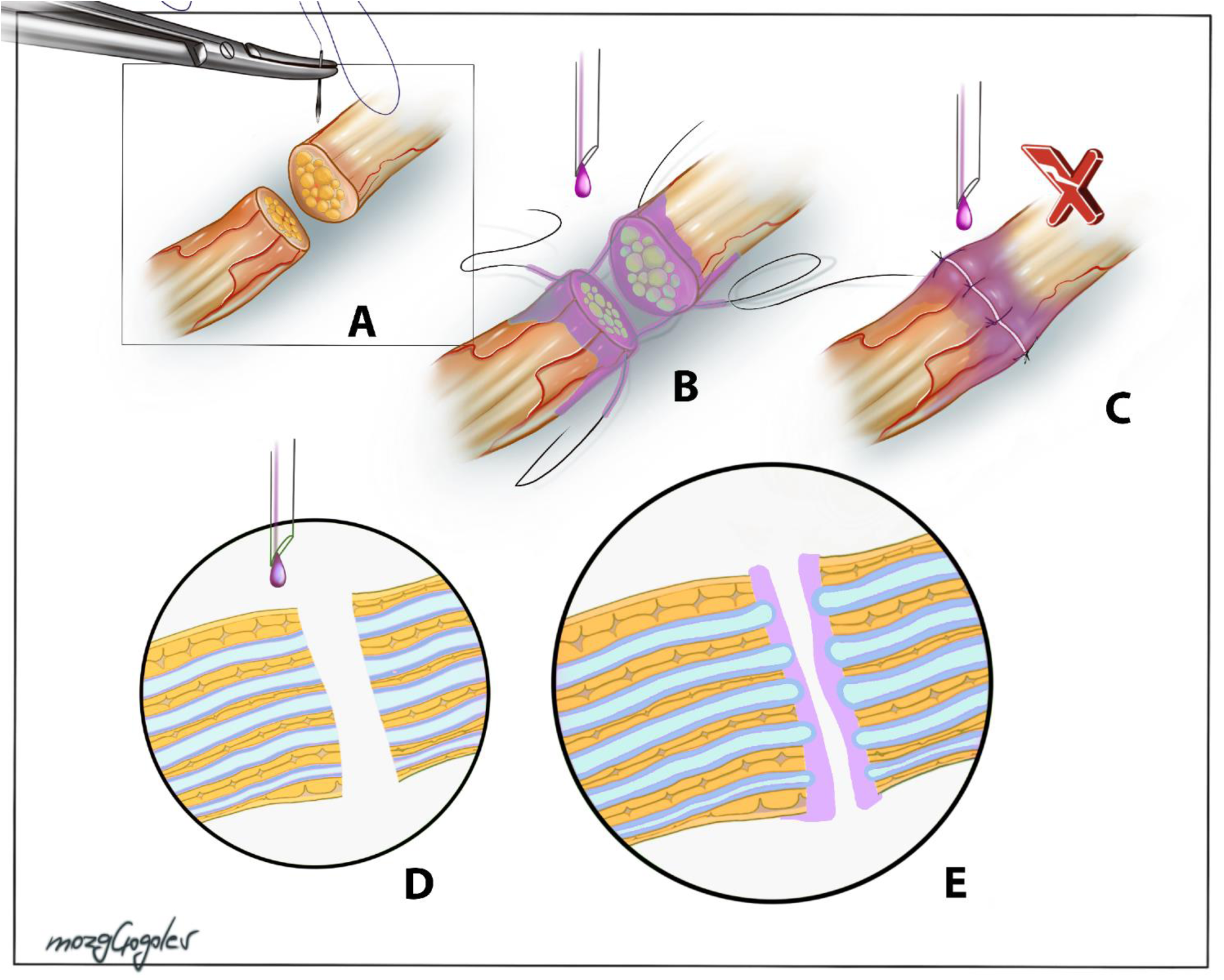
Methodological errors in performing fusogenous fusion. After excision of the injured nerve ends, often without the use of additional solutions **(A)**, fusogen (usually polyethylene glycol) is applied, and alignment and suturing begin **(B, D)**. In some cases, polyethylene glycol is infused before tightening the final sutures and a large diastasis **(C)**. Both options are methodologically flawed, as they lead to PEG sealing of all axons in the transection area, and PEG is not a fusion inducer but a medium that fills the diastasis and isolates the nerve ends **(E)**.

Moreover, some studies reported achieving axonal regeneration by intentionally creating a 1-mm gap between nerve segments filled with PEG (Zhao X. et al., 2019). This work, which applied PEG-600 after dissecting the sciatic nerve in rats, explicitly states that the Bittner protocol is unnecessary but does not explain the mechanism behind the observed effect.

Research on the ability of PEG to influence membrane fusion spanning over 50 years specifically defines the primary prerequisite for its occurrence: the membranes of fusing cells must be in moderately close contact. The step-by-step Bittner protocol logically accounts for this condition. Consequently, creating gaps between the connected stumps of nerves or the spinal cord cannot facilitate PEG-mediated fusion. Thus, if studies have yielded positive results under these conditions, the mechanism underlying the effect of PEG application in this case must be different.

### 7.2 The Dual Role of Calcium Ions

Although calcium ions are known triggers of cellular apoptosis, emerging evidence indicates that their role in axonal fusion is more nuanced. To elucidate the specific function of divalent metal ions in membrane fusion, numerous *in vitro* studies have employed synthetic lipid vesicles. These investigations demonstrate that divalent cations are pivotal for fusion, primarily through their ability to bind to anionic phospholipids such as phosphatidylserine, phosphatidylglycerol, and cardiolipin (Papahadjopoulos et al., 1990). A principal function of these ions is to screen the negative surface charge of the bilayer, thereby reducing electrostatic repulsion and enabling close membrane apposition. However, charge screening alone is insufficient, as evidenced by a marked difference in the fusogenic capacity of Mg^2+^ and Ca^2+^. While Mg^2+^ induces vesicle aggregation, Ca^2+^ is uniquely capable of triggering full fusion (Leventis et al., 1986).

This discrepancy is hypothesized to arise from differences in dehydration potency. Calcium ions exhibit a higher affinity for charged lipid headgroups and a lower degree of coordination with water molecules. The consequent destabilization of the lipid-water interface facilitates close intermembrane contact and promotes fusion (Wilschut et al., 1981). An alternative hypothesis suggests that calcium binding generates destabilizing lateral tension within the membrane (Chanturiya et al., 2000). Regardless of the precise mechanistic pathway, the initial interaction is unequivocally electrostatic, as zwitterionic lipids remain unresponsive (Papahadjopoulos et al., 1974; Pannuzzo et al., 2014). This specificity is further highlighted by the fact that vesicles enriched with certain anionic lipids (e.g., phosphatidylserine) readily fuse in the presence of calcium, while others do not, even if negatively charged (Allolio and Harries, 2021).

Beyond *in vitro* systems, calcium plays a critical role in neuronal survival and regeneration post-injury *in vivo* (Giaume et al., 2013). Axolemmal rupture triggers a massive Ca^2+^ influx, which initiates a self-sealing process, resulting in the compartmentalization of the axon into proximal and distal segments relative to the injury site. This sequestration leads to Wallerian degeneration of the distal segment while preserving the integrity of the proximal axoplasm and the neuronal soma. The Bittner protocol ingeniously co-opts this calcium-dependent sealing mechanism to enhance the efficacy of PEG-mediated fusion. Following the partial PEG-induced restoration of plasmalemmal integrity, the application of a calcium-rich saline solution promotes the accumulation and fusion of vesicles to seal any remaining membrane defects (Bamba et al., 2016a). Thus, calcium is indispensable for PEG-mediated axonal fusion. The sealing of axolemmal damage occurs via a calcium-dependent accrual of membranous structures that form a plug by integrating with the adjacent intact membrane. Paradoxically, in injured neural tissue, this same calcium-dependent system can also cap severed axonal ends, thus preventing their fusion with neighboring fragments (Sexton et al., 2012; Riley et al., 2015; Sexton et al., 2015).

This dual role may be further complicated by the ability of PEG to form coordination complexes with metal ions (Chen J. et al., 2005), including mercury (Rogers et al., 1993), bismuth (Rogers et al., 1992b), lead (Rogers et al., 1996), lanthanides (Rogers et al., 1997), uranium (Rogers et al., 1992a), and strontium (Rogers et al., 1994). PEG-8000 also forms stable complexes with Fe^3+^, Co^2+^, and Ni^2+^ (Sari et al., 2006).

Relevant to its neural application, PEG-4000 forms a stable complex with Ca^2+^, with a stoichiometry of 4.4±0.1(CH_2_CH_2_O):1(CaCl_2_):4.5± 0.5 (H_2_O) and a melting point of 118±2°C (Horikoshi et al., 1990). Similarly, PEG-2000 and PEG-8000 form various Ca^2+^ complexes in alcoholic solutions, which are stable up to 120°C (Sun et al., 2018). In aqueous solutions, PEGs form a PEG:CaCl_2_:6(H_2_O) complex, as characterized by MALDI-TOF mass spectrometry, UV-Vis and FTIR spectroscopy, elemental analysis, and thermogravimetry (Mwelase and Bariyanga, 2002). These studies indicate that calcium ions are coordinated not directly with the PEG ether oxygens, but rather through intermediary water molecules. Upon heating, this complex loses its water of crystallization, yielding a stable dehydrated structure.

Critically, emerging evidence suggests that a PEG-metal ion complex may be the active agent enabling membrane fusion. The treatment of erythrocytes with dehydrating concentrations of commercial PEG-6000 induces hemifusion (Ahkong et al., 1994). However, this fusogenic capacity is abolished when metal ions are removed from PEG solutions using the chelating resin Chelex-100. Furthermore, the cytoplasmic fusion typically observed following the rehydration of dehydrated erythrocytes also fails to occur if both the PEG-6000 and the rehydration buffer are first treated with Chelex-100. Fusogenic activity is restored by replenishing the chelated PEG with La^3+^or Al^3+^ ions at concentrations as low as 10 µM. Other metals, including Ca^2+^, are also effective, though requiring a higher concentration of 100 µM. Significantly, even the trace amounts of calcium found in distilled water are sufficient to support erythrocyte fusion. The stability of the putative PEG-Ca^2+^ complex is underscored by the observation that adding 5 mM of the chelator EGTA to the PEG solution does not inhibit fusion. This is consistent with reports that PEG remains effective under clinically relevant conditions of low Ca^2+^ and reduced temperature in both cultured cells and *ex vivo* spinal cord models (Nehrt et al., 2010).

These findings support a model wherein the PEG-Ca^2+^ complex facilitates fusion by binding to negatively charged lipid headgroups, dehydrating the interfacial water layer, and reducing the net negative charge of the membrane. The fusogenic effect manifests at threshold concentrations above 10 µM for Al^3+^ (0.27 mg/L) and 100 µM (4 mg/L) for Ca^2+^.

Regarding the role of calcium ions in apoptosis, it is noteworthy that irrigation with calcium-free solutions and the use of calcium chelators such as BAPTA (Kang et al., 2021) and EGTA (Petrov et al., 2022) were shown to reduce tissue edema and apoptosis, resulting in improved functional outcomes. Notably, as early as 1984, L. de Medinaceli and A.C. Church proposed the local use of chlorpromazine during neurorrhaphy to inhibit calmodulin, a Ca^2+^-binding protein (de Medinaceli and Church, 1984), which similarly diminished edema and nerve degeneration.

High tissue concentrations of calcium ions undoubtedly negatively impact damaged nervous tissue, and their removal from the injury area—whether via either irrigation with calcium-free solution or the application of calcium chelators or calmodulin inhibitors—reduces apoptosis and inhibits Wallerian degeneration. Meanwhile, PEG-mediated fusion is likely impossible without the formation of a PEG-Ca^2+^ complex. We therefore conclude that this complex can form even with the trace amounts of calcium ions that remain in the surrounding tissues after the application of chelators or Ca^2+^-free solutions, which is sufficient to achieve fusion. We also note the necessity of using calcium chelators to reduce secondary damage in nervous tissue. The more the process of excitotoxicity is inhibited, the greater the number of viable axons that can be fused using fusogens.

### 7.3 Timing of Fusogen Application

Available data on fusogen efficacy underscore the importance of early intervention for improving long-term functional recovery after SCI, given the ongoing cell death due to apoptosis and oxidative stress post-injury (Shi, 2013). For instance, a reduction in the calcium load in synaptosomes is observed within the first 1–4 hours after injury and PEG application, but this effect virtually disappears after 24–72 hours. However, even at later time points, animals treated with PEG exhibit sustained suppression of oxidative stress, despite their intracellular calcium levels remaining similar to those in untreated animals. This suggests that the dominant effect early on is the restoration of plasma membranes, while at later stages, mitochondrial stabilization prevails (Luo et al., 2004).

Evidence supports that PEG retains efficacy even with delayed administration. For instance, partial recovery of conduction and anatomical integrity of axons is observed even when PEG is applied 6–8 hours after contusion injury. Similar results have been obtained with both local application of PEG to the spinal cord and subcutaneous administration (Borgens and Bohnert, 2001; Borgens et al., 2002; Shi, 2013).

Furthermore, axonal fusion remains achievable even after 4 days post-nerve transection under hypothermic conditions, as shown by Marzullo et al. (2002).

### 7.4 Type of Fusogen and Concentration

Despite the widespread use of PEG-600 in SCI models and PEG-3350 in PNI models, a clear understanding of the advantages conferred by specific fusogen types and their MWs remains elusive. Ye et al. (2016) showed that while PEG-1500 outperformed PEG-4000, both led to spinal cord recovery. Additionally, Kouhzaei et al., (2013b) demonstrated that the lower MW of PEG, the higher the ultimate recovery of spinal cord evoked potentials (i.e., PEG-200: 49.5% and PEG-2000: 16.3%). Low-molecular-weight PEGs induced a higher membrane sealing rate (77.8 ± 3.5 for 20% *w*/*w* PEG-400 *vs*. 32.1 ± 6.9 for 20% *w*/*w* PEG-2000). The best results were obtained with PEG-400 under hyperthermic conditions (40°C). Notably, PEG-1000 and PEG-2000 did not show significant effects at high concentrations (>50%).

The debate persists regarding whether PEG should be diluted by mass in distilled water, saline, or phosphate-buffered saline. Given the physical properties of high-molecular-weight PEG (solid form at MWs>1000 Da) (D’souza and Shegokar, 2016), the necessity for mass dilution is easily explained, as a solid substance will exhibit negligible fusogenic capacity due to limited interaction with cell membranes.

The need for mass dilution when using intravenous administration can also be explained by the requirement to achieve a sufficiently fluid fusogen form to avoid embolisms, thrombosis, and intravascular hemolysis.

Furthermore, most studies have exclusively investigated PEG, with other fusogens being used extremely rarely, despite their potential to yield better results in some studies (Ren X. and Canavero, 2020).

Combinations of chitosan and PEG, prepared either as homogeneous mixtures containing a photosensitive chitosan form or via covalent conjugation, showed good results in experiments on rabbits and pigs, resulting in rapid regression of neurological deficits (Lebenstein-Gumovski et al., 2021; Lebenstein-Gumovski et al., 2022; Lebenstein-Gumovski et al., 2023a; Lebenstein-Gumovski et al., 2023b; Lebenstein-Gumovski et al., 2023c; Lebenstein-Gumovski et al., 2023d). Good results were also achieved with a chitosan-PEG combination in a rat model of peripheral nerve repair (Amoozgar et al., 2012).

### 7.5 Method of Fusogen Administration and Systemic Application

Successful axonal repair requires the implementation of axonal-fusion technology aimed at direct membrane sealing of nerve tissue segments, which can only be achieved through intraoperative local application of the fusogen directly to the injury site. This explains why other administration methods—subcutaneous, intraperitoneal, subdural, intravenous—individually show less pronounced effects in neurological functional recovery.

Baptiste et al. (2009) obtained a moderate positive effect from the intravenous administration of a 30% PEG-2000 solution in a rat spinal cord crush injury at the C8 level. The author noted that this moderate degree of motor function recovery suggests that the neuroprotective capacity of this compound may be inadequate for clinical application as a monotherapy (Baptiste et al., 2009).

In a comparative study on rabbits, PEG-chitosan was applied locally during surgery and supplemented with 5 mL of 20% PEG-400 IV daily for 40 days (Lebenstein-Gumovski et al., 2023d). A similar regimen was used in pigs, where Neuro-PEG was administered intraoperatively at the lesion site, accompanied by intravenous injections of 50 mL of 25% PEG-600 IV twice daily for 7 days (Lebenstein-Gumovski et al., 2023b). Both studies reported superior outcomes compared to those achieved with local fusogen application alone in other research.

Given that several authors have achieved positive effects using systemic PEG application (intraperitoneal or intravenous), we propose that this method should be used in combination with local PEG application. Notably, following subcutaneous administration, FITC-labeled PEG accumulated with in the area of hemorrhagic contusion but failed to penetrate traumatically unchanged areas of the spinal cord (Borgens and Bohnert, 2001). This selective accumulation of PEG at the injury site is likely attributable to damage to the blood-spinal cord barrier at the contusion site. Nevertheless, such delivery of a substance with strong neuroprotective and fusogenic effects could also be used to stabilize the injury and the excitotoxic cascade of secondary damage development in the acute phase of trauma. Thus, we suggest that PEG infusions could be beneficial for all patients with spinal trauma, starting at the prehospital stage.

Since the intraoperative using of fusogens is brief and repair processes can last from seconds to hours, maintaining constant low doses of fusogens at the injury site via the microcirculation requires intravenous infusions that are continued into the postoperative period.

It has been shown that the prolonged intravenous use of subtoxic doses in dogs did not cause negative systemic effects, while repeated intravenous injections of high-dose PEG-400 elicited only minimal toxicity, with any occurring changes being reversible (Li et al., 2011). Another study demonstrated that intravenous administration of PEG to dogs was clinically safe and had no acute impact on hematological parameters (Ruble et al., 2006).

### 7.6 Selection of the Optimal Model for Preclinical Studies

Although laboratory rats remain the predominant model in both spinal cord and PNI research, we argue that they are suboptimal for translational spinal cord trauma studies. Based on anatomical and physiological considerations, large animal models—such as dogs, pigs, and non-human primates—offer superior translational fidelity.

However, practical constraints, including high cost, limited availability, specialized housing needs, and complex bioethical considerations, render non-human primates and dogs less accessible. Accordingly, the domestic pig emerges as the most viable and optimal translational model. The anatomical and physiological organization of its motor system, including the trajectory of the corticospinal tract, closely mirrors that of humans, making it an exceptionally relevant model for SCI research (Leonard et al., 2017; Schomberg et al., 2017; Toossi et al., 2021).

### 7.7 Application of Fusogens in Humans

Clinical data on fusogen application for human SCI or PNI remains limited. To date, only two reports have documented PEG-mediated peripheral nerve fusion in humans. One, by Bamba et al. (2016b), details a clinical case where two patients underwent repair of four transected digital nerves using the Bittner protocol. This involved the resection of sclerotic segments, stump freshening, irrigation with calcium-free solution, microsurgical suture, methylene blue application, and subsequent PEG treatment, resulting in significant sensory recovery 1, 4, and 8 weeks post-operatively.

A second study by Lopez et al. (2022) also reported PEG application in two patients. One patient with a sensory digital nerve injury regained fingertip sensation within 1–2 hours post-surgery, achieving near-normal two-point discrimination by day 80. The other patient, who presented with median nerve rupture, showed improved sensation, though motor function outcomes were ambiguous.

Additional cases involve PEG-600 administration combined with spinal cord segment transposition or sural nerve grafting between stumps in chronic ASIA A SCI patients (Ren X. et al., 2022a; Ren X. et al., 2022b). The best outcome in these experiments was recovery to ASIA C (*n*=1).

### 7.8 The Debridement Theory in Spinal Cord Injury: A Primary Direction for Clinical Fusogen Implementation

A pivotal determinant for successful neural repair and neuroregeneration is the absence of a hostile injury microenvironment. Standard contusion injury results in conditions profoundly inhibitory to repair—excitotoxicity, electrolyte imbalance, ischemia, edema, immune activation, necrosis, and cellular infiltration. Later stages are characterized by fibrotic-cystic-gliotic transformation and scar formation, which actively impede axonal growth and render reparative therapies ineffective.

Seminal work by L.W. Freeman in a canine model in the 1960s demonstrated that extensive debridement of a ∼2 cm crush injury, followed by resection to healthy tissue, vertebrectomy, precise re-anastomosis of fresh cord ends, dural closure, spondylodesis, and subdural trypsin infusion, resulted in functional recovery. Months later, the dogs could ambulate short distances and stand on their hind limbs. Histology revealed mutual axonal sprouting between proximal and distal segments (Freeman, 1954, 1962, 1963). Crucially, intraoperative transection with an ultra-thin blade applies minimal force (<10 N) compared to traumatic contusion forces (∼26,000 N) (Sledge et al., 2013), thereby minimizing secondary injury and fostering an environment conducive to repair.

Yoshida et al. (2013) further showed that a clean transection generates negligible edema and avoids subsequent cyst/scar formation, whereas blunt injury precipitates edema, fibro-gliotic scarring, and cystic changes. However, delayed approximation (>10–20 min) led to stump edema, as transected axons remain stable for only this brief window before undergoing Wallerian degeneration (Yoshida et al., 2013). Thus, both cut precision and rapid re-anastomosis are critical.

Research over the past 15 years has generated a foundational framework for optimal SCI repair, emphasizing key steps: a clean transection, tight and rapid approximation, the creation of a pro-regenerative milieu, the use of membrane-active reparative agents, and staged postoperative rehabilitation.

For acute SCI, our team has proposed a “Debridement Theory” (Lebenstein-Gumovski and Grin, 2024), which posits that timely excision of the primary injury zone can interrupt the mutually reinforcing cascade of excitotoxicity and secondary damage. In practice, this entails transecting the spinal cord to the level of viable tissue, thereby halting necrotic progression, and establishing a favorable substrate for recovery—a strategy feasible only with axonal fusion technology (Lebenstein-Gumovski and Grin, 2024).

The concept of spinal cord segment allotransplantation has recently emerged (Shen H. et al., 2019). While matching individual tracts poses practical challenges, neuroplasticity may mitigate this limitation (Guérout, 2021). Axonal funiculi maintain structural and functional integrity for ∼20 minutes, permitting functional adaptation without exact axon-to-axon matching. Promisingly, allotransplantation of a vascularized spinal cord segment in dogs using PEG-600 and tacrolimus has yielded encouraging results (Zhang et al., 2024a).

In contrast, replacing chronic scar tissue is likely futile. Chronic injury entails irreversible white matter degeneration, distal axon loss, and extensive cystic-fibrotic transformation, though some gray matter may persist.

The efficacy of fusogens in SCI may be related to their action on short axonal tracts, such as the cortico-truncoreticulo-propriospinal pathways (Canavero, 2015; Canavero et al., 2016; Canavero and Ren X. P., 2016). Even after corticospinal tract injury at C4–C5, the recovery of precise motor function can occur within 1–3 months, likely relay via rostral interneurons such as cervical propriospinal neurons (Nishimura and Isa, 2012; Tohyama et al., 2017). In primates, direct cortico-motoneuronal connections underpin voluntary movement, yet neural plasticity enables recovery after corticospinal tract damage at C4/C5 through circuit reorganization across the central nervous system (Nishimura and Isa, 2012; Tohyama et al., 2017).

We hypothesize that fusogen-mediated repair after complete transection involves not only white matter axonal fusion but also non-selective gray matter fusion. The potent neuroprotection offered by PEG and chitosan may shield gray matter from excitotoxicity and vascular insult, thus preserving neuronal pools. Minimizing secondary damage via sharp transection and fusogen application could enable long-term functional recovery through short-neuron relay mechanisms.

This mechanism might also operate in chronic injury models where residual gray matter persists, explaining the partial success of neural transplantation in chronic SCI (Ren X. et al., 2022a; Nourbakhsh et al., 2024), albeit with inferior outcomes compared with those achieved in acute injury.

We contend that PEG or chitosan monotherapy is insufficient without a holistic understanding of their mechanisms and a conducive environment for fusion. Successful clinical translation necessitates a multifaceted strategy. We pioneered an integrated fusogenic therapy approach (Lebenstein-Gumovski et al., 2023b) that incorporates local hypothermia, a novel PEG-chitosan conjugate, electrical stimulation, and comprehensive rehabilitation. We also developed a technique for limited spinal cord resection with wedge osteotomy or Smith-Petersen traction spondylotomy, coupled with combined fusogen delivery (intraoperative local + prolonged intravenous infusion) (Lebenstein-Gumovski et al., 2021; Lebenstein-Gumovski et al., 2022). In one study, rabbits with complete transection receiving local fusogen and IV PEG infusion showed superior neurological recovery relative to controls not receiving infusion (Lebenstein-Gumovski et al., 2023d).

### 7.9 Conclusion

Fusogens have been known for over 50 years, however recent discoveries are opening up a new niche for them in neuroregeneration and neurosurgery (Figure 15; additional information in Note 1). The application of fusogens to enable accelerated morphological and functional recovery represents a transformative advancement in neurosurgery for treating spinal cord and peripheral nerve injuries. This review and analysis of results from diverse animal and human studies demonstrates the high efficacy of PEG, chitosan, their combinations, and derivatives as agents that induce immediate membrane fusion, culminating in the functional restoration of neural tissue. Future research into these technologies must focus on optimizing the most effective therapeutic combinations, parameters, and delivery methods, while also driving the development and implementation of novel clinical pathways and refined treatment protocols, including a critical re-evaluation of intervention timeframes.

**Figure 15.**
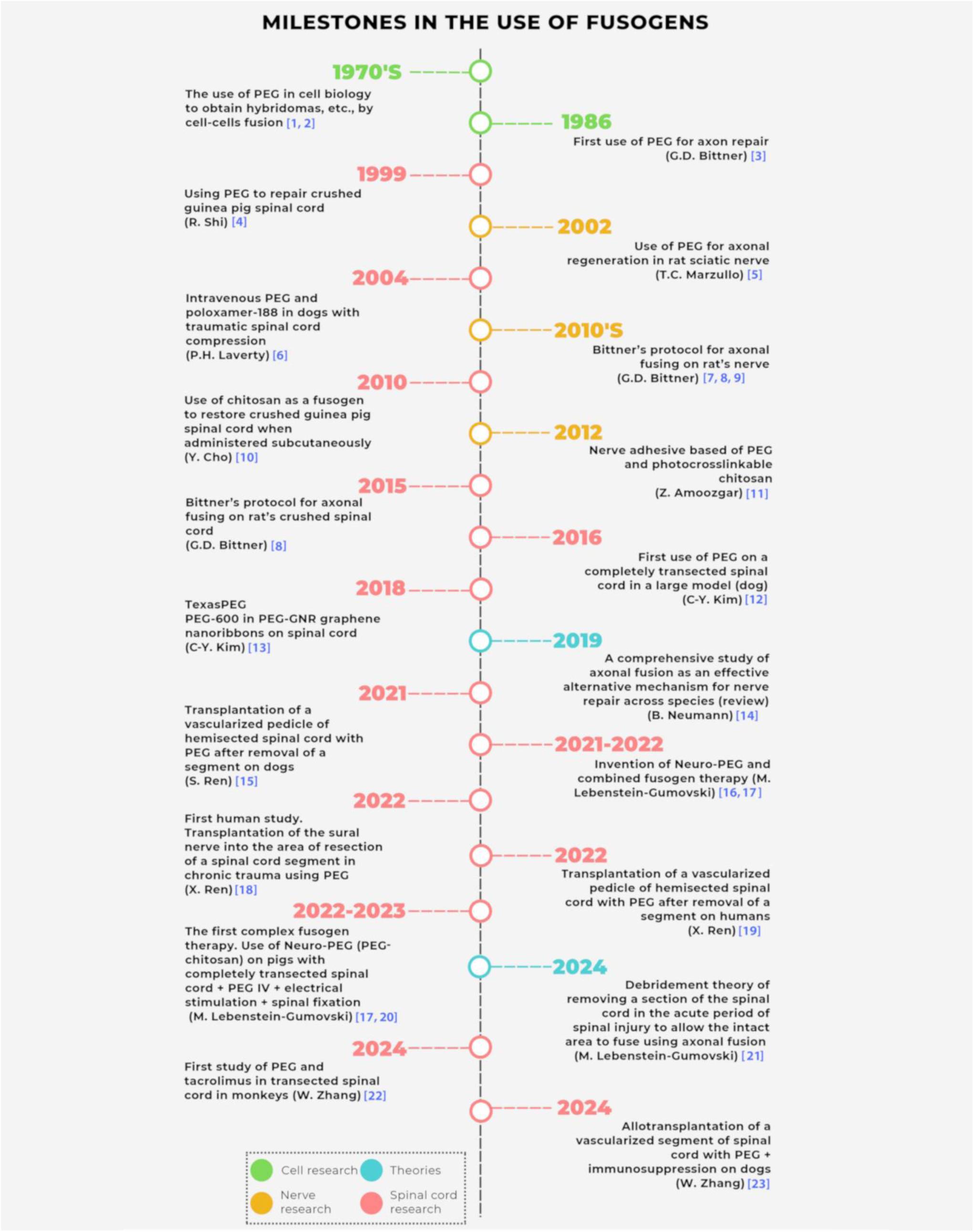
Key milestones in the use of fusogens.

## 8 Conflict of Interest

The authors declare that the research was conducted in the absence of any commercial or financial relationships that could be construed as a potential conflict of interest.

## 9 Author Contributions

ML-G: Conceptualization, Supervision, Formal analysis, Writing - original draft, Writing - review & editing. YR: Formal analysis. DK: Writing - original draft. TR: Data curation, Writing - original draft. SC: Writing - review & editing. AZ: Writing - original draft. AT: Writing - review & editing. AG: Writing - review & editing.

## 10 Funding

The authors declare that no financial support was received for the research, authorship, and/or publication of this article.

## 11 Acknowledgments

We thank Andrey Panferov, Alina Nurislamova for management and language support of this article.

Additional studies information, indicated by numbers, is provided in Note 1.

**Note 1.** Description of the studies in Figure 15.

## Authors Information

M.V. Lebenstein-Gumovski – MD, PhD, Senior Scientific Officer at Sklifosovsky Clinical and Research Institute for Emergency Medicine, Neurosurgery Department: conceptualization, supervision, formal analysis, writing (original draft), writing (review & editing).

Y.V. Romanenko – Junior Researcher at Sklifosovsky Clinical and Research Institute for Emergency Medicine, Neurosurgery Department: formal analysis.

D.A. Kovalev – PhD, Head of Department at Stavropol Anti-Plaque Research Institute, laboratory of biochemistry: writing (original draft).

T.S.-M. Rasueva – Junior Researcher at Sklifosovsky Clinical and Research Institute for Emergency Medicine, Neurosurgery Department: data curation, writing (original draft).

S. Canavero – MD, PhD, Head of GEMINI international group: writing (review & editing).

A.M. Zhirov - Researcher of Department at Stavropol Anti-Plaque Research Institute, laboratory of biochemistry: writing (original draft).

A. Talypov - MD, Doctor of Medical Sciences, Professor, Senior Scientific Officer at Sklifosovsky Clinical and Research Institute for Emergency Medicine, Neurosurgery Department: writing (review & editing).

A.A. Grin’ – MD, Doctor of Medical Sciences, Professor, Corresponding Member of the Russian Academy of Sciences, Chief Neurosurgeon of Moscow, Head of the Neurosurgery Clinic at Sklifosovsky Clinical and Research Institute for Emergency Medicine, Neurosurgery Department, Honored Doctor of the Russian Federation: writing (review & editing).

## Notes

### Competing Interest Statement

The authors have declared no competing interest.

